# Pervasive neurovascular dysfunction in the ventromedial prefrontal cortex of female depressed suicides with a history of childhood abuse

**DOI:** 10.1101/2024.07.29.605502

**Authors:** Marina Wakid, Daniel Almeida, Ryan Denniston, Anjali Chawla, Zahia Aouabed, Maria Antonietta Davoli, Kristin Ellerbeck, Reza Rahimian, Volodymyr Yerko, Elena Leonova-Erko, Gustavo Turecki, Naguib Mechawar

**Affiliations:** McGill Group for Suicide Studies, Douglas Research Centre, Montréal, Quebec, Canada; Integrated Program in Neuroscience, McGill University, Montréal, Quebec, Canada; Department of Psychiatry, McGill University, Montréal, Quebec, Canada

## Abstract

Exposure to early life adversity (ELA) poses a significant global public health concern, with profound pathophysiological implications for affected individuals. Studies suggest that ELA contributes to endothelial dysfunction, bringing into question the functional integrity of the neurovascular unit in brain regions vulnerable to chronic stress. Despite the importance of the neurovasculature in maintaining normal brain physiology, human neurovascular cells remain poorly characterized, particularly with regard to their contributory role in ELA-associated pathophysiologies. In this study, we present the first comprehensive transcriptomic analysis of intact microvessels isolated from postmortem ventromedial prefrontal cortex samples from adult healthy controls (CTRL) and matched depressed suicides with histories of ELA. Our findings point to substantive differences between men and women, with the latter exhibiting widespread gene expression changes at the neurovascular unit, including the key vascular nodal regulators *KLF2* and *KLF4*, alongside a broad downregulation of immune-related pathways. These results suggest that the neurovascular unit plays a larger role in the neurobiological consequences of ELA in human females.

## Introduction

Exposure to adverse childhood experiences (ACEs), defined as physical, sexual, and/or emotional abuse or neglect, is a persisting global public health concern. In developed countries, 44% of children have been subjected to ACEs, while the percentage rises to 59% in developing countries (1). Early life adversity (ELA) can lead to profound disturbances in psychological and physical trajectories that, in turn, strongly correlate with increased lifetime risk of negative health outcomes. Notably, compelling evidence supports ACE-induced vascular endothelial dysfunction, characterized by increased arterial stiffness, higher peripheral vascular resistance, and reduced endothelial function (2, 3). Such dysfunction primes for the development of cardiovascular disease later in life (2, 4–10). The observed correlation between ELA and cardiovascular disease is physiologically sound. Indeed, the prefrontal cortex, an emotion-modulating (11) core component of a broader network of forebrain systems, mediates stress-evoked changes in cardiovascular activity (12, 13); and there is ample evidence demonstrating sustained and pervasive molecular (14–18), cellular (19–22) and functional abnormalities (23–28) of the prefrontal cortex following ELA. Although the blood-brain barrier (BBB) is functionally distinct from the peripheral vasculature and possesses a highly specialized neurovascular unit (NVU) to precisely regulate the influx and efflux between the blood and brain parenchyma (29), the entire blood supply of the brain relies on the dorsal aorta (30). This reliance therefore maintains a structural connection to the peripheral vasculature, raising the question as to whether the NVU is similarly affected by ELA. Critically, ELA is associated with 44.6% of all psychiatric childhood-onset disorders and with 25.9% to 32.0% of adult-onset disorders (31), such as major depressive disorder (MDD) (32–37). Despite this, direct evidence implicating NVU dysfunction in the pathophysiology of ELA relies primarily on rodent models of chronic stress, which have focused on downregulated tight junction protein Claudin5 (Cldn5) (38–40) or on the formation of cerebral microbleeds (41–44). In contrast, studies investigating NVU dysfunction in adult humans with histories of ELA are limited to non-invasive techniques that characterize markers of plasma inflammation (45–50), using these markers to infer the state of the NVU. Critically, the use of animal proxy models cannot completely recapitulate human experience and its effects on human neurobiology (51), and recent breakthroughs have demonstrated that there are numerous species-specific differences between mouse and human neurovasculature (52–54). For this reason, we set out to generate the first transcriptomic dataset derived from intact microvessels isolated from postmortem ventromedial prefrontal cortex (vmPFC) samples from controls and matched depressed suicides with a history of ELA. Isolated microvessels effectively capture the NVU, comprising brain microvascular endothelial cells (BMECs) sealed by tight junction proteins, astrocytic endfeet, and mural cells (55). Here, we combined differential gene expression analysis and network-based approaches to provide an integrative and unbiased characterization of male and female transcriptional profiles of the NVU in humans with histories of ELA.

## Methods

### Human post-mortem brain samples

This study was approved by the Douglas Hospital Research Ethics Board. Brains were donated to the Douglas-Bell Canada Brain Bank (www.douglasbrainbank.ca; Montreal, Canada) following written informed consent from next of kin, in the context of collaboration with the Quebec Coroner’s Office. Phenotypic information was retrieved through standardized psychological autopsies (56). In brief, proxy-based interviews with one or more informants best acquainted with the deceased were supplemented with information from archival material obtained from hospitals, Coroner’s office, and social services. Clinical vignettes were then produced and assessed by a panel of clinicians to generate a diagnosis based on the DSM-IV. Toxicological assessments and medication prescription are also obtained. As described previously (15), characterization of early-life histories was based on adapted Childhood Experience of Care and Abuse interviews assessing experiences of sexual and physical abuse, as well as neglect (57). The severity early-life adversity was assessed based on reports of non-random major physical and/or sexual abuse during childhood (up to 15 years). Only cases with the maximum severity ratings of 1 and 2 were included in this study. Because of this narrow selection criterion, it was not possible to stratify different types of abuse within the sample. Presence of any or suspected neurological/neurodegenerative disorder reported in clinical files constituted an exclusion criterion. Individuals were all Caucasians of French-Canadian descent. Samples from 21 healthy controls (13 males, 8 females) and 24 depressed suicides with a history of severe childhood abuse (17 males, 7 females) were analyzed in this study. Group characteristics are described in Table 1.

**Table 1:** Demographic and sample characteristics of CTRL and ELA cohorts. Numeric values in each cell represent the mean ± SD. *MDD:* major depressive disorder, *PMI:* postmortem interval. The postmortem interval is a metric for the delay between an individual’s death, the collection and processing of the brain.

|  | **CTRL (sexes pooled)** | **ELA (sexes pooled)** |  |  |
| --- | --- | --- | --- | --- |
| n | **21** | **24** |  |  |
| Sex (M/F) | 13/8 | 17/7 |  |  |
| Age (years) | 49.10 ± 18.71 | 42.54 ± 14.70 |  |  |
| Axis 1 diagnosis | 0 | MDD (24) |  |  |
| History of abuse | 0 | 24 |  |  |
| PMI (h) | 40.32 ± 28.21 | 41.51 ± 22.24 |  |  |
| Tissue pH | 6.43 ± 0.31 | 6.58 ± 0.39 |  |  |
|  | **Male CTRL** | **Male ELA** | **Female CTRL** | **Female ELA** |
| n | 13 | 17 | 8 | 7 |
| Age (years) | 41.00 ± 13.64 | 37.88 ± 10.18 | 62.25 ± 19.00 | 53.86 ± 18.48 |
| Axis 1 diagnosis | 0 | MDD (17) | 0 | MDD (7) |
| History of abuse | 0 | 17 | 0 | 7 |
| PMI (h) | 31.18 ± 23.43 | 36.18 ± 20.97 | 55.18 ± 30.42 | 54.46 ± 21.13 |
| Tissue pH | 6.51 ± 0.24 | 6.53 ± 0.38 | 6.29 ± 0.37 | 6.71 ± 0.39 |

### Tissue dissections

Grey matter samples were dissected from the vmPFC by expert brain bank staff on fresh-frozen 0.5 cm-thick coronal sections with the guidance of a human brain atlas (58). vmPFC samples were dissected in sections equivalent to plate 3 (approximately −48 mm from the center of the anterior commissure) of this atlas, corresponding to Brodmann areas 11 and 12. Samples in every form, whether extracted total RNA, RNA sequencing library, or mounted tissue sections, were stored frozen at -80° C until further use.

### Microvessel isolation from tissue microdissection

To capture the complete, multicellular composition of the NVU, microvessels were enriched and isolated using our recently developed protocol (59), which is a method that is gentle enough to dissociate brain tissue while preserving the structural integrity and multicellular composition of microvessels. In brief, 100 mg of microdissected vmPFC grey matter was homogenized in 2 ml of cold homogenization buffer (1M sucrose + 1% BSA dissolved in DEPC-treated water) using the benchtop gentleMACS™ Dissociator (Miltenyi Biotec, Germany) with the rotating paddle set to the Lung 02.01 program. After tissue homogenization, an additional 8 ml of cold homogenization buffer was pipetted into the tube, topping up the homogenate to 10 ml. The homogenate was gently inverted to mix, and then centrifuged at 3200 g for 30 min at 4° C. Following centrifugation, a microvessel-enriched pellet formed at the bottom of the falcon tube, after which the supernatant was carefully vacuum-aspirated (which may include an upper layer of clumped dissociated myelin) without disturbing the microvessel-enriched pellet. To assess RNA quality, RNA integrity number (RIN) was measured for microvessel isolation samples, with an average value of 4.

### Visualization of microvessels from the enriched pellet using immunofluorescence

Immunophenotypic characterization of isolated microvessels was carried out following the resuspension of microvessel-enriched pellets, as per Wakid et al. (2023). Pellets were gently resuspended in 400 μl of cold PBS and 50 µl of the resuspension was pipetted into each well of an 8-well chamber slide (Nunc™ Lab-Tek™ II Chamber Slide™ System; Thermo Scientific™, United States). The chamber slide was left open-faced to incubate in a 37°C oven overnight, allowing microvessels to dry flush to the surface of the slide after the PBS evaporated. After drying, microvessels were fixed by covering them to a depth of 2–3 cm with ice-cold 100% methanol for 15 min on ice or at 4°C. Subsequently, the methanol was aspirated, and the wells were washed with 1X PBS three times for 5 min each. Microvessels were then incubated in blocking buffer (1% BSA + 0.5% Triton X dissolved in PBS) under gentle agitation for 60 min at 4°C. Afterwards, blocking solution was replaced with 500 µl of primary antibody dilution (1:500 in 1% BSA + 2% normal donkey serum + 0.5% Triton X dissolved in PBS) and incubated under gentle agitation overnight at 4°C. Primary antibody dilution was then aspirated and wells were washed in 1X PBS three times for 5 min each. Microvessels were finally incubated in fluorophore-conjugated secondary antibody dilution (1:500 in 1% BSA + 2% normal donkey serum + 0.5% Triton X dissolved in PBS) under gentle agitation for 2 hours at room temperature, protected from all light. Following washes, the media chamber was removed, and the microscope slide carrying microvessels was coverslipped using VECTASHIELD Antifade Mounting Medium with DAPI (Vector Laboratories, Inc., United States). To visualize the expression of canonical NVU markers, primary antibodies targeting key expression markers were used: for BMECs, Laminin (anti-Laminin antibody L9393; Sigma-Aldrich, United States) and *PECAM1* (Anti-PECAM1 antibody (JC70); Santa Cruz Biotechnology Inc., United States); for tight junctions, *CLDN5* (anti-Claudin 5 antibody ab15106; Abcam, United Kingdom); for pericytes, *PDGFR*β (anti-PDGFRβ monoclonal antibody G.290.3; Thermo Fisher Scientific, United States); for smooth muscle cells, Vimentin (anti-Vimentin antibody RV202; Abcam, United Kingdom); and for astrocytic endfeet, *AQP4* (anti-Aquaporin 4 antibody [4/18]; Abcam, United Kingdom). We also employed appropriate fluorophore-conjugated secondary antibodies to thoroughly characterize the collected microvessels.

### Extraction of total RNA from isolated microvessels

For RNA extraction experiments, the microvessel-enriched pellet was gently resuspended in 500 μl of cold PBS and gradually pipetted through a 35 μm Strainer Cap for *FlowTubes*™ (Canada Peptide, Canada) using vacuum-aspiration underneath to encourage filtration. The result is intact microvessels trapped within the strainer mesh, where smaller cellular debris and free-floating nuclei have passed through. Using a flat-ended spatula and point-tip forceps, the strainer mesh was removed from its plastic frame and immediately submerged into 100 μl of RL buffer from the Single Cell RNA Purification Kit (Norgen Biotek Corp., Canada), according to step 1A of the manufacturer’s protocol. The mesh was discarded after transferring 100 μl of fresh 70% ETOH and pipetting 10 times. Total RNA extraction, including on-column DNase digestion, were followed through according to the manufacturer’s instructions. RNA concentration and RIN were quantified using the Agilent TapeStation 2200. RNA samples were then frozen and stored at -80° C until further use.

### Library construction and bulk RNA-sequencing

Microvessel-enriched pellets yielded an average of 10.7 ng/µl of total RNA per sample. Libraries were constructed using the SMARTer Stranded Total RNA-Seq Kit v3-Pico Input Mammalian (Takara Bio Inc., Japan), which features integration of unique molecular identifiers (UMIs) to allow for the distinction between true biological duplicates and PCR duplicates. Libraries were constructed using 10 ng of RNA as input, 2 min of fragmentation at 94°C (ProFlex PCR; Applied Biosystems Corporation, United States), 5 cycles of amplification at PCR1, 12 cycles of amplification at PCR2 and clean-up of final library using NucleoMag NGS Clean-up and Size Select beads (Takara Bio Inc., Japan). Libraries were then quantified at the Genome Quebec Innovation Centre (Montreal, Quebec) using a KAPA Library Quantification kit (Kapa Biosystems, United States), and average fragment size was determined using a LabChip GX (PerkinElmer, United States) instrument. Libraries were sequenced on the NovaSeq 6000 system (Illumina, Inc., United States) using S4 flow cells with 100bp PE sequencing kits.

### Bioinformatic pipeline and analyses of RNA sequencing data

#### UMI extraction, alignment, de-duplication, metrics and generation of count matrix

Bulk RNA sequencing of microvessel libraries yielded an average of ∼64 million reads per library, which were then processed following our in-house bioinformatic pipeline to generate a count matrix with contributions of phenotype (53.3% ELA) and sex (33.3% female). UMI extraction based on fastq files was performed using the module extract from umi_tools (v.1.1.2) (60). Reads were then aligned to the Human Reference Genome (GRCh38) using STAR software (v2.5.4b) (61), with Ensembl v90 as the annotation file and using the parameters: **--twopassMode Basic--outSAMprimaryFlag AllBestScore--outFilterIntronMotifs RemoveNoncanonical--outSAMtype BAM SortedByCoordinate--quantMode TranscriptomeSAM GeneCounts**. Resulting bam files were sorted and indexed using SAMtools (v.1.3.1) (62). Duplicate reads with the same UMI were removed using the dedup module of umi_tools (v.1.1.2) (60). Different metrics, including the fraction of exonic, intronic and intergenic reads were calculated using the CollectRnaSeqMetrics module of Picard (v.1.129; Broad Institute), and the expected counts and transcripts per million (TPMs) were generated using RSEM (v1.3.3; reverse strand mode) (63). RNA sequencing metrics are shown in Supplementary Table 1.

### RNA-sequencing deconvolution

10X Chromium for single-nucleus data from human brain vasculature were accessed from Yang et. al (2022) (54) and used as a reference to perform bioinformatic deconvolution of our bulk RNA-sequencing data. Seurat (64) was used to pre-process raw count expression data, removing genes with less than 3 cells or cells with less than 200 expressed genes. 23054 genes from a total of 23537 and 141468 nuclei from a total of 143793 passed these QC criteria. To generate the Yang signature input for the CIBERSORTx deconvolution tool (65), counts per million (CPM) values were averaged across nuclei of each cell type.

### Differential gene expression analysis and normalization

All subsequent analyses were performed in R. Differential gene expression (DGE) was performed using DESeq2 (1.42.0) (66). Male and female count data were separated, but the same pre-filtering step was applied to both, removing genes with raw counts<25 in 85% of subjects. Data for both males and females were each processed and analysed as follows: **Preprocessing:** key metadata variables Age, pH, PMI, and RIN scaled using the scale function, and Group was converted to a categorical variable using the factor function. **Normalization and PCA Implementation:** Using DESeq2, the count data was normalized via the median of ratios method and stabilized via the vst function, followed by Principal Component Analysis (PCA) to assess potential outliers and the influence of potential covariates on gene expression profiles (Supplementary Fig. 1). **Correlation and Regression Analysis:** Linear regression was employed by the lm function to establish correlations between principal components and covariates, quantifying the relationships between transcriptional variations and covariates. Covariates with correlations p.adj<0.05 were deemed significant and were added to the design formula required by DEseq2. For males, age (PC3: p.adj=0.007; r^2^=0.23) and pH (PC2: p.adj=0.00043; r^2^=0.36) were identified as covariates and, for females, age (PC2: p.adj=0.036; r^2^=0.30) was identified as a covariate. **Differential Expression Model Setup:** An initial DE model is constructed using DESeq2, with relevant covariates included in the design formula. **Normalization and Surrogate Variable Analysis:** counts are normalized using the median of ratios method. The SVA package is then incorporated to identify and include surrogate variables in the model to account for potential unobserved confounders or unmodeled artifacts. A revised DE model is created, integrating the computed surrogate variables alongside the original covariates (Supplementary Fig. 1). **Differential gene expression (DGE):** DGE analysis is conducted using the revised model, and *p*-values were adjusted for multiple testing using the procedure of Benjamini and Hochberg. An adjusted p-value (p.adj)<0.05 and fold change ≥10% (∣log2(FC)∣≥log2(1.1)) were used to designate significant differentially expressed genes.

### PsyGeNET analysis

The list of ELA-associated DEGs from males and females was separately queried in the PsyGeNET database (67) using the psygenet2r R package. The psygenetGene function was used with database=ALL and other parameters set to default to retrieve information about associations between our specific genes and psychiatric diseases. To concisely visualize the output, we generated an evidence index barplot showing, for each psychiatric disorder, the number of gene-disease associations.

### Functional enrichment analysis

Metascape (68) was used to perform an in-depth enrichment analysis of identified DEGs, with the aim of elucidating the functional pathways and cell-type signature significantly associated with identified DEGs. Metascape utilizes Fisher’s exact test to compute p-values and enrichment factors for each ontology category. The input species was set to Homo Sapiens, and a tailored selection of databases was used for annotation, membership, and enrichment analysis (FDR<0.05) was used. The background gene set used for the enrichment analysis comprised genes that passed the filtering threshold during DGE analysis (i.e. those input into DGE analysis). Only pathway terms with a p.adj<0.05, a minimum count of three, and an enrichment factor>1.5 were considered significant.

### Gene set enrichment analysis (GSEA)

For differential expression results, GSEA was performed using the fgsea package (69). A log2(FC)-ranked list of genes was generated, a combined list of gene sets from Reactome pathways, Gene Ontology Molecular Functions (GOMF), KEGG pathways, and WikiPathways was compiled from MSigDB (70), and the following parameters were used for the fgsea function: minSize=15, maxSize=500; and any pathways with Benjamini-Hochberg p.adj<0.05 were considered to be significantly enriched. Finally, we ran collapsedPathways with pval.threshold=0.05 to get the main pathways for each cluster. An additional GSEA was performed for the female data using Brain.GMT (71), which consists of curated brain-related gene sets.

### Rank-rank hypergeometric overlap

A threshold free, rank-rank hypergeometric overlap analysis was performed using the RRHO2 R package (72). Genes were scored using the product of the logFC and the negative log base 10 uncorrected *p*-value taken from DGE analysis in the male and female datasets separately. The scored gene lists were input to RRHO2_initialize function (with method “hyper” and log10.ind “TRUE”) and the results were plotted using the RRHO2_heatmap function.

### Weighted Correlation Network Analysis

Weighted correlation network analysis (WGCNA) (73) was performed to explore the relationship between co-expressed gene modules and exposure to ELA using data specifically from females. This focus on females was chosen based on the results from DGE analysis, which indicated that the female NVU is more greatly affected by ELA compared to males. A pre-filtering step was first applied, removing genes with raw counts<20, followed by normalization via DESeq2’s normalizeCounts function and variance stabilizing transformation. Transformed counts were corrected for the effect of age using limma (74), because it was previously identified as a covariate for females CTRL vs ELA groups. A network was constructed using the following parametres: soft power threshold=14, networkType="signed", minimum module size=100, mergeCutHeight=0.2, maxBlockSize=30000, and deepSplit=3. For each identified module, the eigengene was calculated alongside its correlation with ELA status. Finally, gene significance and module membership were calculated. For modules turquoise and black, the topmost hub gene was identified using the chooseTopHubInEachModule function. Additional hub genes were selected based on their soft connectivity scores, which exceeded a hub-defining threshold. The hub-defining threshold was determined by examining the histogram distribution of soft connectivity scores for all genes within each module.

### Validation of DEGs with fluorescence *in situ* hybridization

DEG candidates were validated using fluorescence *in situ* hybridization. Frozen vmPFC blocks of (primarily) grey matter were prepared from the same CTRL and ELA female subjects that were sequenced, cut into serial 10 µm-thick sections with a cryostat, and mounted on Superfrost charged slides. FISH was performed using Advanced Cell Diagnostics RNAscope® probes and reagents following the manufacturer’s instructions. Sections were first fixed in cold 10% neutral buffered formalin for 15 min, dehydrated by increasing gradient of ethanol baths, and then air dried for 5 min. Hybridization with Hs-KLF2 (408711-C-1; Advanced Cell Diagnostics, United States) and Hs-KLF4 (457461-C-2; Advanced Cell Diagnostics, United States) probes was conducted for 2 hours at 40°C in a humidity-controlled oven. Amplifiers were added using the proprietary AMP reagents and the signal was visualized through probe-specific HRP-based detection by tyramide signal amplification (TSA) with Opal dyes Opal 520 and Opal 570 (FP1487001KT and FP1488001KT; Akoya Biosciences, United States) diluted to 1:700. Immunofluorescent blood vessels were observed in the same sections, post-hybridization. Sections were washed in PBS and then incubated overnight at 4 °C under constant agitation with anti-laminin antibody (anti-Laminin antibody L9393; Sigma-Aldrich, United States) diluted in blocking solution (1:500 in 2% normal donkey serum + 0.2% Triton X dissolved in PBS). After washing in 1X PBS three times for 5 min each, sections were incubated for 2 h at room temperature in fluorophore-conjugated secondary antibody (1:500; Alexa Fluor® 647 AffiniPure™ Donkey Anti-Rabbit IgG, 711-605-152; Jackson ImmunoResearch Laboratories, United States). TrueBlack was used to remove endogenous autofluorescence from lipofuscin and cellular debris. Slides were finally coverslipped in Vectashield mounting medium with DAPI to enable nuclear staining (Vector Laboratories, Inc., United States).

### Imaging and analysis of *in situ* mRNA expression of DEG candidates

Images were acquired with an Olympus VS120 slide scanner and, for each experiment and subject, the whole section was scanned and imaged using a ×20 objective. Exposure parameters were kept consistent between subjects for each set of experiment, where the TRITC channel was designated for a DEG candidate (either *KLF2* or *KLF4*) and the Cy5 channel was designated for laminin. As TSA amplification with Opal dyes yields a high signal-to-noise ratio, parameters were optimized so that autofluorescence from lipofuscin and cellular debris was filtered out. Microvessels were defined by tubular structures with laminin signal (a validated marker that is exclusively expressed by endothelial cells. Image analysis was performed in QuPath (v 0.5.1). Each subject had one section stained with *KLF2* or *KLF4*, as well as DAPI and laminin. To recapitulate the non-discriminatory approach that was used during microvessel isolation, where microvessels of all diameters from every spatial point within the tissue sample is isolated, grey matter microvessels in stained tissue sections were randomly sampled for analysis. This was conducted by carrying out the following steps: grey matter area was outlined using the polygon annotation tool and computing intensity features (namely mean and standard deviation) with a preferred pixel size of 1 µm for the TRITC channel across the entire grey matter area. A grid with spacing of 1000×1000 µm was overlaid onto the section, and each tile covering grey matter was numbered. Using a random number generator, 5 grey matter tiles were selected and annotated using the square annotation tool. Nuclei that did not display laminin staining but were in direct contact with the vessel surface were classified as constituent cells of the NVU. Consequently, these nuclei were included in the analysis as integral components of the NVU. To automate microvessel detection and include those with contacting nuclei, a pixel thresholder for laminin was set, turning these areas into detections within the 5 selected tiles. Similarly, DAPI-stained nuclei were detected. Custom Groovy scripts were used to merge detections of nuclei in contact with microvessels into singular detections and convert these detection objects into annotations. A pixel thresholder was established in the TRITC channel to identify puncta, using a calculated threshold value of the mean+(5xSD). Using the pixel thresholder, the area of annotated microvessels covered by target puncta in the TRITC channel was measured, from which the percent area covered by target puncta within these annotated microvessels was calculated. Percent area covered by target puncta was the preferred measure to reflect RNA expression, as punctate labeling generated by FISH often aggregates into clusters that cannot readily be dissociated into single dots or molecules.

### Statistics

FISH experiments: For each subject, mean percent area for *KLF2/4* expression were calculated, and the following statistical analyses were conducted in R using the stats and car packages: the Shapiro-Wilk test was utilized to assess the normality of the data and Levene’s Test was conducted to compare the variances between groups. For *KLF2*, a student’s t-test was applied to test the hypothesis that *KLF2* expression was higher in the CTRL group than in the ELA group. For *KLF4*, a Welch’s Two Sample t-test was utilized to assess the same hypothesis under the assumption of unequal variances.

### Identification of genes with *KLF2* or *KLF4* binding motifs

Identification of genes that are potentially regulated by *KLF2/4* was performed using a snATAC-seq dataset previously generated by our group (75). This dataset includes comprehensive profiling of open chromatin across all cell types of the brain, including a vascular cluster. The Cisbp position weight matrix (PWM) was used to find motif occurrences of *KLF2/4* in any open chromatin peaks identified in the snATAC-seq dataset. To find target genes of *KLF2/4* transcription factor binding sites, we used Peak-to-gene (p2g) linkages, which were computed using Pearson correlation between snATAC-seq open chromatin peak accessibility and paired snRNA-seq gene expression within 500Kbp windows. Moreover, *KLF2/4* transcription factor motif containing p2g links were further restricted to those with the highest enrichment for these binding sites. In order to do this, we ran Homer **findMotifsGenome.pl with-find option for our motifs of interest.** As a result, Motif Score (log odds score of the motif matrix, higher scores are better matches) were obtained, and *KLF2/4* p2g links with motif scores≥median motif score across all peaks were retained (Median value=8.28). Finally, among filtered p2g links with highest evidence for *KLF2/4* transcription factor motifs, only peaks which also showed differentially high accessibility (FDR<0.05 and LOGFC>0.5) in the vascular cell type cluster compared to other brain cell types were kept.

## Results

### Immunophenotypic characterization of isolated brain microvessels: preserved morphology and expression integrity

The angioarchitecture of brain microvessels is composed of several vascular cell types. To characterize the composition and morphological integrity of microvessels isolated by our method, we utilized immunostaining to visualize and assess a range of NVU markers (Fig. 1a). We observed continuous expression of laminin (*LAM;* Fig. 1b), a marker of endothelial cells (76); vimentin (*VIM*; Fig. 1c), primarily found in smooth muscle cells (77); claudin5 (*CLDN5;* Fig. 1d), the primary constituent of endothelial tight junctions (78); Platelet-derived growth factor receptor beta (*PDGFR*β; Fig. 1e), a marker of pericytes (79); and aquaporin4 (*AQP4;* Fig. 1f), which is localized to astrocytic endfeet (80). Overall, our microvessel isolation methodology, as outlined in Wakid et al. (2023; 63), effectively captures the NVU in high yield and maintains microvessel structural integrity *ex vivo*.

**Fig. 1:**
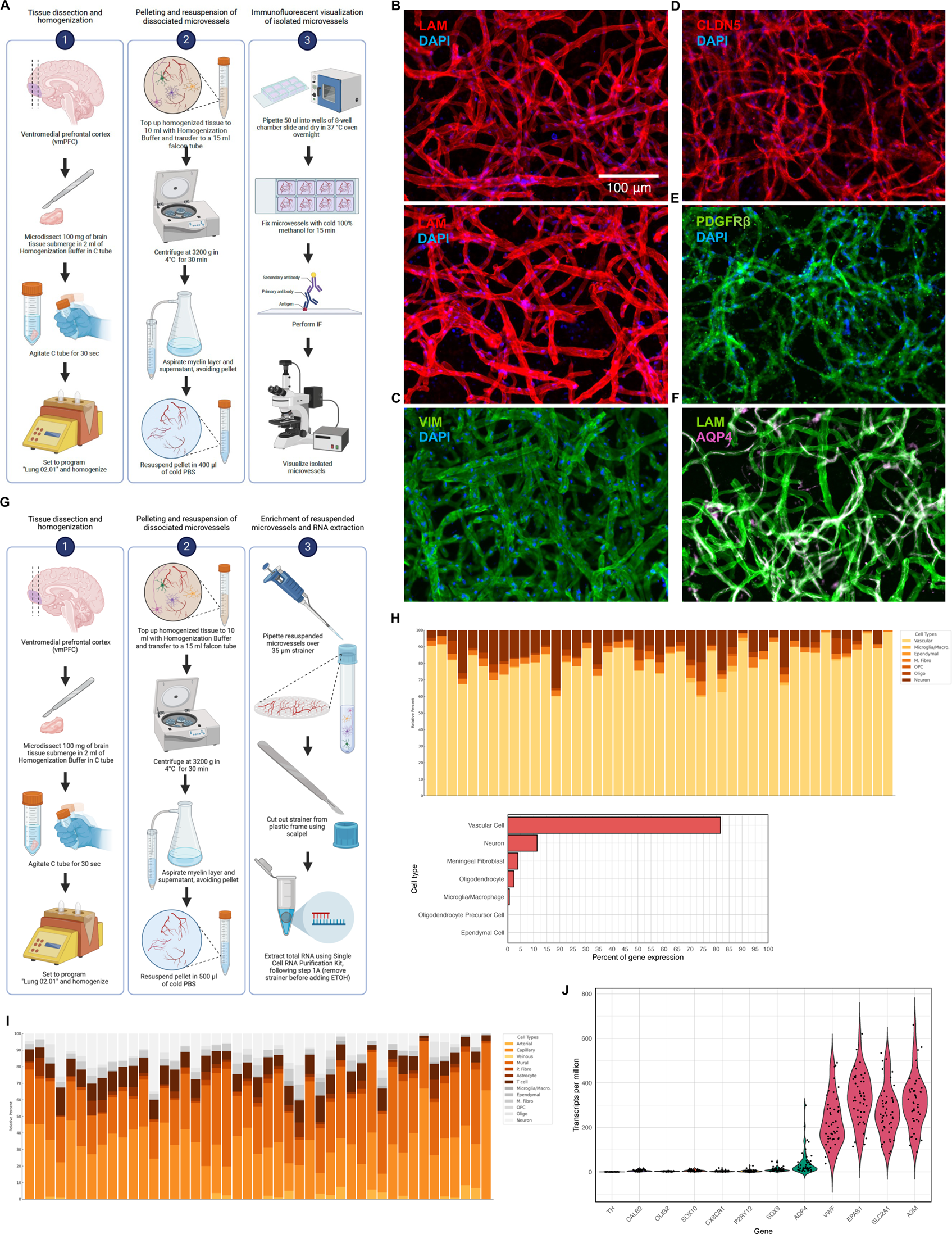
Effective isolation and enrichment of microvessels. a) Schematic overview of experimental workflow for isolating microvessels from postmortem brain tissue and processing for immunostaining of neurovascular markers. b) Representative micrograph of isolated microvessels immunostained with LAM (red). c) Representative micrograph of isolated microvessels immunostained with VIM (green). d) Representative micrograph of isolated microvessels immunostained with CLDN5 (red). e) Representative micrograph of isolated microvessels immunostained with PDGFRβ (green). f) Representative micrograph of isolated microvessels immunostained with LAM (green) and AQP4 (magenta). Nuclei were stained with DAPI (blue). Scale bar = 100µm and applies to all micrographs in panels b)-f). Bulk RNA sequencing and bioinformatic deconvolution of isolated microvessels. g) Schematic overview of experimental workflow for isolating microvessels from postmortem brain tissue and processing library construction and bulk RNA sequencing. h) Deconvolution of sequenced microvessel samples was performed via CIBERSORTX algorithm. Top: Bar plot showing cumulative percentage of vascular gene expression across sequenced subjects. Bottom: Bar plot displaying the percentage of vascular gene expression averaged across sequenced subjects, as well as percentages of non-vascular cell types averaged across sequenced subjects. These data highlight the significant enrichment of vascular cells within the microvessel RNA sequencing data. i) Bar chart showing significant enrichment of genes associated with capillary and mural cells, compared to those of arterial and venous endothelial cells. This pattern underscores the selective enrichment of microvessels, as opposed to larger diametre vessels along the arteriovenous axis. j) Transcript Per Million (TPM) values for different cell type-defining genes, demonstrating a predominance of vascular cell transcripts, particularly endothelial markers. In contrast, there is limited expression of the astrocytic endfeet marker *AQP4*, and minimal expression of the astrocytic nuclear marker *SOX9*, neuronal markers *TH* and *CALB2*, oligodendrocytic markers *OLIG2* and *SOX10*, and microglial markers *CX3CR1* and *P2RY12*.

### Profiling isolated microvessels of the human vmPFC by bulk sequencing Computational estimation of cell type proportion from sequenced microvessels shows enrichment for vascular cell types

To confirm the enrichment of microvessels and estimate the proportions of vascular cell types that have been enriched, we used the CIBERSORTX algorithm (65) and average reference signatures from 10X Chromium single-nucleus data of human brain vasculature (54) to deconvolute our data. Deconvolution of our data revealed a significant enrichment for vascular cell types across subjects (Fig. 1h), accounting for an average of 81.7% of the sequenced genes (representative plots for vascular markers and calculation methods in Supplementary Fig. 2). Importantly, a high representation of capillary (33.3%) and mural cell (32.6%) assigned genes were estimated (Fig. 1i), as is typically observed in the microvessel zone (81). In contrast, there was more limited representation of genes assigned to arterial (1.3%) but no venous endothelial cells, which corroborates with the intended objective of isolating the smallest of vessels from brain tissue, as well as lower estimations for astrocyte, perivascular fibroblast, and T cell genes (Fig. 1i). The average percentage estimate of summed vascular genes did not differ between groups (W=295, p-value=0.337; Supplementary Fig. 2), nor did the average percentage estimate for capillary genes (W=253, p-value=0.991; Supplementary Fig. 2) or mural cell genes (*t*(43)=0.91953, *p*=0.3629; Supplementary Fig. 2). As previously reported (59), microvessels isolated using the described method are obtained in high yield.

Although the protocol enriches for microvessels, it does not purify them; consequently, a limited proportion of non-vascular cell types are co-enriched in the collected pellets, as estimated by computational deconvolution (Fig. 1h-i). This outcome is to be expected when using postmortem tissue, as reported by others, a subset of “vasculature-coupled” neuronal and glial cells with distinct expression signatures (from the corresponding canonical cell types) are physically adhered to microvessels (53). Taking this into consideration, we compared deconvolution estimates per subject against two exclusion criteria: 1) neuronal contamination exceeding two standard deviations from the mean, and/or 2) oligodendrocyte contamination exceeding two standard deviations from the mean, as both scenarios were shown to skew subsequent analyses. Adhering to these criteria, 4 subjects (2 CTRL and 2 ELA); the matching between the groups remained unaffected. Transcript Per Million (TPM) values for cell type-defining genes in CTRLs and ELA subjects illustrate that our data predominantly reflect transcripts from vascular cells (Fig. 1j), as evidenced by the high expression of endothelial markers such as *VWF*, *EPAS1*, *SLC2A1*, and *A2M*.

### Neurovascular changes in the vmPFC of cases

Our understanding of neurovascular-related changes in psychopathologies and ELA remains limited in humans, and particularly with respect to NVU-specific sex differences. In this study, both male and female subjects were included in CTRL and ELA groups but with no prior evidence of major sex differences nor *a priori* hypothesis that sex differences manifest after ELA, we initially conducted DGE analysis with the sexes pooled together. We identified a total of 463 DEGs associated with a history of ELA irrespective of sex. Further examination, however, highlighted a proportion of DEGs showing marked sex-driven expression differences, most notably among females, thus, the potential influence of sex on gene expression was thoroughly investigated through several rigorous analytical avenues (Supplementary Fig. 2-3; Supplementary Table 3). The observation of sexual dimorphism, coupled with the presence of approximately double the number of males compared to females in both groups, prompted our decision to analyze male and female counts separately and, as a result, characterize the sex-specific effects of ELA on gene expression.

### Sex-driven differences in the vmPFC neurovascular transcriptome of cases

To identify sex-specific differences in gene expression patterns between CTRL and ELA groups, we performed DGE analysis separately in males and females. In males, we identified a total of 34 DEGs, with 19 upregulated and 15 downregulated genes in cases vs controls (p.adj < 0.05, ∣log2(FC)∣≥log2(1.1); Fig. 2a-c, Supplementary Table 4). This contrasted with females which displayed 774 DEGs, with 343 upregulated and 431 downregulated genes in cases vs controls (p.adj < 0.05, ∣log2(FC)∣≥log2(1.1); Fig. 2e-g, Supplementary Table 5). The magnitude to which the female NVU was affected was greater, with an average log2(FC) of 0.645 for females (equivalent to a fold change of approximately 1.564, or a 56% change in expression), compared to an average log2(FC) of 0.350 for males (equivalent to a fold change of approximately 1.275, or a 28% expression change). This indicates that the female NVU experiences substantially greater transcriptomic disturbances in depressed suicides with a history of ELA compared to males. The robustness of these sex differences was supported by a sub-sampling analysis (Supplementary Fig. 3).

**Fig. 2:**
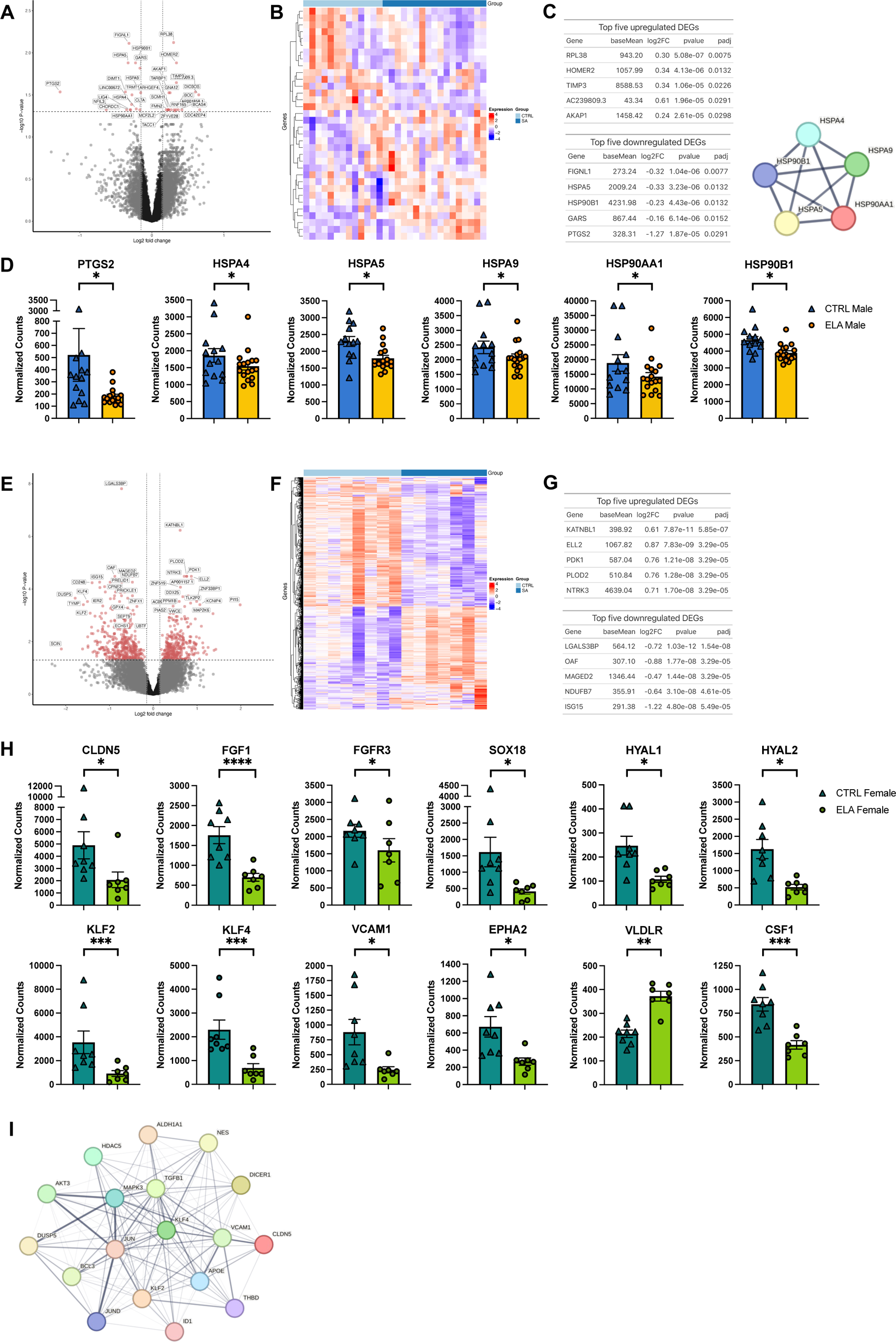
DGE of CTRL versus ELA for male and female microvessels, respectively. a) Volcano plot showing the results from differential gene expression analysis between male CTRL and ELA groups, performed using DESeq2. Data points represent individual genes, with significance determined by adjusted p-value (padj) < 0.05 and absolute log2 fold change > log2(1.1). Genes above the threshold line are significantly differentially expressed. b) Heatmap showing the results from differential gene expression analysis between male CTRL and ELA groups, performed using DESeq2. Each row represents a gene, and the color intensity corresponds to the magnitude of log2 fold change, with red indicating upregulation and blue indicating downregulation. The heatmap is organized to display genes meeting the significance criteria of adjusted p-value (padj) < 0.05 and |log2(fold change)| > log2(1.1). The clustering tree alongside the heatmap categorizes genes based on the similarities in their expression changes, effectively grouping genes with similar expression patterns together. c) Tables displaying the top 5 upregulated (top) and top 5 downregulated (bottom) DEGs for males, indicating their gene names, log2 fold changes, and adjusted p-values. d) Bar plots showcasing selected DEGs in males. Normalized counts for PTGS2, HSPA4, HSPA5, HSPA9, HSP90AA1, and HSP90B1 show a downregulation in the male ELA group compared to the male CTRL group. (*: p < 0.05; **: p < 0.01; ***: p < 0.001; ****: p < 0.0001). e) Volcano plot showing the results from differential gene expression analysis between female CTRL and ELA groups, performed using DESeq2. Data points represent individual genes, with significance determined by adjusted p-value (padj) < 0.05 and absolute log2 fold change > log2(1.1). Genes above the threshold line are significantly differentially expressed. f) Heatmap showing the results from differential gene expression analysis between female CTRL and ELA groups, performed using DESeq2. Each row represents a gene, and the color intensity corresponds to the magnitude of log2 fold change, with red indicating upregulation and blue indicating downregulation. The heatmap is organized to display genes meeting the significance criteria of adjusted p-value (padj) < 0.05 and |log2(fold change)| > log2(1.1). The clustering tree alongside the heatmap categorizes genes based on the similarities in their expression changes, effectively grouping genes with similar expression patterns together. g) Tables displaying the top 5 upregulated (top) and top 5 downregulated (bottom) DEGs for females, indicating their gene names, log2 fold changes, and adjusted p-values. h) Bar plots showcasing normalized counts for selected DEGs in females, namely CLDN5, FGF1, FGFR3, SOX18, HYAL1, HYAL2, KLF2, KLF4, VCAM1, EPHA2, VLDLR, and CSF1. (*: p < 0.05; **: p < 0.01; ***: p < 0.001; ****: p < 0.0001). STRING analysis reveals DEGs shown in g) are involved in a network of protein-protein interactions. i) STRING analysis for KLF2 and KLF4, both identified as DEGs in females, demonstrates their interaction with other female DEGs.

To assess whether critical genes at the NVU were impacted in cases, we cross-referenced DEGs identified in males and females with vascular cell type defining markers from recent studies by Yang et al. (2022; 54) and Garcia et al. (2022; 53) (Supplementary Fig. 3). Male DEGs included *PTGS2*, a marker of BMECs, along with *TACC1*, *TIMP3*, *CDC42EP4*, and *DIO3OS*, all of which exhibit high expression levels in pericytes. Conversely, the female NVU revealed a distinct signature, showcasing DEGs that included several vascular cell type defining markers. Noteworthy among these are *CSF1* (log2(FC)=-0.77, p.adj=0.0003, Fig. 2h), essential for monocyte maintenance (35508166, 24890514) and proliferation (82–84); *FGF1* (log2(FC)=-0.89, p.adj=0.00023, Fig. 2h), a potent mitogen and angiogenic factor (85); and its receptor *FGFR3* (log2(FC)=-0.70, p.adj=0.038, Fig. 2h).

Binding partners *KLF2* (log2(FC)=-1.44, p.adj=0.00083, Fig. 2h-i) and *KFL4* (log2(FC)=-1.50, p.adj=0.0003, Fig. 2h-i) were both downregulated in female cases (but not males with ELA, *KLF2*: p.adj=0.92; *KLF4*: p.adj=0.84). Belonging to the Krüppel-like family of transcription factors, *KLF2/4* are enriched in the endothelium and orchestrate vascular homeostasis, serving as a central transcriptional switch point between a proinflammatory, atheroprone versus quiescent, atheroresistant endothelial phenotype. *KLF2/4* determine endothelial transcription programs (>1000 genes) (86) that control key functional pathways such as cell migration, vasomotor function, hemostasis (86–88), as well as barrier integrity by induction of multiple anti-inflammatory, anti-thrombotic factors (89, 90), and by regulating endothelial expression of *CAMs*, *NF-kB*, and tight junction protein *CLDN5* (90–93). Similarly downregulated in females is *CLDN5* (log2(FC)=-0.65, p.adj=0.012, Fig. 2h), as previously observed in models of chronic stress (38–40). Other female DEGs with well characterized functions at the NVU include *VLDLR*, *HSPB1*, *HYAL1*, *HYAL2*, *ST6GALNAC3*, *ADGRA2*, *BST2*, *VCAM1*, *IL32*, *SOX18*, *MT2A*, *CTSB*, *RGL3*, *PAG1*, and *GJA4*. Additionally, several ATP-binding cassette transporters and solute carriers are also differentially expressed, namely *ABCD3*, *ABCA5*, *SLC24A4*, *SLC36A4*, *SLC4A7*, *SLC25A36*, *SLC9A3R2*, *SLC35A4*, *SLC6A6*, *SLC39A14*, *SLC8B1*, *SLC22A3*, *SLC7A10*.

A hypergeometric test revealed a statistically significant overlap between female DEGs and vascular cell type defining markers (hypergeometric *p*-value=2.48×10^-7^). Focusing solely on DEGs that overlap with vascular cell type defining markers, *KLF2* emerged as highly significant, and the foremost with a well-characterized function. Thus, we chose to experimentally validate the expression changes of *KLF2/4* using FISH in the same female CTRL and ELA subjects that were previously sequenced. Consistent with our RNA sequencing findings, significant downregulation of *KLF2* (t(12)=8.7474, p=7.45×10^-7^) and *KLF4* (t(7.2153)=2.3976, p=0.023) expression was observed in the grey matter microvessels of ELA vs CTRL females (Fig. 3a-b). Importantly, there was no statistically significant difference in blood vessel density between the CTRL and ELA females (W=38, p=0.09732).

**Fig. 3:**
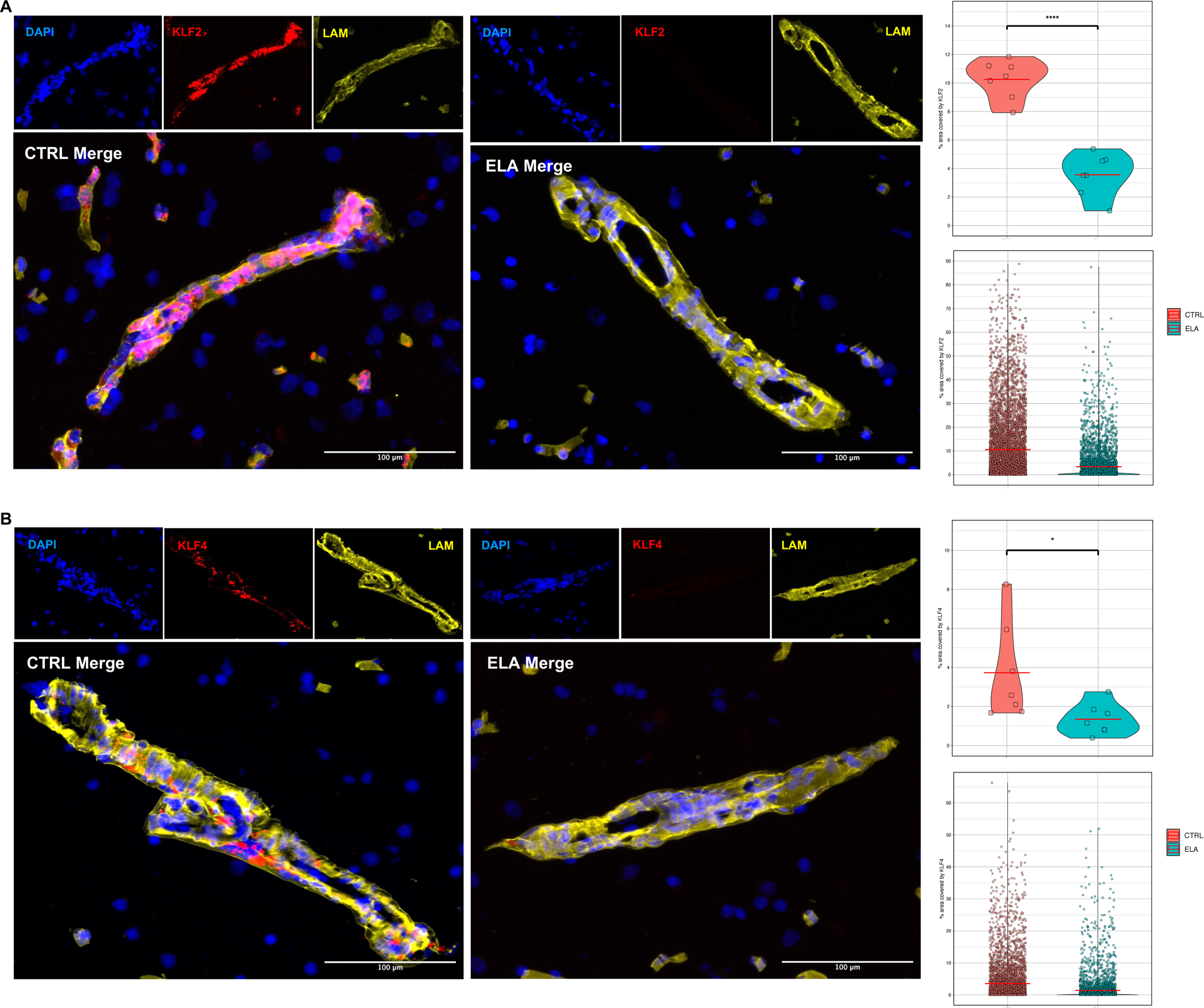
In situ validation of decreased KLF2 and KLF4 expression in grey matter microvessels of females with ELA compared to female CTRLs. a) Representative images of FISH for KLF2 (red) expression in CTRL (left) and ELA (right) microvessels (*LAM*+ cells, yellow). Nuclei were counterstained with DAPI (blue). Scale bar = 100 µm. Mean microvessel expression of KLF2 is significantly reduced in females with ELA (p = 7.45 × 10^-7^). This is further demonstrated by the distribution of individual data points for KLF2 expression across the two groups, with a higher percent of microvessel area covered by KLF2 in the CTRL group compared to the ELA group. b) Representative images of FISH for KLF4 (red) expression in CTRL (left) and ELA (right) microvessels (*LAM*+ cells, yellow). Nuclei were counterstained with DAPI (blue). Scale bar = 100 µm. Mean microvessel expression of KLF4 is significantly reduced in females with ELA (p = 0.023). This is further demonstrated by the distribution of individual data points for KLF2 expression across the two groups, with a higher percent of microvessel area covered by KLF4 in the CTRL group compared to the ELA group.

### Females – but not males-display significant neurovascular dysfunction in cases

The relevance of the DEGs we identified in relation to psychiatric disorders was assessed using the PsyGeNET (67) text-mining database. Upon evaluating male DEGs, we found that depressive disorders exhibited the most gene-disease associations (>9 DEGs; Fig. 4a; Supplementary Fig. 4). In the case of female DEGs, schizophrenia had the highest number of gene-disease associations (60 DEGs; Fig. 4b; Supplementary Fig. 4), closely followed by depressive disorders (>50 DEGs; Fig. 4b). Therefore, our microvessel derived-DEG findings in both sexes, especially in females, corroborate previously reported gene-disease associations.

**Fig. 4:**
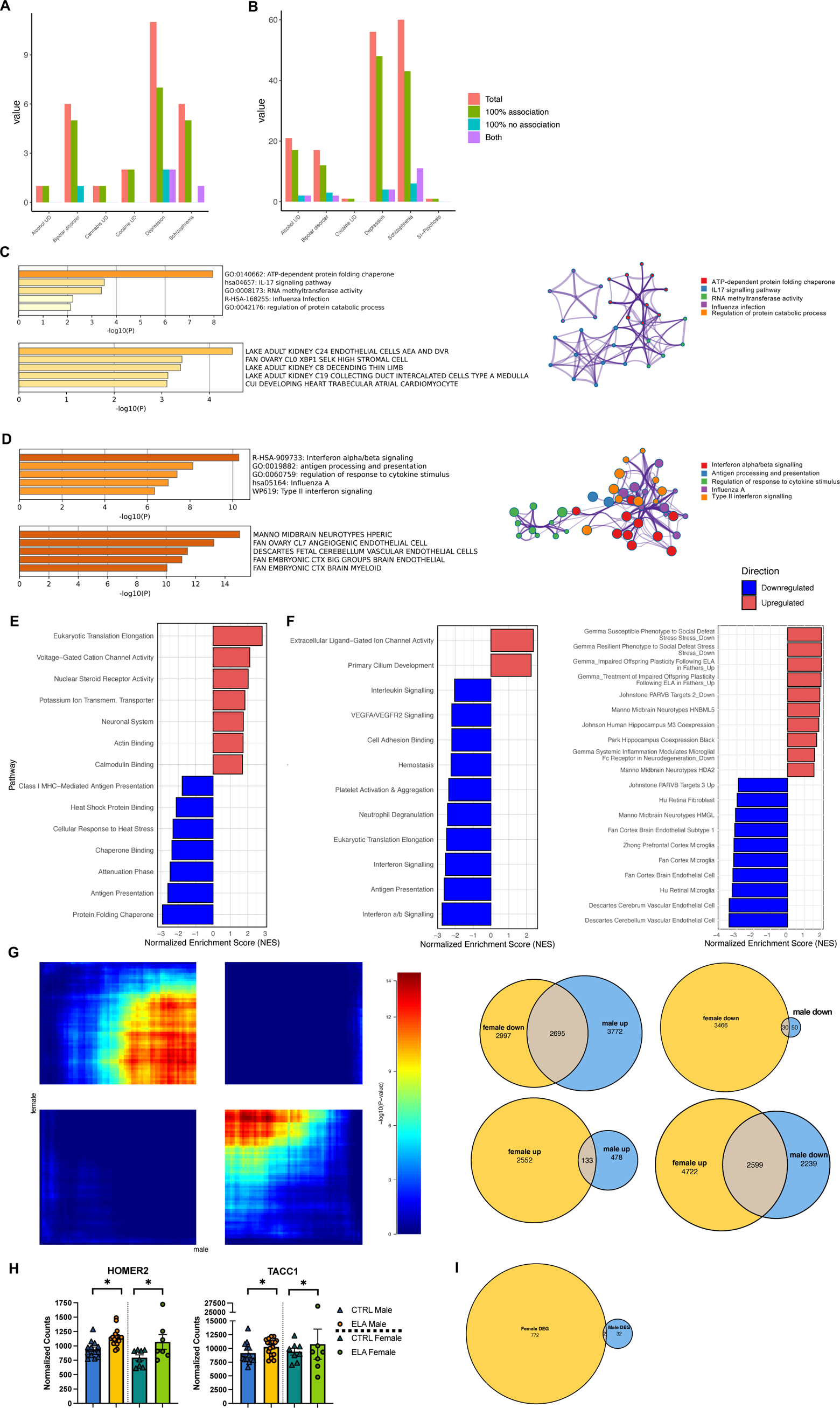
Functional annotation and enrichment in male and female DEGs. a-b) Bar plot displaying the number of gene-disease associations for psychiatric disorders in a) male DEGs and b) female DEGs, queried via the PsyGeNET database. c-d) Top results for (top left) functional annotation and (bottom left) cell-type enrichment of c) male and d) female DEGs were conducted using Metascape. (Bottom) The resultant clusters are represented in a network plot where nodes represent enriched terms and edges represent relationships between terms (right). Terms with a p-value < 0.05, a minimum count of 3, and an enrichment factor > 1.5 were considered significant. e-f) Results from fGSEA in e) males and f) females, referencing merged data from Reactome, Gene Ontology Molecular Function (GOMF), KEGG, and WikiPathways (WP). Upregulated pathways are indicated in red, while downregulated pathways are shown in blue, each according to their Normalized Enrichment Score (NES). Comparative analysis of male and female DEGs. g) (Left) Hypergeometric heatmap from RRHO analysis comparing transcriptional changes between males and females with ELA. (Right) Corresponding Venn diagrams demonstrating the number of genes with shared or distinct directional changes between the two sexes. h) Bar plots showing the differential expression of HOMER2 and TACC1 between CTRL and ELA groups for both males and females. (*: p < 0.05). i) Venn diagram showing that only two genes (HOMER2 and TACC1) are shared DEGs between males and females with ELA.

Functional annotation and cell-type enrichment of DEGs were conducted using Metascape (68). In male cases, the "ATP-dependent protein folding chaperone" functional cluster was as the most significantly enriched (Fig. 4c), pertaining to DEGs *HSPA4*, *HSPA5*, *HSPA9*, *HSP90AA1*, and *HSP90B1* (Fig. 2d). Additionally, there was an endothelial cell type signature enrichment amongst male DEGs (Fig. 4c). Functional enrichment of female DEGs revealed significantly enriched functional clusters related to immune and vascular pathways, namely “Interferon alpha/beta signalling” and “Regulation of response to cytokine stimulus” (Fig. 4d). Furthermore, there was a pericyte and endothelial cell type signature enrichment amongst DEGs in female. Further scrutiny of female DEGs with known immune functions confirmed “global immune suppression” where, irrespective of whether pro-or anti-inflammatory in nature, immune-related DEGs were downregulated, some of which are presented in Table 2 (Supplementary Fig. 12). To assess the functional significance of gene expression differences in females, we examined whether the proteins encoded by DEGs were part of interacting networks (see Supplementary Fig. 12).

**Table 2:** A representative list of significantly downregulated genes with known immune functions in females with ELA includes genes with both pro-inflammatory and anti-inflammatory roles. This widespread downregulation of immune-related genes suggests a ‘global immune suppression’.

| Immune-related gene expression in females with ELA |  |  |  |  |
| --- | --- | --- | --- | --- |
| Gene | baseMean | log2FC | pvalue | padj |
| IFI6 | 808.29 | -0.85 | 0.0003 | 0.0138 |
| HLA-B | 7025.47 | -0.92 | 0.0011 | 0.0299 |
| HLA-F | 905.24 | -0.62 | 0.0009 | 0.0262 |
| IFI35 | 171.00 | -0.88 | 0.0000 | 0.0002 |
| IFIT3 | 701.97 | -0.88 | 0.0000 | 0.0049 |
| IRF1 | 488.35 | -1.17 | 0.0004 | 0.0168 |
| IRF7 | 162.96 | -0.76 | 0.0014 | 0.0355 |
| ISG20 | 153.28 | -0.67 | 0.0022 | 0.0459 |
| ISG15 | 291.38 | -1.22 | 0.0000 | 0.0001 |
| IRF9 | 726.61 | -0.42 | 0.0003 | 0.0144 |
| CD74 | 1945.60 | -1.11 | 0.0001 | 0.0065 |
| HLA-DMA | 84.91 | -1.12 | 0.0005 | 0.0198 |
| HLA-DMB | 151.62 | -1.29 | 0.0014 | 0.0358 |
| HLA-DPA1 | 344.01 | -1.11 | 0.0000 | 0.0001 |
| HLA-DPB1 | 233.66 | -1.00 | 0.0001 | 0.0096 |
| HLA-DRA | 259.14 | -1.33 | 0.0001 | 0.0097 |
| TAP1 | 583.88 | -0.86 | 0.0000 | 0.0001 |
| TAP2 | 634.83 | -0.53 | 0.0023 | 0.0460 |
| CSF1 | 643.41 | -0.77 | 0.0000 | 0.0003 |
| IRAK1 | 408.95 | -0.42 | 0.0021 | 0.0449 |
| IKBKE | 41.79 | -1.08 | 0.0010 | 0.0284 |
| IFNGR2 | 339.98 | -0.51 | 0.0001 | 0.0068 |
| TGFB1 | 510.19 | -0.54 | 0.0000 | 0.0029 |
| VCAM1 | 583.52 | -1.61 | 0.0005 | 0.0189 |
| CCRL2 | 41.08 | -0.85 | 0.0021 | 0.0448 |

We used GSEA (Fig. 4e-f) to explore the biological pathways underlying transcriptomic changes observed in male and female cases. For males, significant enrichment was found in pathways “Voltage-gated potassium channels” and “Muscle contraction” (Fig. 4e), suggesting potential alterations in K_V_ channel facilitated smooth muscle contraction (94–96). The "Nuclear Steroid Receptor Activity” pathway was similarly enriched (Fig. 4e), featuring DNA-binding transcription factors such as *PGR*, *ESR2*, and *PPARA*, which are recognized for their neuroprotective and broad vascular functions (97, 98). In contrast, chaperone binding protein pathways had significantly lower normalized enrichment scores (Fig. 4e). Similar to males, most significant pathways identified by GSEA in females were downregulated. Females exhibited downregulation in immune-related pathways, akin to ORA findings, including "Interferon signalling" and "Interleukin signalling” (Fig. 4f), indicating suppressed immune signalling in ELA. Additionally, gene sets related to vascular functions, namely “Platelet activation and aggregation” and “Hemostasis” were negatively enriched in females. Intriguingly, exposure to chronic stress is associated with prolonged proinflammatory platelet bioactivity (99–102) and platelet pathology in MDD (103–111), with greater effects in depressed women (112). Intriguingly, analysis using the newly curated gene set list Brain.GMT (71) (Fig. 4f) shows upregulation in genes typically downregulated in “Susceptible Phenotype to Social Defeat Stress" and in “Resilient Phenotype to Social Defeat Stress". Thus, our GSEA of microvessels from cases revealed dysregulated pathway gene sets that are functionally significant in both vascular cell types and the ELA phenotype. Furthermore, these findings align with DEGs identified in males and females.

### Little overlap between male and female neurovascular dysfunction

We next used RRHO analysis (72) to identify shared transcriptional changes between males and female cases. Consistent with previous studies showing distinct brain transcriptomic changes in males and females with MDD (113), we found remarkably little overlap between male and female ELA transcriptional signatures (Fig. 4g); thus, the directionality of the changes observed in males and females with ELA are not preserved. The limited overlap in transcriptional changes between male and female depressed suicides with a history of ELA was further confirmed by focusing our analysis exclusively on DEGs. By directly comparing male and female DEGs, we observed minimal commonality by identifying only two shared DEGs: *HOMER2* (ELA males: p.adj=0.013, log2(FC)**=**0.34; ELA females: p.adj=0.02, log2(FC)**=**0.63) and *TACC1* (ELA males: p.adj=0.048, log2(FC)**=**0.22; ELA females: p.adj=0.03, log2(FC)**=**0.47), both of which are upregulated in both sexes (Fig. 4h-i). While Homer proteins are best known for their interaction with group 1 metabotropic glutamate receptors (*mGluR1/5*) (114), *HOMER2* is also expressed in endothelial cells (Supplementary Fig. 4) and plays a crucial role in regulating Ca² signalling and activation of platelets by agonists such as thrombin (115, 116). Meanwhile, *TACC1*, a member of the Transforming Acidic Coiled-Coil family, acts as a coregulator of various nuclear receptors (117) that modulate transcription by binding to their target DNA response elements. Overall, these findings highlight a strong sexual dimorphism in the transcriptional signatures exhibited by the NVU in depressed suicides with a history of ELA.

### WGCNA identifies specific gene networks in the NVU of female cases

Having characterized the broad pattern of transcriptome-wide changes at the female NVU, we then sought to resolve specific gene coexpression networks that could be critical in determining NVU dysfunction in cases. Using WGCNA, we constructed a gene coexpression network that returned 73 modules. To gain insight into the biology of implicated modules, we identified a subset of interesting modules for further analysis based on two criteria: a significant association with cases (p-value<0.05) and a correlation coefficient greater than 0.7, ensuring a focus on modules strongly associated with depressed suicides with ELA. A total of 5 modules exhibited a significant correlation coefficient greater than 0.7 (Fig. 5a-b), as well as biological processes pertinent to ELA: membrane trafficking (Fig. 5c; turquoise: p=1.46×10^−7^, 1.74x), interferon alpha/beta signalling (Fig. 5d; black: p=8.95×10^−9^, 6.95x), *VEGFA*/*VEGFR2* signalling (green: p=2.04×10^−8^, 2.10x), response to lipopolysaccharide (greenyellow: p=2.88×10^−12^, 4.64x), and eukaryotic translation elongation (orange: p=1.10×10^−34^, 28.36x). Enrichment of cell-type associated genes in these modules identified microglial genes in turquoise (Fig. 5c; p=1×10^−14^, 2.3x), endothelial genes in black (Fig. 5d; p=1×10^−15^, 3.4x), green (p=1×10^−34^, 3.7x), and greenyellow (p=1×10^−14^, 5.2x), along with T cells in orange (p=1×10^−3^, 17x). Modules were then handed over to the GeneOverlap R package, where we assessed significant overlap of module genes with female DEGs. Of the 73 generated modules, this subset of 5 modules were also enriched for female DEGs (Fig. 5e; turquoise: p=2.3×10^−17^; green: p=4.5×10^−30^; black: p=1.4×10^−27^; greenyellow: p=2.5×10^−8^; orange: p=3.8×10^−6^), in a way that was consistent with regional RRHO patterns. Indeed, many of the co-downregulated genes in the RRHO analysis were also found within these 5 modules (e.g. *PDGFA*, *TFRC*, *EPHA3*, *FN1*, *CX3CL1*, *ICAM1*, *PTGS2*, *PGF*; Supplementary Fig. 5). Together, these analyses suggest that the turquoise, green, black, greenyellow, and orange modules may be particularly important in governing NVU dysfunction in female cases, which is further substantiated by a markedly lower median for each module’s eigengenes (Fig. 5f).

**Fig. 5:**
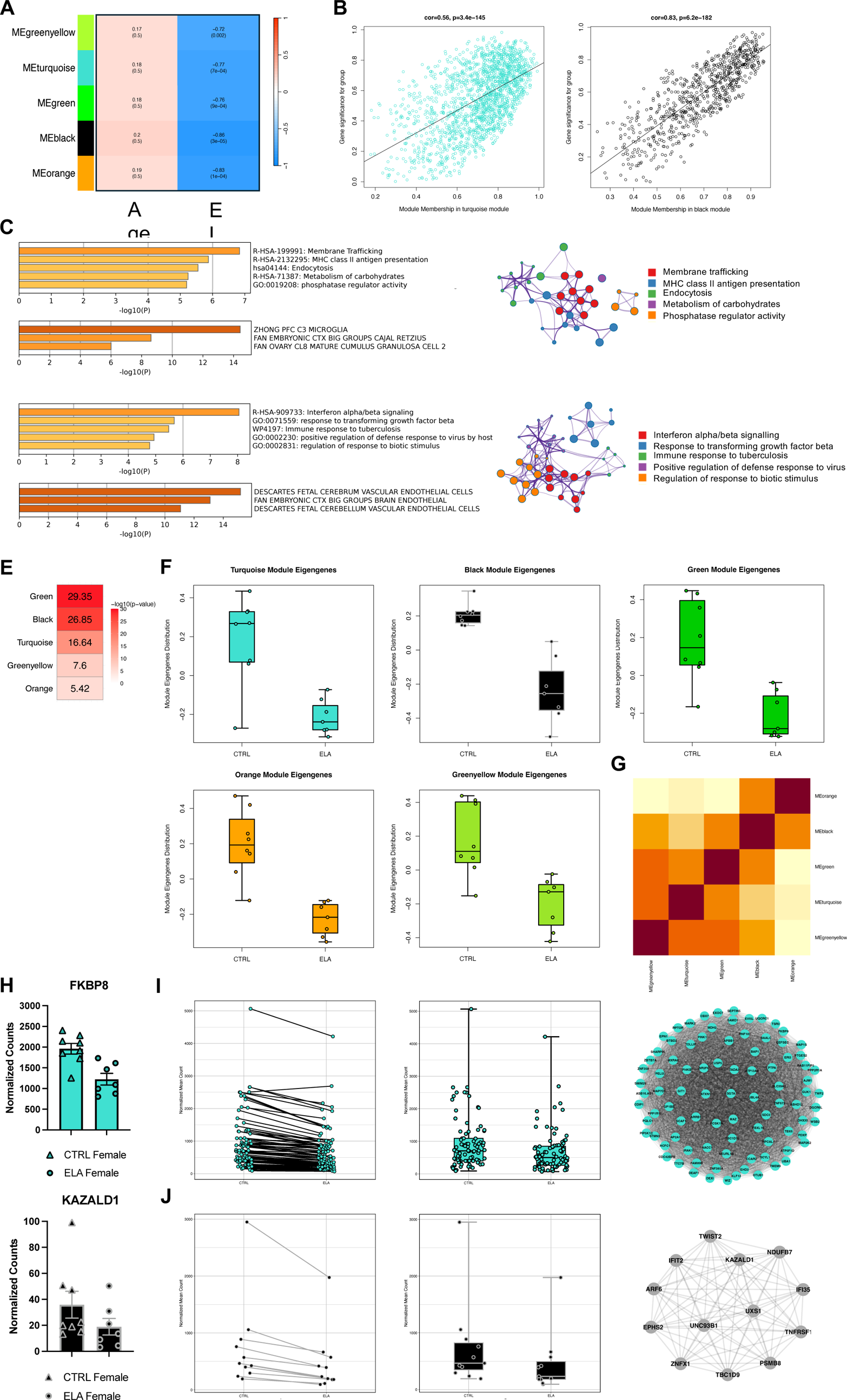
Assessing networks of gene co-expression and ELA using data specifically from females. a) (Left) Plot showing the five WGCNA modules (greenyellow, turquoise, green, black, orange) with significant correlation (coefficients greater than 0.7) with ELA in females. Modules with negative correlation values are shown in blue, indicating their inverse relationship with ELA. b) Scatter plots for module membership versus gene significance for turquoise (cor = 0.56, p = 3.4×10−145), black (cor = 0.83, p = 6.2×10−182), and greenyellow (cor = 0.72, p = 1.2×10−181) modules. c-d) Top results for (Top left) functional annotation and (bottom left) cell-type enrichment of c) turquoise and d) black modules were conducted using Metascape. (Right) The resultant clusters are represented in a network plot where nodes represent enriched terms and edges represent relationships between terms. Terms with a p-value < 0.05, a minimum count of 3, and an enrichment factor > 1.5 were considered significant. e) Fisher’s Exact Test was performed via GeneOverlap to assess genes common between female DEGs and the turquoise, black, green, greenyellow, and orange modules. f) Box and whisker plots showing median eigengene values across turquoise, black, green, orange and greenyellow modules between CTRL and ELA females. g) Correlation plot between the eigengenes of the turquoise, black, green, orange and greenyellow modules. h) (Top) Bar plots showing normalized counts for turquoise top hub gene FKBP8 and black top hub gene KAZALD1 between CTRL and ELA females. i-j) (Left) scatter plot showing i) turquoise and j) black hub genes between CTRL and ELA females. (Middle) Box and whisker plots showing i) turquoise and j) black hub genes between CTRL and ELA females. (Right) Network plots of i) turquoise and j) black hub genes.

### KLF2 and KLF4 regulate endothelial activation transcriptional networks in female cases

Interestingly, *KLF2* and *KLF4* (along with *CLDN5*) emerged in modules turquoise and black, respectively (Supplementary Table 6). Recognizing that these modules may represent genes for which *KLF2/4* are nodal regulators, we assessed potential correlation between the eigengenes of the turquoise and black modules. A strong positive correlation (r = 0.51, Fig. 5g) was observed, suggesting some level of co-regulation or functional overlap. Importantly, *KLF2* not only exhibits a strong correlation with the turquoise eigengene (MM = 0.76) and a strong association with ELA (GS = 0.83), but its closeness centrality (CC = 0.12) also indicates a functionally central position within the module. *KLF4* similarly demonstrates a strong correlation with the black eigengene (MM = 0.86), a strong association with ELA (GS = 0.86), and a closeness centrality of 0.15, underscoring its central role within the module. We next sought to explore the potential impact of *KLF2/4* downregulation, as observed in female cases, on genes identified within the turquoise and black modules. To this end, genes within these modules were cross-referenced against human homologues of genes experimentally validated to be affected by dual *KLF2/4* knockout. This comparative analysis revealed substantial overlap (Supplementary Fig. 5): 448 genes within the turquoise module (p-value = 5.1×10^−9^) and 193 genes within the black module (p-value = 3.3×10^−6^). Furthermore, additional *KLFs* are present in modules green (*KLFs 8*, *10*, and *16*), greenyellow (*KLF6*), and turquoise (*KLF13*), and modules that possess *KLFs* exhibit strong correlation with one another (Fig. 5g); thus, it is conceivable that these *KLFs* may also play a regulatory role within their respective modules. Hub genes, particularly the topmost hub genes (Fig. 5h), within the turquoise and black modules exhibit decreased mean normalized count in depressed females with a history of ELA (Fig. 5i).

Using a snATAC-seq dataset generated by our group (75), we then explored potential regulatory actions performed by *KLF2/4* in the turquoise and black modules, respectively. The Cisbp PWM was used to identify DNA sequences within the open chromatin peaks that matched known motifs for *KLF2* or *KLF4*. These motifs were then analyzed in conjunction with corresponding open chromatin regions that were highly accessible in the vascular cluster (Fig. 6a) compared to non-vascular cell types. This approach allowed us to filter peak-to-gene (p2g) associations for peaks covering DNA sequences matching known motifs for *KLF2* or *KLF4* and combined with corresponding open chromatin regions that were highly accessible in the vascular cluster (Fig. 6a) compared to non-vascular cell types. By virtue of this approach, specific genes that are potentially regulated by *KLF2* or *KLF4* in endothelial cells can be identified. Disregarding the location of potential *KLF2/4* binding sites, genes with motif scores ≥ median score and significant FDR (FDR ≤ 0.05; indicating high confidence that the identified peak is a true regulatory site where *KLF2* or *KLF4* is likely to bind) were then cross-referenced with turquoise and black module genes, respectively, demonstrating that 33% of turquoise module genes are motif-linked to *KLF2* (Fig. 6b; p-value = 1.8×10^−93^) and that 30% of black module genes are motif-linked to *KLF4* (Fig. 6c; p-value = 1×10^−14^). Several hub genes were identified amongst motif-linked genes in both the turquoise and black modules, including several *KLF2* binding sites in distal, promoter, exonic and intronic regions of the topmost turquoise hub gene *FKBP8* (further breakdown of gene regions with potential *KLF2/4* binding sites found in Supplementary Table 7). These findings underscore the substantial influence of *KLF2/4* on the transcriptional landscape of endothelial cells, specifically within the promoter and distal regions, which are crucial for gene regulation.

**Fig. 6:**
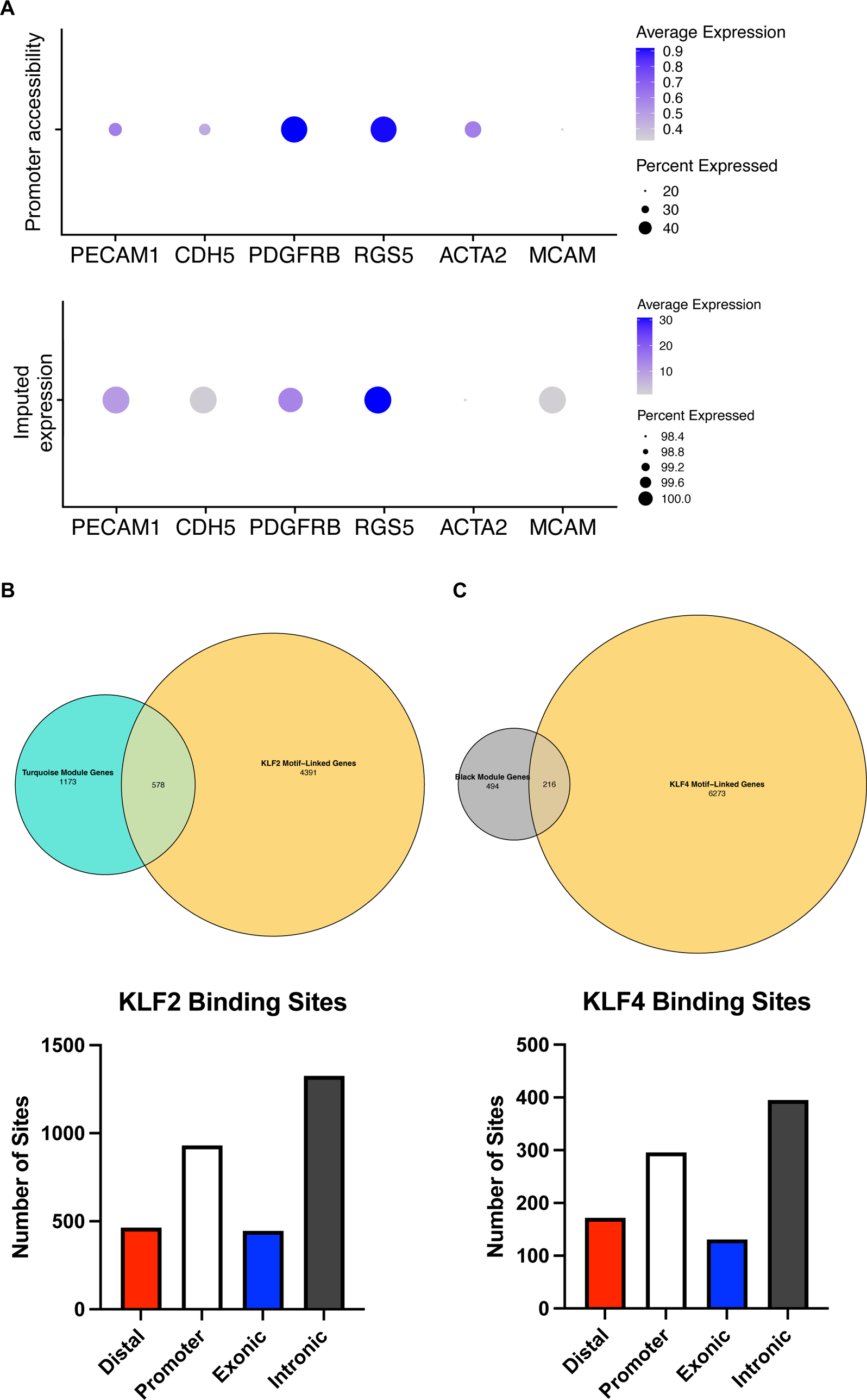
Identification of turquoise and black module genes motif-linked to KLF2 and KLF4, respectively. a) (Top) Promoter accessibility across key endothelial (PECAM1, CDH5), pericyte (PDGFRB, RGS5) and smooth muscle cell (ACTA2, MCAM) markers in the vascular cluster from ATACseq data (Chawla et al., 2024). Average expression indicates the average chromatin accessibility score for the promoter region (where the promoter region is defined as within 2kbp from gene transcription start site) of the listed genes in the vascular cluster. Percent expressed indicates the percentage of cells within the vascular cluster where the promoter region of these genes are accessible. (Bottom) Imputed expression of key endothelial (PECAM1, CDH5), pericyte (PDGFRB, RGS5) and smooth muscle cell (ACTA2, MCAM) markers in the vascular cluster from single nuclei RNA sequencing data (Chawla et al., 2024). Average expression indicates the average count for the gene and percent expressed indicates the percentage of cells within the vascular cluster that expression the gene. These results show a high expression across the cluster, indicating a mixed cell population with higher proportions of endothelial cells and pericytes. Expression was imputed cell-type wise, so vascular cluster snRNA data was integrated with vascular cluster snATAC data. b) (Top) Venn diagram showing overlap between KLF2 motif-linked genes and turquoise module genes, (Bottom) Bar plot showing the cumulative number of KLF2 binding sites found at distal, promoter, exonic, and intronic regions across the 577 turquoise genes with KLF2 binding sites. c) (Top) Venn diagram showing overlap between KLF4 motif-linked genes and turquoise module genes, (Bottom) Bar plot showing the cumulative number of KLF4 binding sites found at distal, promoter, exonic, and intronic regions across the 216 turquoise genes with KLF4 binding sites.

## Discussion

The pernicious effects of adversity experienced during developmental periods of heightened plasticity linger as psychopathology and disease into adulthood. The complex neurobiological and stress-mediated mechanisms through which these enduring impacts manifest, however, remain insufficiently characterized in individuals with histories of ELA. One proposed mechanism is NVU dysfunction: chronic stress leads to the production of blood-borne inflammatory cytokines and activation of leukocytes, which then act on BMECs (118). This interaction instigates a temporal sequence of biological pathways that orchestrate the vascular response, potentially impacting the properties of the BBB and, consequently, affecting the brain parenchyma. As developmental windows of critical plasticity close, the vascular response is redirected towards "off-trajectory" pathways, which are then sustained throughout life. In this study, we used our recently developed protocol (59) to isolate intact microvessels from archived snap-frozen vmPFC grey matter samples from healthy controls and depressed suicides with histories of ELA, for which we demonstrate preserved morphological integrity as well as expression of canonical vascular cell type markers. We isolated intact microvessels in favour of investigating the whole NVU as opposed to dissociated, single nuclei. Our decision to do so lies in the fact that a structurally preserved unit can provide insight into the neurovasculature in a manner that dissociated nuclei cannot: not only is unwanted, technically-related depletion of vascular-associated cells avoided, but cytoplasm – which carries much higher amounts of mRNA and protein – as well as interstitium, are also preserved. This is not trivial, as work done by others comparing microglia single cells versus single nuclei from postmortem tissue reveals that certain populations of genes are depleted in nuclei (119). Ultimately, the NVU is so interconnected that dysfunction is typically observed in all vascular-associated cells, conferring whole-unit dysfunction that becomes difficult to parse when looking at dissociated nuclei.

Total RNA extracted from isolated microvessels was subjected to RNA sequencing to generate the first NVU-specific transcriptomic profile in individuals with a history of ELA. Overall, our findings suggest that NVU function and integrity in these individuals is impacted through widespread gene expression changes. Transcriptomic changes associated with ELA or chronic stress have been the focus of several investigations, yet the majority of these studies exclusively characterize male ELA signatures (120–126). In contrast, fewer studies include females (127–129), thereby omitting the opportunity to integrate or make direct comparisons between males and females and, as such, the extent to which the transcriptional profiles defining ELA differs in males versus females remains unknown. Our data-informed decision to perform DGE analysis on males and females separately allowed us to identify dramatic, fundamental differences in NVU dysfunction in males versus females, providing a framework for better understanding the molecular basis of the sexual dimorphism characterizing depression and ELA. The male NVU appears to be relatively unaffected in male cases; in contrast, the female NVU experiences widespread differential expression in a large number of genes. These findings are further supported by outcomes from GSEA and RRHO analyses, revealing a lack of overlap in both pathway dysregulation and the direction of gene expression changes across all genes, not solely those that are significantly differentially expressed.

Male DEGs showed enrichment for pathways delineating voltage-gated potassium (K_V_) channels and vascular smooth muscle contractility, which together may indicate that smooth muscle activity and vascular tone are modulated in adult male depressed suicides with a history of ELA. Indeed, impaired neurovascular coupling has been experimentally determined in chronically stressed rodents (130–132), in which decreased vasomodulator enzymes neuronal nitric oxide synthase (*nNOS*) and heme oxygenase-2 (*HO2*) (131, 132) as well as malfunction of inwardly rectifying potassium (K_ir_) channels in parenchymal arteriolar myocytes were observed (130). Complimentarily, psychosocial stress rapidly increases peak latency of the hemodynamic response function in the human prefrontal cortex (133), suggesting shifted neurovascular coupling. Pathways pertaining to steroid hormone receptor activity (“Nuclear receptor transcription pathway” and “*HSP90* chaperone cycle for steroid hormone receptors SHR in the presence of ligand”) are also modulated in males, and correspond to abundant research demonstrating the role of *NR3C1* (encoding glucocorticoid receptor) (134–137) in mediating the effects of chronic stress, and the neuroprotective effects of progesterone and estrogen (138) (*PGR* and *ESR2* are upregulated in male ELA). In females, ORA and GSEA indicated downregulation of numerous, complimentary immune-related pathways, particularly those pertaining to cytokine signalling. Put succinctly, we report a global suppression of gene expression related to immune functions, encompassing both genes with pro-and anti-inflammatory roles. Our finding contradicts decades of correlative clinical evidence suggesting that elevations in circulating pro-inflammatory cytokines reflect a similarly pro-inflammatory state within the brain parenchyma in stress-related psychopathology (45, 49, 139–155). Importantly, suppressed immune signalling suggests that ELA-associated dysregulation of gene expression is not as straightforward as "peripheral inflammation equals brain inflammation". We reanalyzed a previously published RNA sequencing dataset generated in the ACC brain parenchyma (15), which is anatomically adjacent to the vmPFC, from the same female subjects. A moderately strong positive correlation was found between log2(FC) of immune-related and vascular genes within the brain parenchyma and NVU, indicating a similar pattern of downregulation for these genes (Supplementary Fig. 5).

The accuracy of our findings is bolstered by the significant downregulation of *CLDN5* expression in the female vmPFC, which corresponds to previous observations that chronic social stress alters BBB integrity through loss of the tight junction protein Cldn5 in the female mouse PFC (38), possibly by circulating cytokines (an experimentally validated mechanism: (156–158). Hence, this apparent contradiction in immune state between the blood and NVU may be reconciled by the following speculation: following ELA, the adult NVU is chronically exposed to circulating cytokines, prompting a continuous response or adaptation. Overtime, a new homeostatic point is set by the NVU. This adaptation modulates the NVU transcriptome, including the suppression of the NVU’s own immune response, which may function as a neuroprotective mechanism to shield the brain parenchyma from the deleterious effects of a pro-inflammatory milieu in the blood, as suggested by similarly downregulated immune-related genes in the ACC of the same female subjects (15). Indeed, we see possible protective mechanisms at play in females, in the form of suppressed platelet activation signalling and hemostasis pathways.

One such change in the transcriptome that demands further investigation is altered Krüppel-like Factor signalling, namely downregulated *KLF2/4* in the female NVU. Expression of *KLF2/4* acts as a central transcriptional switch point favouring a healthy, quiescent state of adult endothelial cells, while also maintaining stable expression of endothelial marker genes (86) by opening chromatin and binding to enhancer and promoter regions throughout the endothelial genome to regulate core BMEC transcriptional programs (e.g., tight junctions, adhesion, guidance cues) (159). Interestingly, *KLF2/4* impedes endothelial cell activation induced by inhibiting *TGF*β (160) and *VCAM1* (both of which are downregulated in our female data; (161) induced by diverse proinflammatory stimuli (89, 162). However, chronic pro-inflammatory stimuli may, in turn, inhibit *KLF2/4* expression (161, 163, 164). It is possible that the continuous need to compensate for peripheral inflammation eventually leads to decreased *KLF2/4* expression, reflecting an exhausted response where the system is no longer able to maintain effective levels of protection, while further losing BBB properties, as evidenced by downregulated *CLDN5* expression, which is regulated by *KLF2/4* (165, 166) (Supplementary Fig. 5). The highly significant differential expression of *KLF2/4*, coupled with their strong association with the ELA trait, and their central position within the turquoise and black modules underscores their regulatory significance in the context of ELA. Furthermore, the significant proportion of genes within the turquoise and black modules harboring *KLF2* or *KLF4* binding motifs in their distal or promoter regions (Supplementary Fig. 5) highlights the pivotal role of *KLF2/4* in modulating the BBB’s capacity to uphold brain homeostasis and safeguard the neuronal environment against stress-induced disorders.

This study has limitations that should be considered. First and foremost, the number of subjects investigated was small relative to the large number of genes tested for associations with ELA, a typical constraint in postmortem studies as well-characterized brain tissue is a limited resource. While the relatively small number of subjects included in this study limits our statistical power, we succeed in illustrating robust changes in gene regulation in cases with robust sex differences, and further validated some of the main results. It will be interesting for future studies to extend this work to additional cohorts of ELA subjects and to assess the potential associations between brain, NVU and blood gene networks, with the promise that establishing “holistic” blood to NVU to brain transcriptomic profiles might better reveal diagnostic subtypes or treatment responses. Lastly, with this study design, it is difficult to associate our findings specifically to major depression or to ELA, and a contribution of both phenotypes is likely. That ELA might drive many/most of the expression changes we identified, however, is supported by recent studies having identified NVU or immune-related contributions to ELA or chronic stress (38–44). Indeed, several gene targets identified in these investigations are found within the top gene networks identified in the present study. In conclusion, our results provide a comprehensive characterization of sex-specific transcriptional signatures in the vmPFC of depressed suicides with a history of ELA, identifying regulatory mechanisms of endothelial function that may act as potential new avenues for the development of vascular-targeted therapeutic strategies for the treatment of ELA-related psychopathology, particularly in women.

## Acknowledgements

This study was funded by an ERA NET Neuron grant and a CIHR Project grant to N.M. M.W. received scholarship support from the Réseau Québécois sur le Suicide, les Troubles de l’Humeur et les Troubles Associés (FRQ-S). The Douglas-Bell Canada Brain Bank is supported in part by platform support grants from the FRQ-S, Healthy Brains for Healthy Lives (CFREF), and Brain Canada. The present study used the services of the Molecular and Cellular Microscopy Platform (MCMP) at the Douglas Research Centre. The authors would like to thank the expert help of Douglas-Bell Canada Brain Bank staff members (J. Prud’homme, M. Bouchouka, D. Mirault, V. Lariviere, A. Baccichet) and the technology development team at the McGill University and Genome Quebec Innovation Centre.

