## Supplemental Figures for "Pervasive neurovascular dysfunction in the ventromedial prefrontal cortex of female depressed suicides with a history of childhood abuse"

**Supplementary Figures**

**Supplementary Fig. 1**

a-h) For male and female data separately, PCA was used to assess potential outliers and covariates. PCA was set up with a model including RIN, PMI, pH, age, and group status as variables. Afterwards, a DESeq2 object is generated in order to normalize the data using a median-of-ratios method and then stabilized using the vst function. A function for making PCAs was made, principal components were computed, and the variance explained by each component was calculated. Plots for the first five principal components, coloured by different variables (RIN, PMI, pH, and age) to visualize how the variable is distributed across PCs. For each variable, linear models were constructed to correlate principal components with the variables. For both males and females, no subjects were deemed outliers and, thus, none were removed. Variables with correlations padj < 0.05 were deemed significant and were added to the design formula required by DEseq2 when performing differential gene expression analysis. For males, age (PC3: padj = 0.007; r^2^ = 0.23) and pH (PC2: padj = 0.00043; r^2^ = 0.36) were identified as covariates and, for females, age (PC2: padj = 0.036; r^2^ = 0.30) was identified as a covariate. i) Output from SVA and variance partitioning to assess whether incorporating SVA within DGE analysis is appropriate. This approach can inform whether the surrogate variables identified by SVA are significant enough to warrant their inclusion in the DESeq2 design formula, ensuring that we are not only controlling for known confounding variables but also for unknown factors that could affect the expression levels. The first plot, prior to SVA, shows a larger proportion of variance categorized as "Residuals," indicating that the known variables do not account for much of the variance in the expression data. In the second plot, after SVA, the surrogate variables (SV1 to SV6) capture a significant amount of variance. This suggests that there are substantial effects in the data that are not accounted for by the measured variables like pH, PMI, RIN, and age. The residuals are noticeably smaller when surrogate variables are included, which indicates that SVA is capturing unwanted variation in the expression data, warranting the inclusion of SVA within DGE analysis to improve the detection of truly differentially expressed genes by reducing background noise from unmeasured confounders.

a) Males: RIN

| 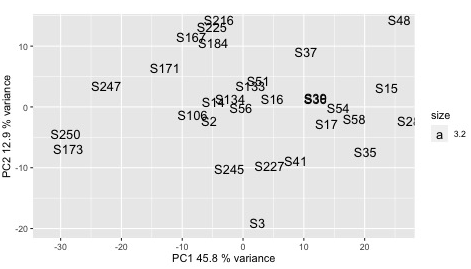 | 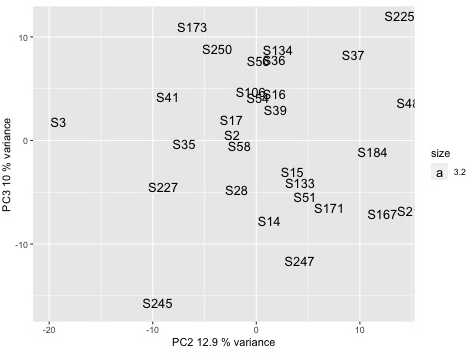 |
| --- | --- |
| 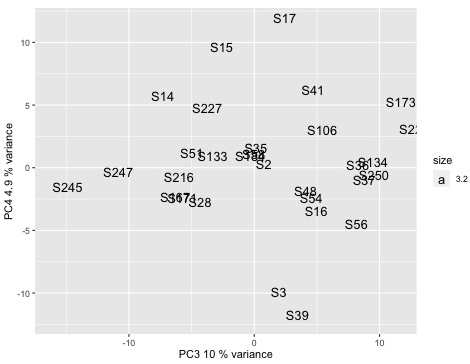 | 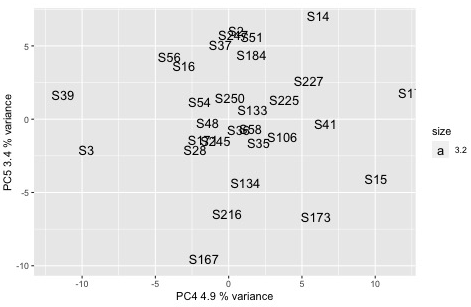 |

b) Males: PMI

| 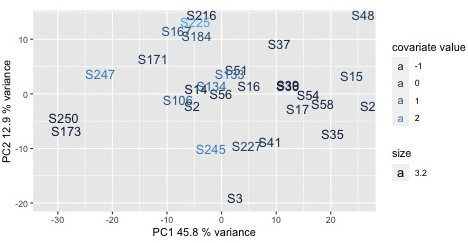 | 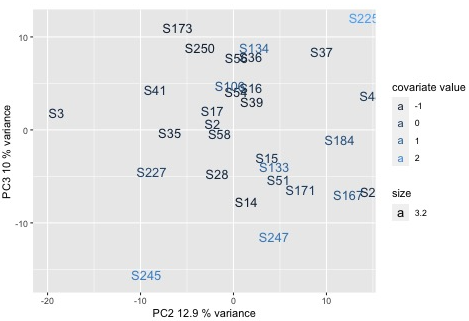 |
| --- | --- |
| 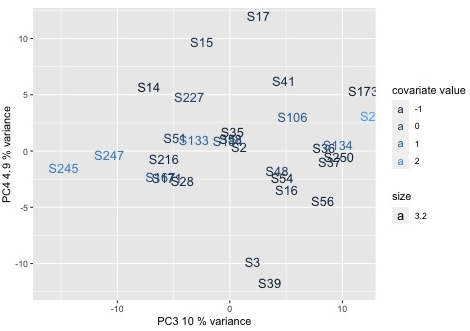 | 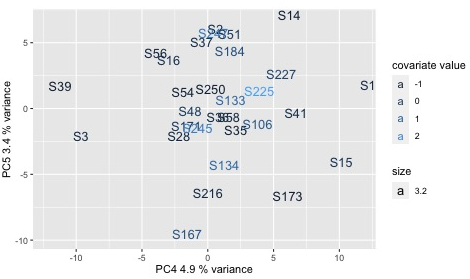 |

**c) Males: pH**

| 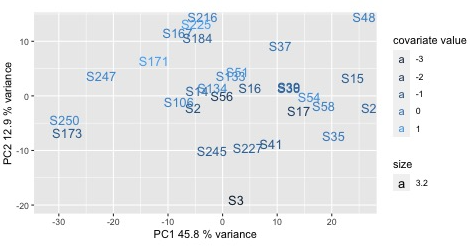 | 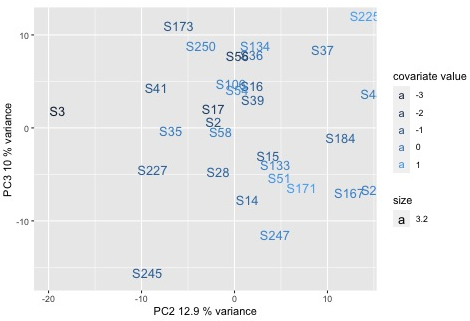 |
| --- | --- |
| 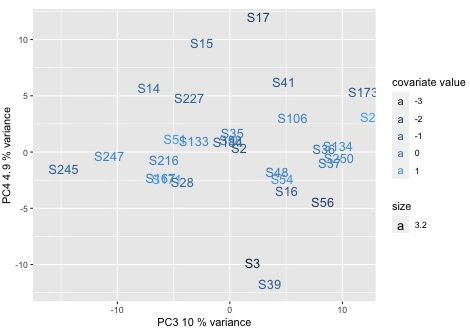 | 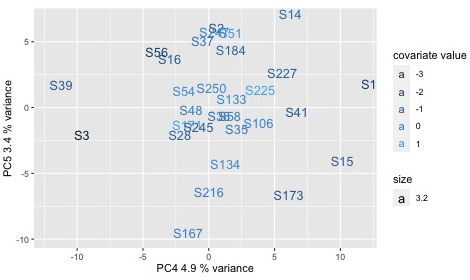 |

**d) Males: Age**

| 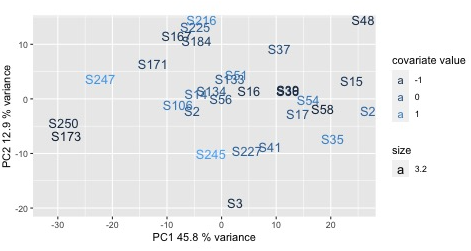 | 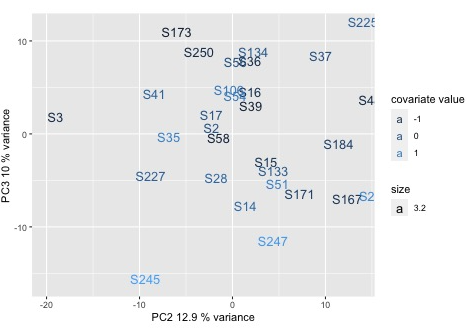 |
| --- | --- |
| 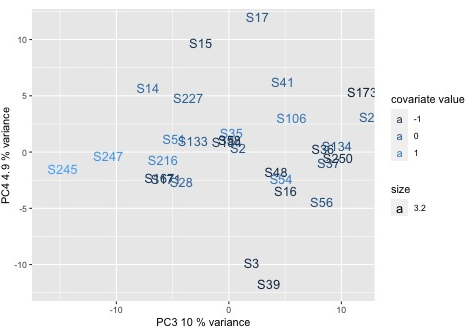 | 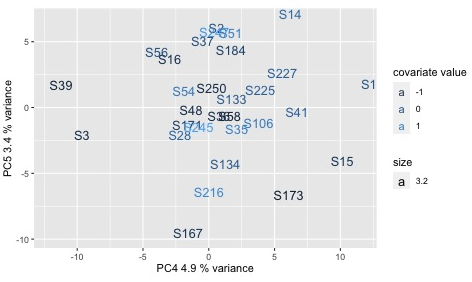 |

e) Females: RIN

| 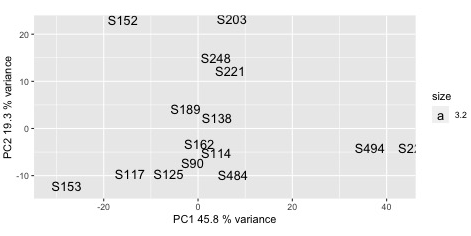 | 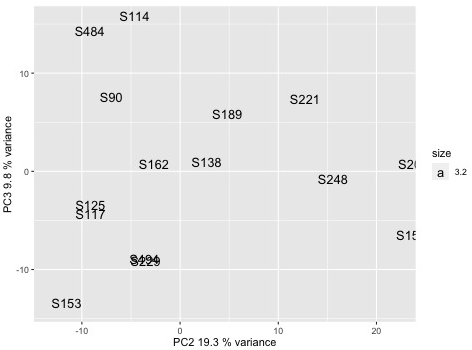 |
| --- | --- |
| 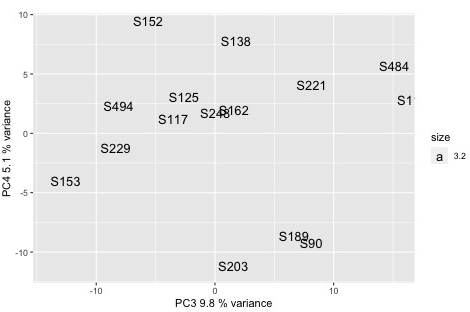 | 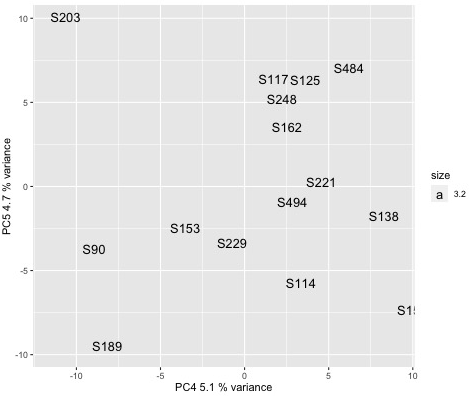 |

f) Females: PMI

| 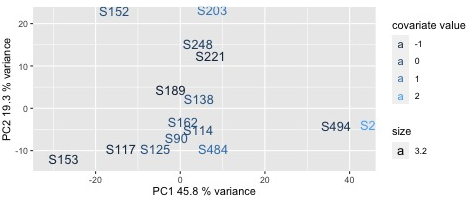 | 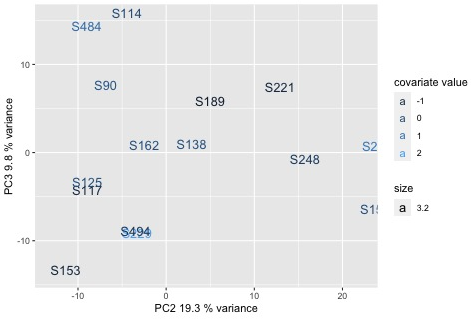 |
| --- | --- |
| 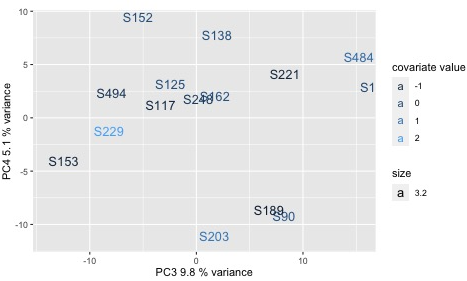 | 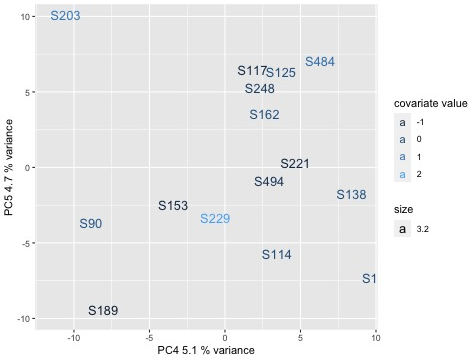 |

g) Females: pH

| 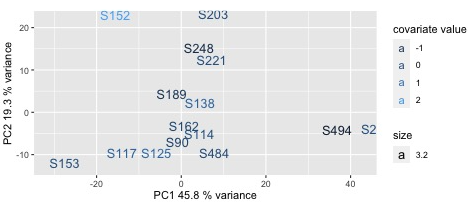 | 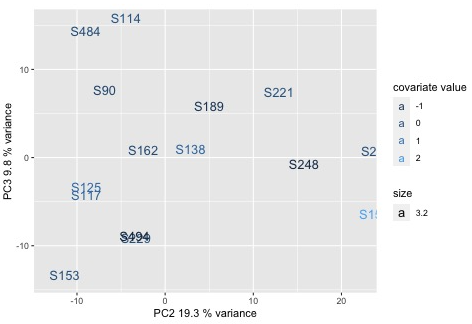 |
| --- | --- |
| 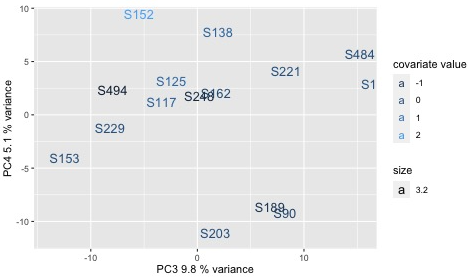 | 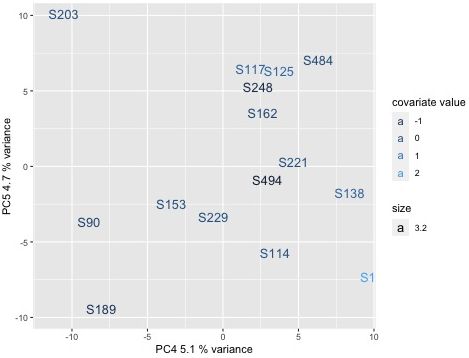 |

h) Females: Age

| 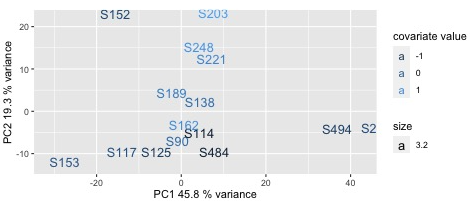 | 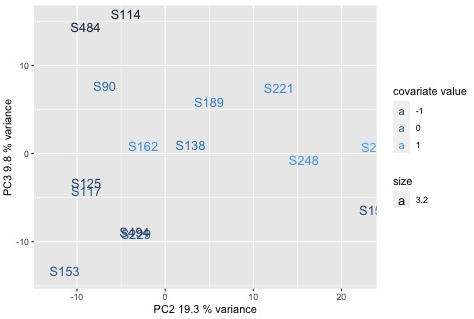 |
| --- | --- |
| 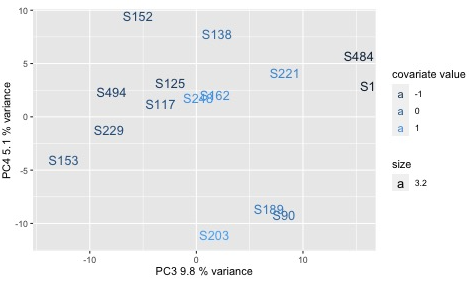 | 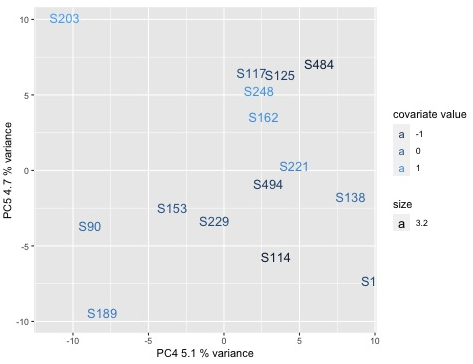 |

i)

| 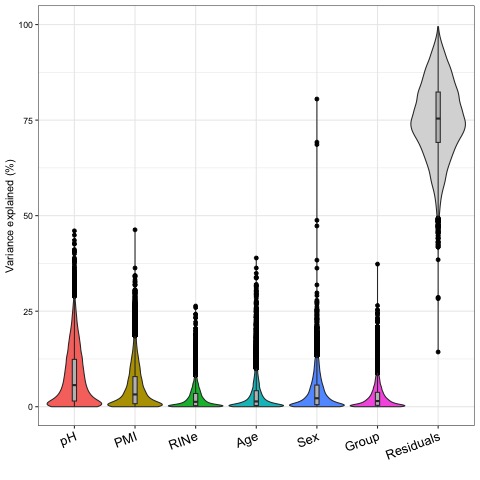 | 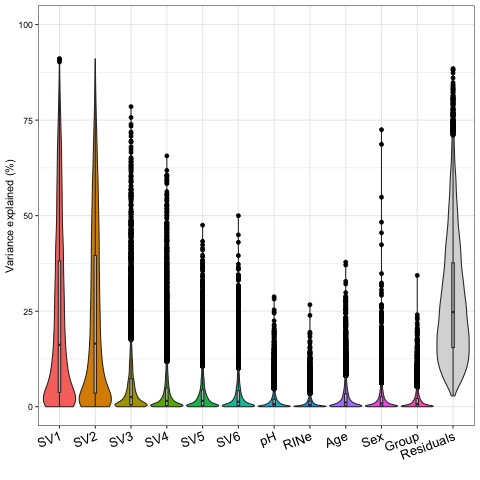 |
| --- | --- |

**Supplementary Fig. 2**

a) Select plots for representative vascular markers. b) Table summarizes p-values from statistical tests comparing cell type proportions (percentage) between CTRL and ELA groups, to determine if there are statistically significant differences. The proportions were taken from the results for bioinformatic deconvolution using CIBERSORTX, evaluating target cell types: 1) sum of all vascular genes, 2) endothelial genes, 3) mural and perivascular fibroblast genes, as well as contaminating cell types 4) neuronal genes and 5) all genes from contaminating cell types. The results indicate that there were no significant differences across all comparisons, except for a marginally significant result in the CTRL vs ELA comparison among female subjects. Prior to analysis, Shapiro-Wilk tests for normality and Levene's tests for equality of variances were conducted to ensure that the data met the assumptions required for each statistical test applied. Where the normality assumption was not met, the Mann-Whitney test was utilized instead of the two-sample t-test. All Vascular = arterial genes + capillary genes + mural cell genes + perivascular fibroblast genes + astrocyte genes + T cell genes. All Contamination = oligodendrocyte precursor cell genes + oligodendrocyte genes + neuron genes. We identified a total of 463 DEGs, comprising 281 upregulated and 182 downregulated genes associated with a history of ELA irrespective of sex. c) Select DEGs from the initial pooled-sex DGE analysis, identifying DEGs irrespective of sex. This figure illustrates DEGs previously implicated in the literature, underscoring the value of analyzing larger, combined cohorts for enhanced statistical power. Refer to Supplementary Table 4 for a complete list of DEGs from the initial pooled-sex DGE analysis. d) Representative violin plots showing sex-driven expression differences in DEGs identified in sex-pooled subjects with a history of ELA compared to sex-pooled controls.

a)

| 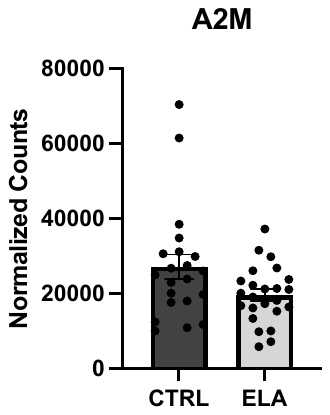 | 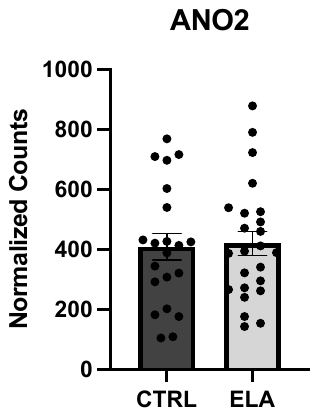 | 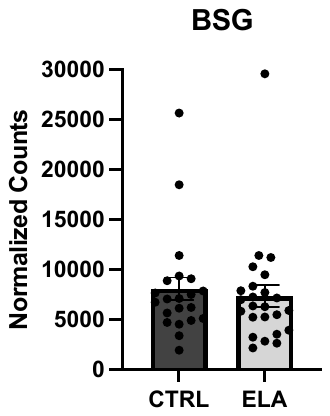 | 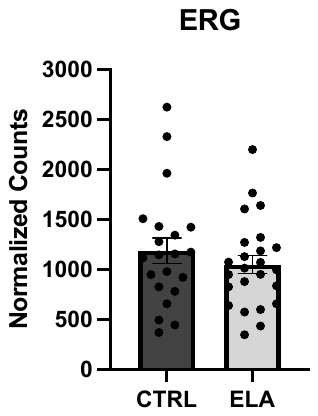 | 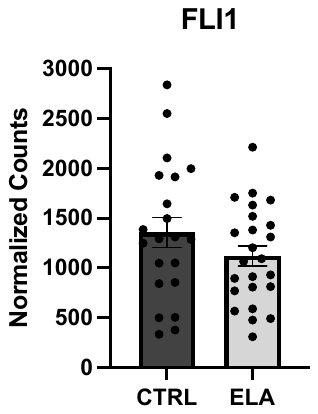 | 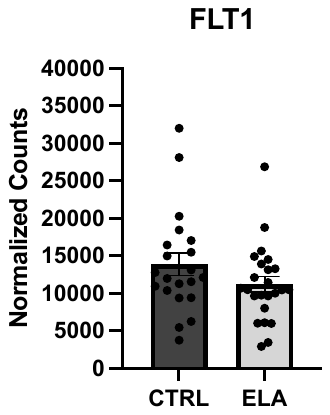 |
| --- | --- | --- | --- | --- | --- |
| 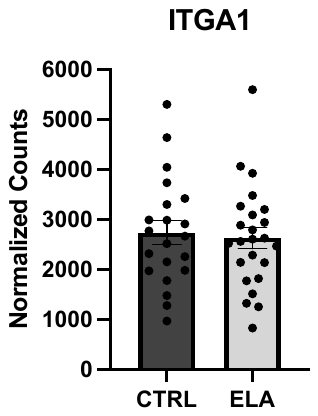 | 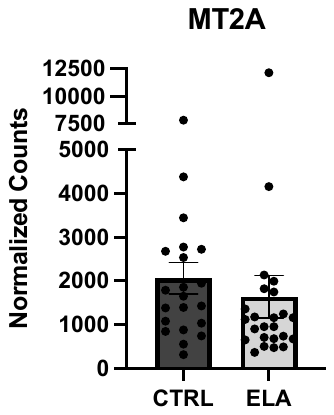 | 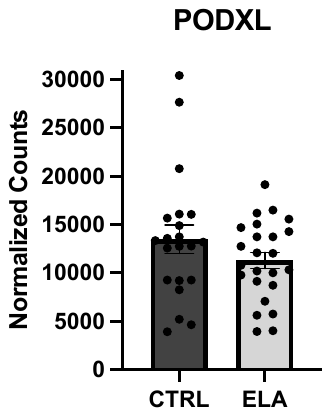 | 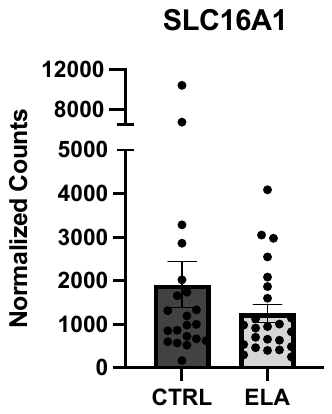 | 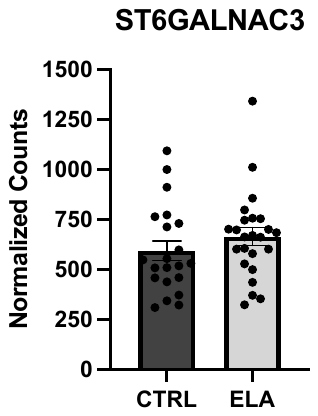 | 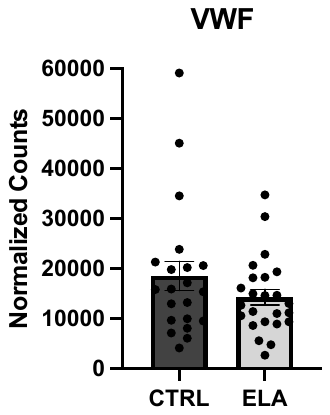 |
| 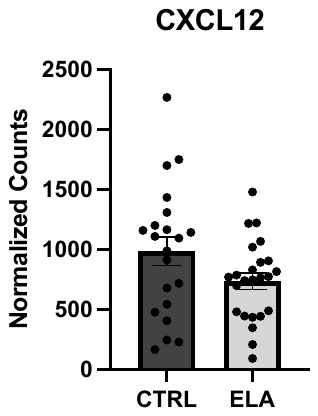 | 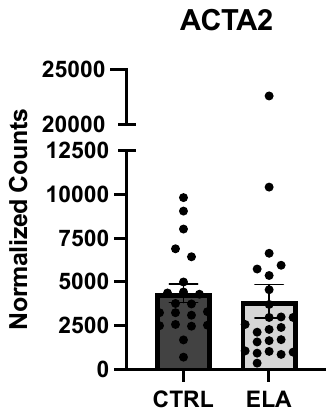 | 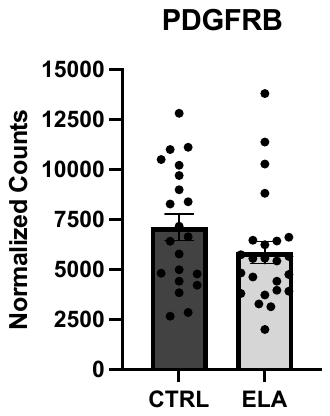 | 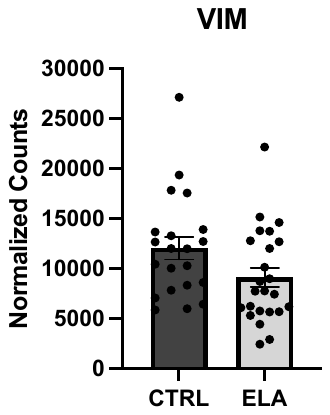 | 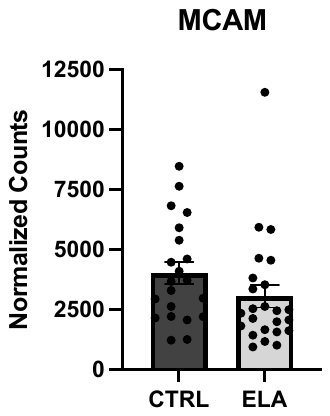 | 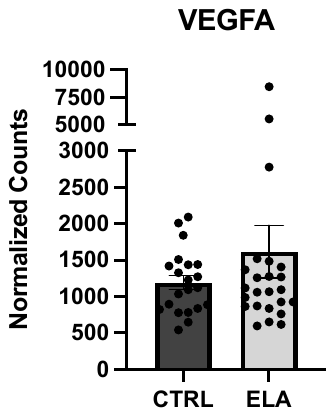 |
| 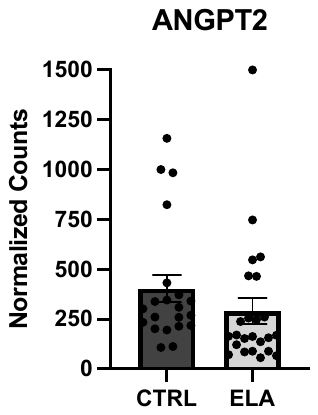 | 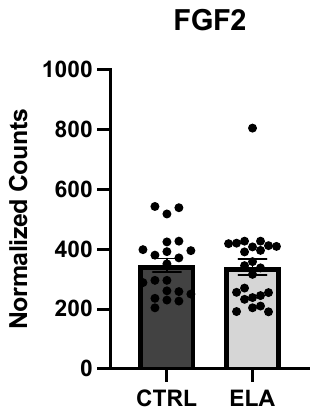 | 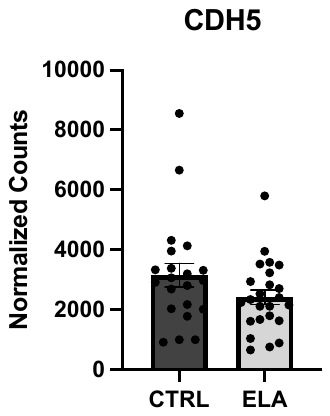 | 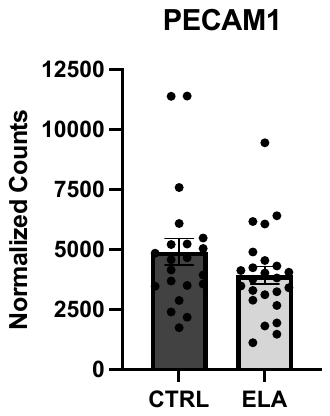 | 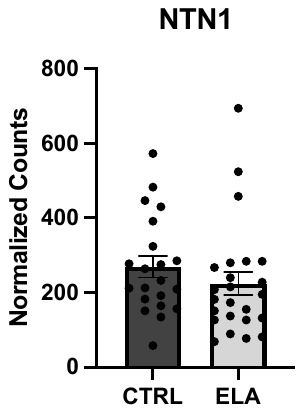 | 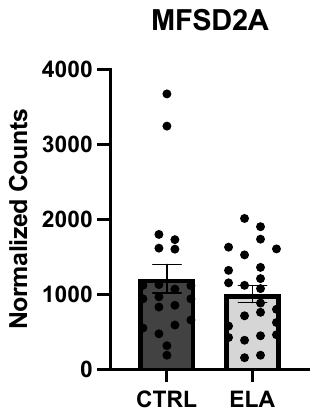 |
| 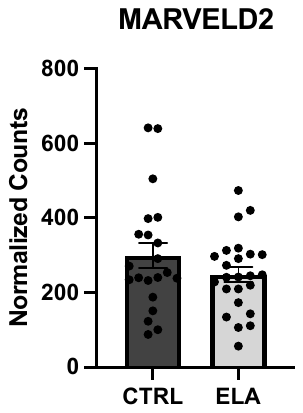 | 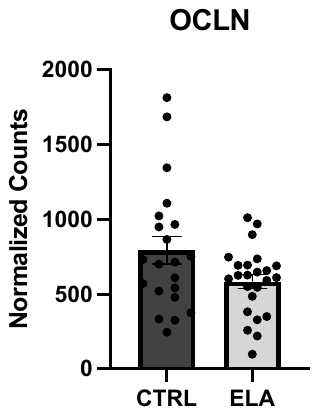 | 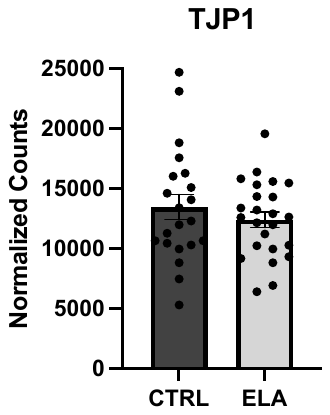 | 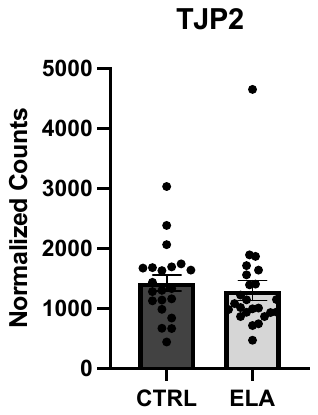 |  |  |

b)

| **Category (%)** | **CTRL vs ELA (sexes pooled)** | **CTRL vs ELA (Male)** | **CTRL vs ELA (Female)** |
| --- | --- | --- | --- |
| **All Vascular** | 0.337 | 0.8213 | 0.04073 |
| **Endothelial** | 0.991 | 0.3753 | 0.3357 |
| **Mural + P. Fibro** | 0.3629 | 0.5128 | 0.7046 |
| **Neuron** | 0.2506 | 0.9343 | 0.09975 |
| **All Contamination** | 0.3812 | 0.6799 | 0.05286 |

c)

| 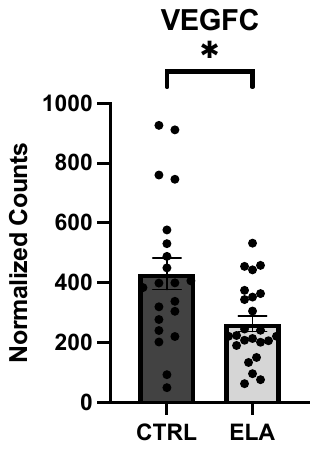 | 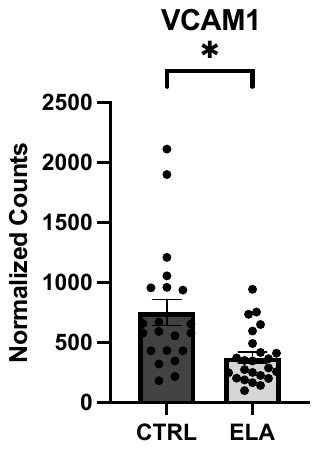 | 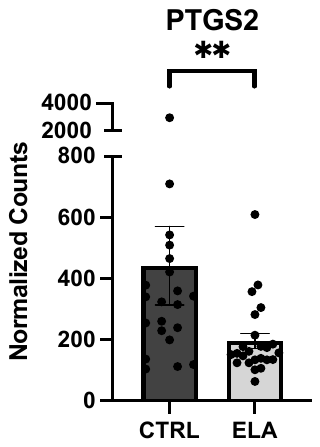 | 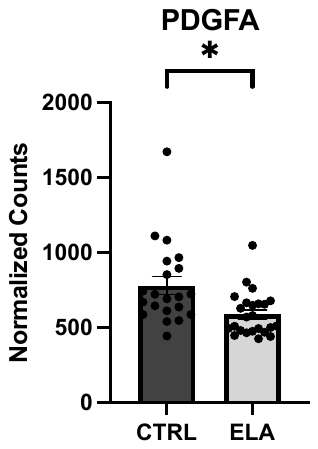 |
| --- | --- | --- | --- |
| 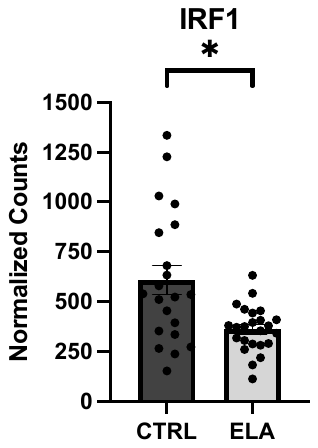 | 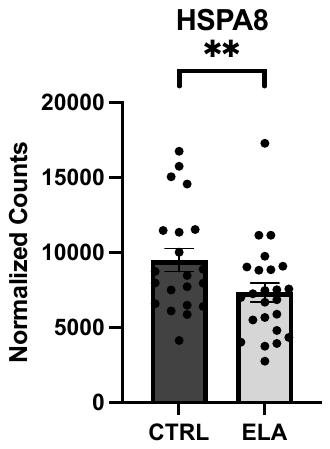 | 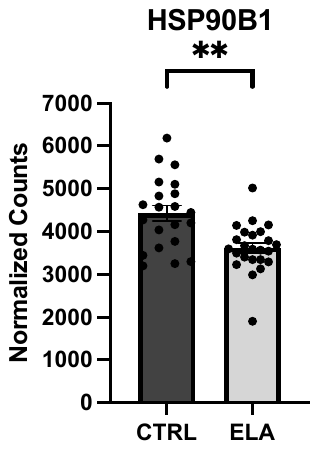 | 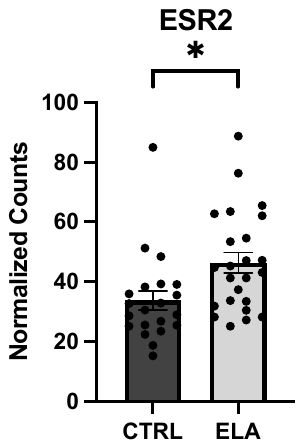 |
| 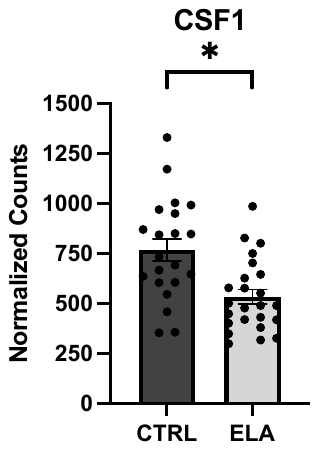 | 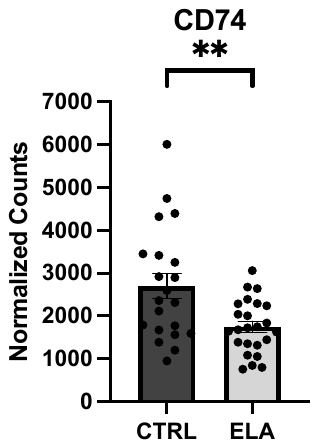 | 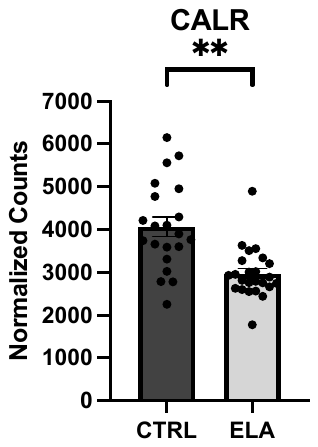 | 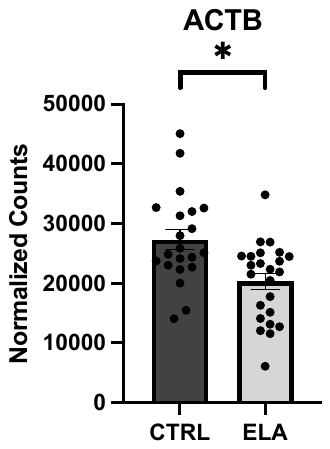 |

d)

| 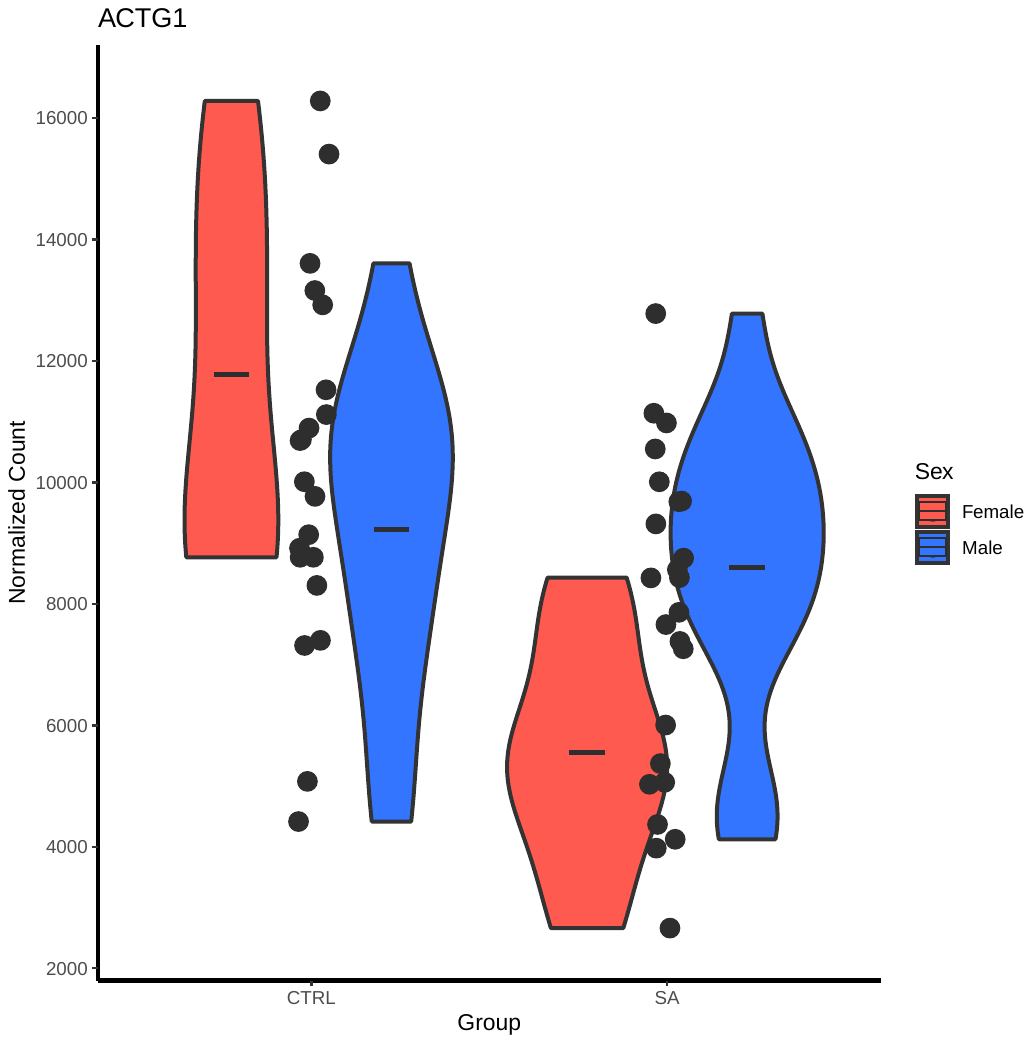  CTRL ELA | 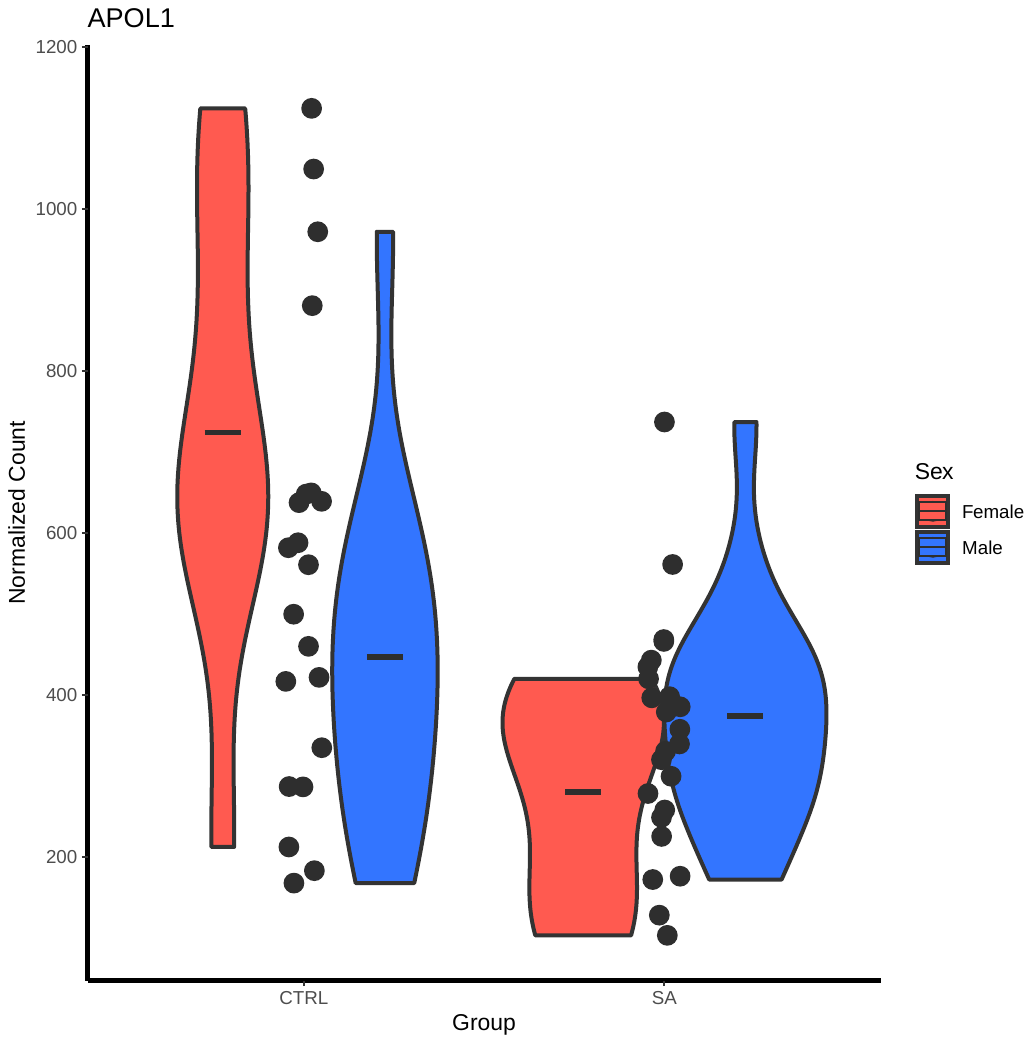  CTRL ELA | 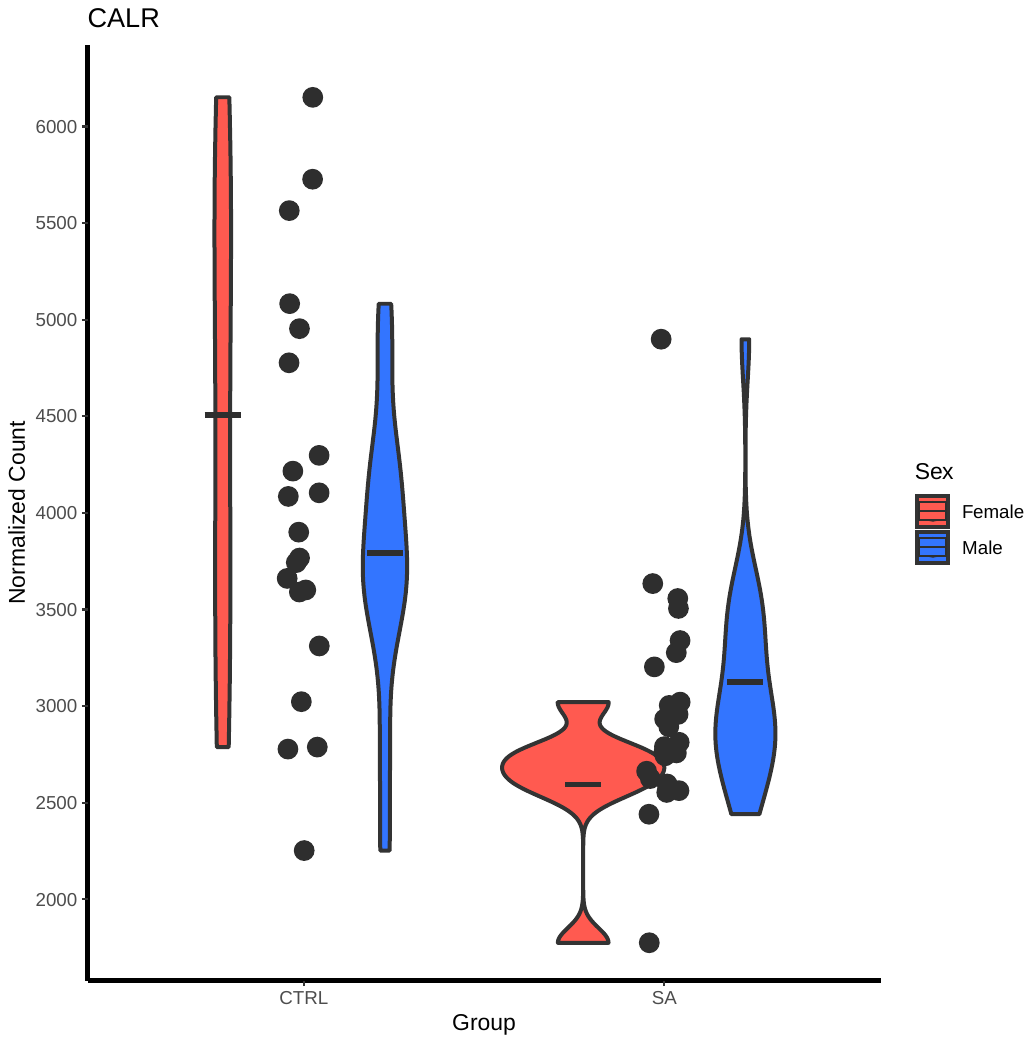 | 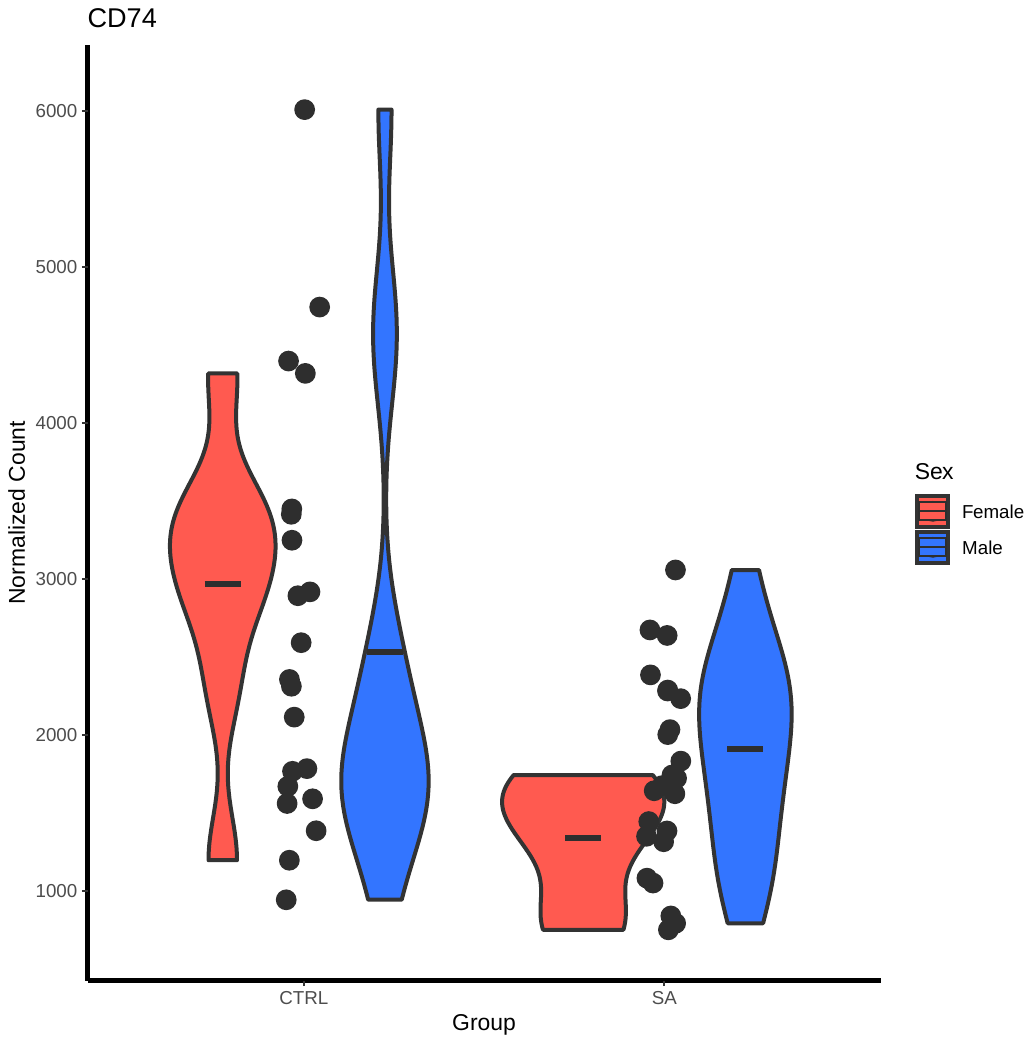 |
| --- | --- | --- | --- |
| 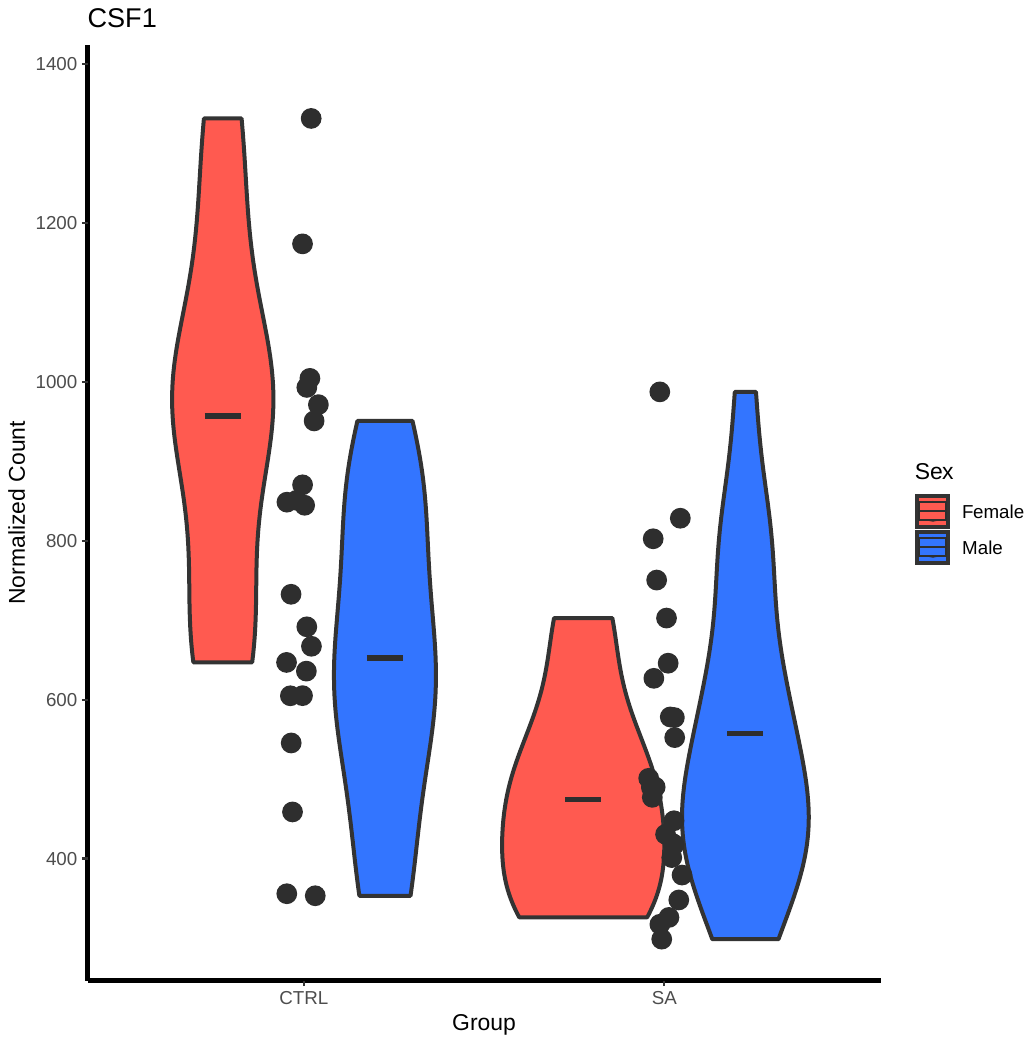  CTRL ELA | 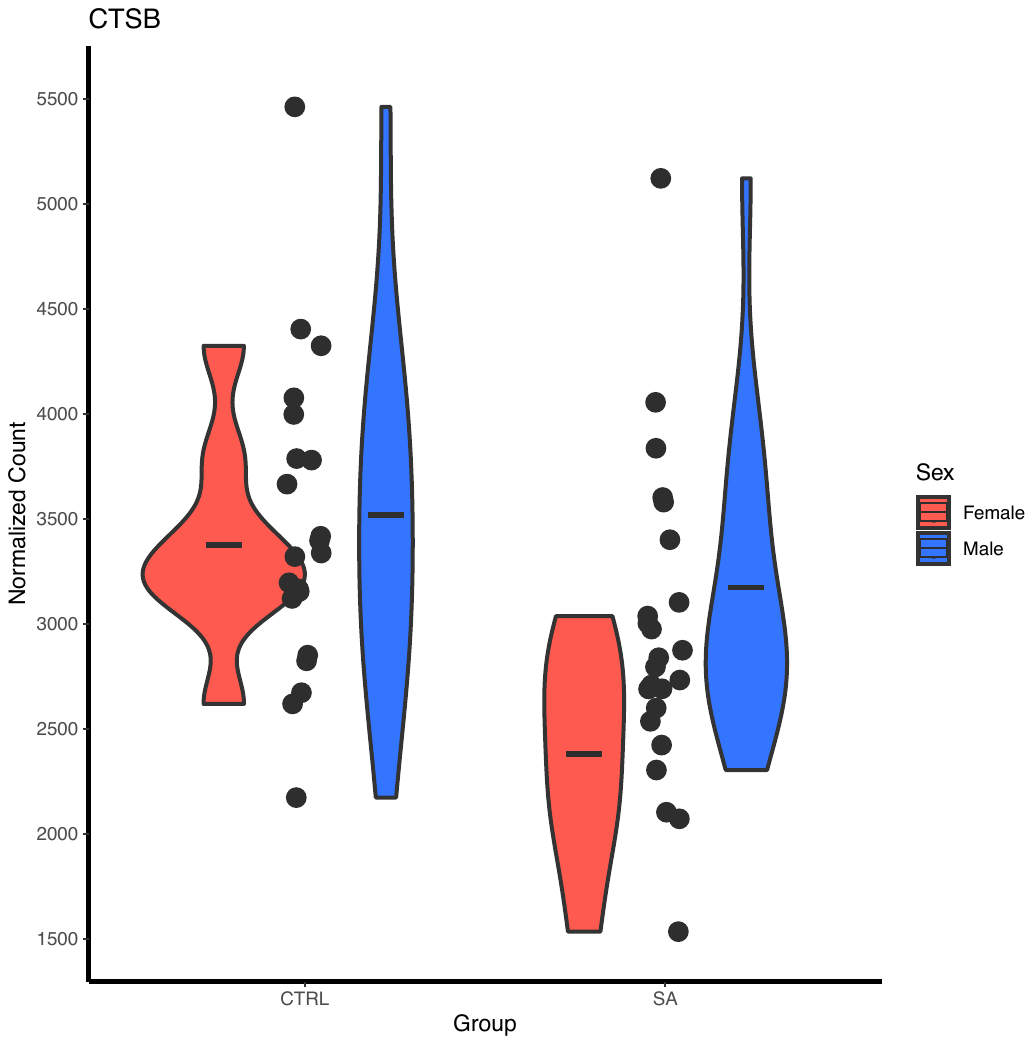  CTRL ELA | 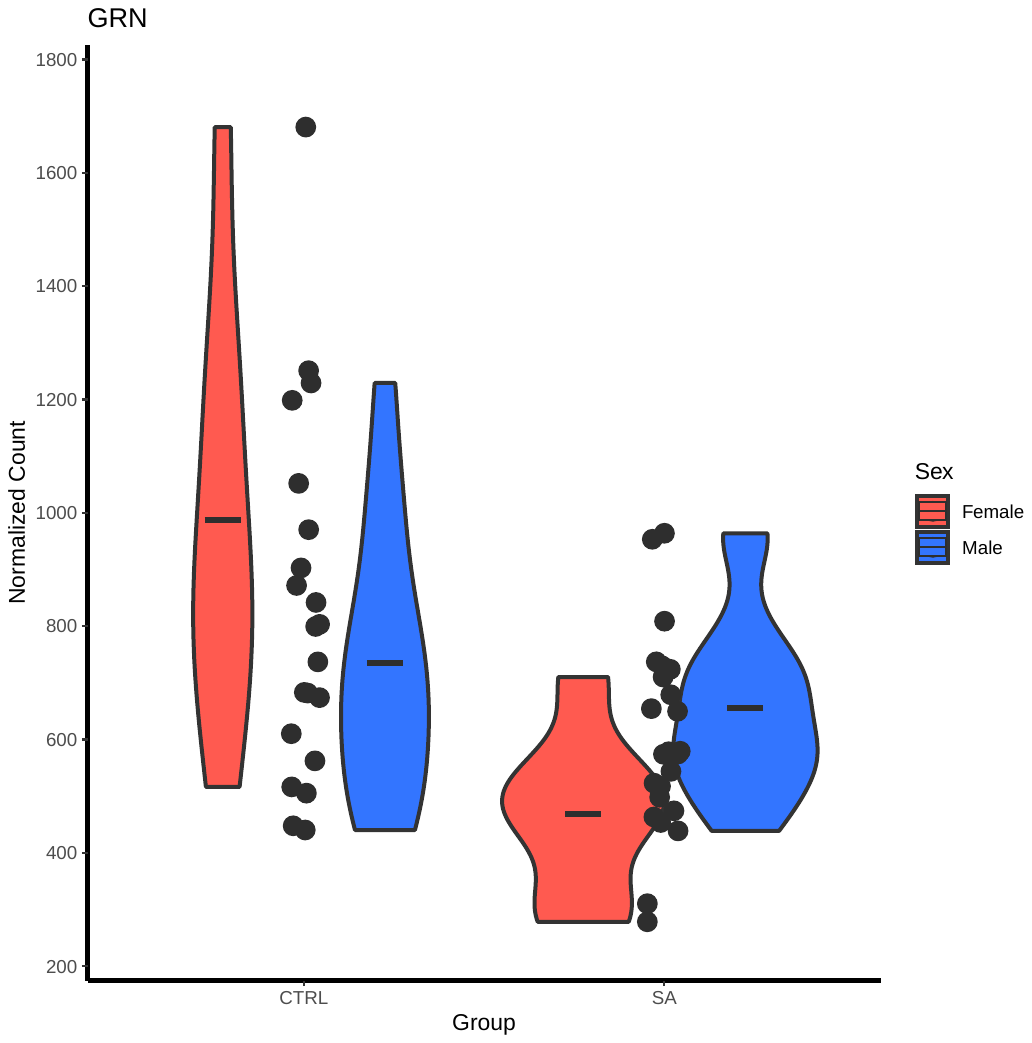  CTRL ELA  CTRL ELA | 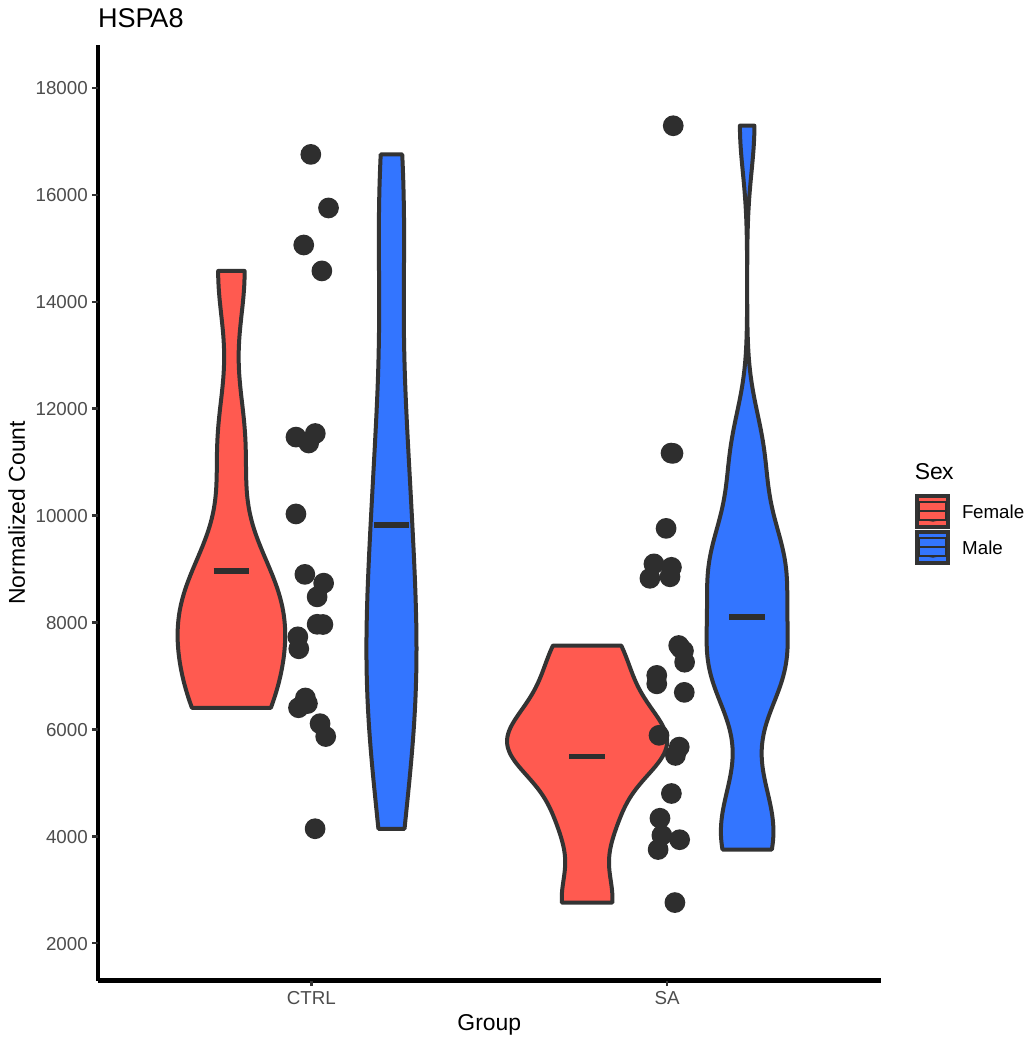  CTRL ELA  CTRL ELA |
| 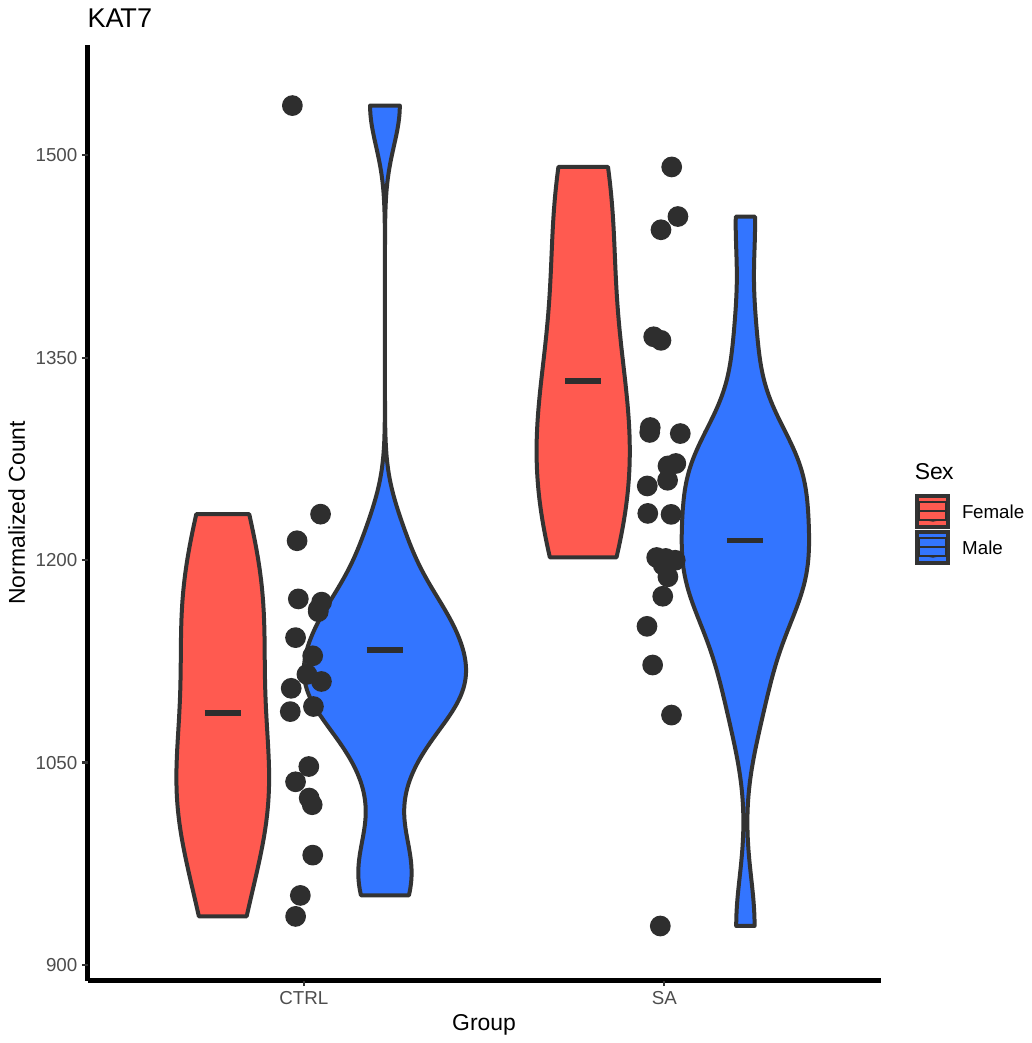  CTRL ELA | 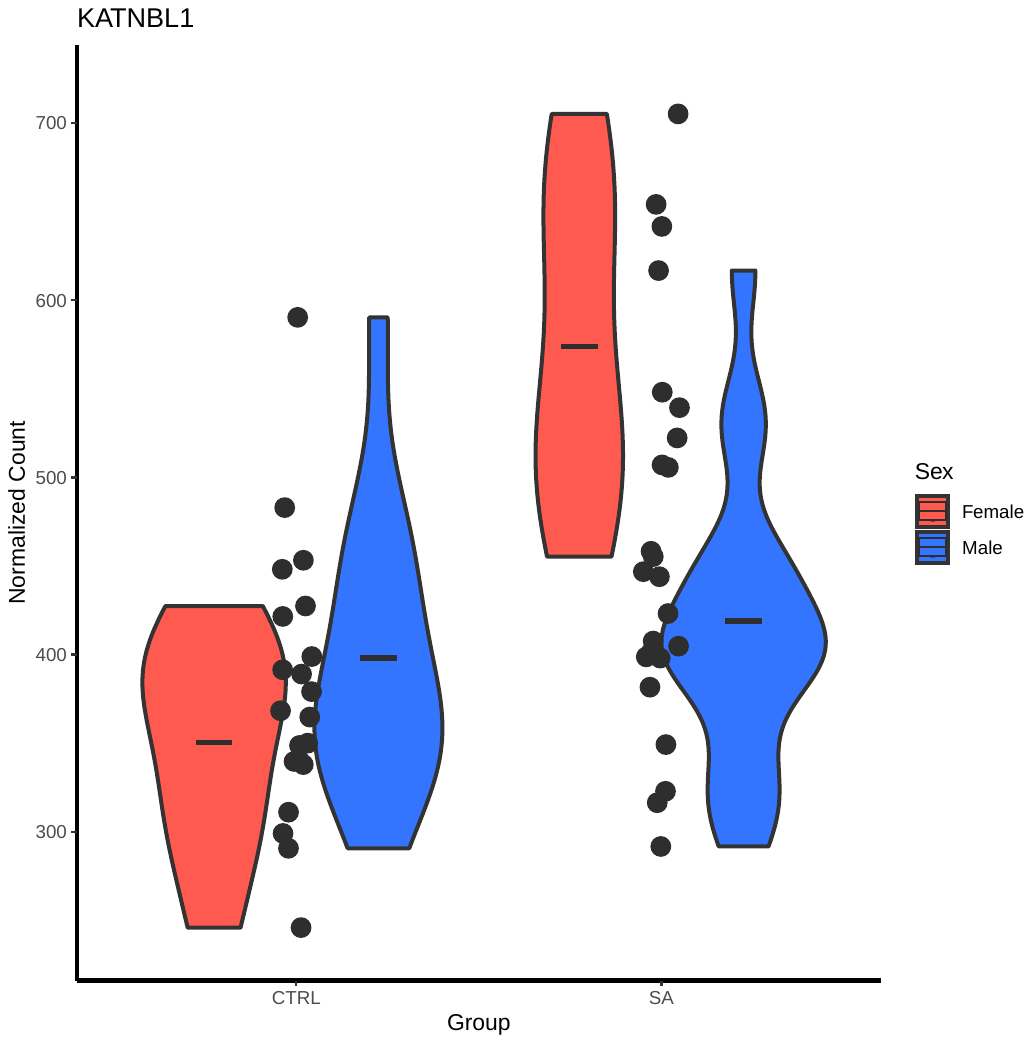  CTRL ELA | 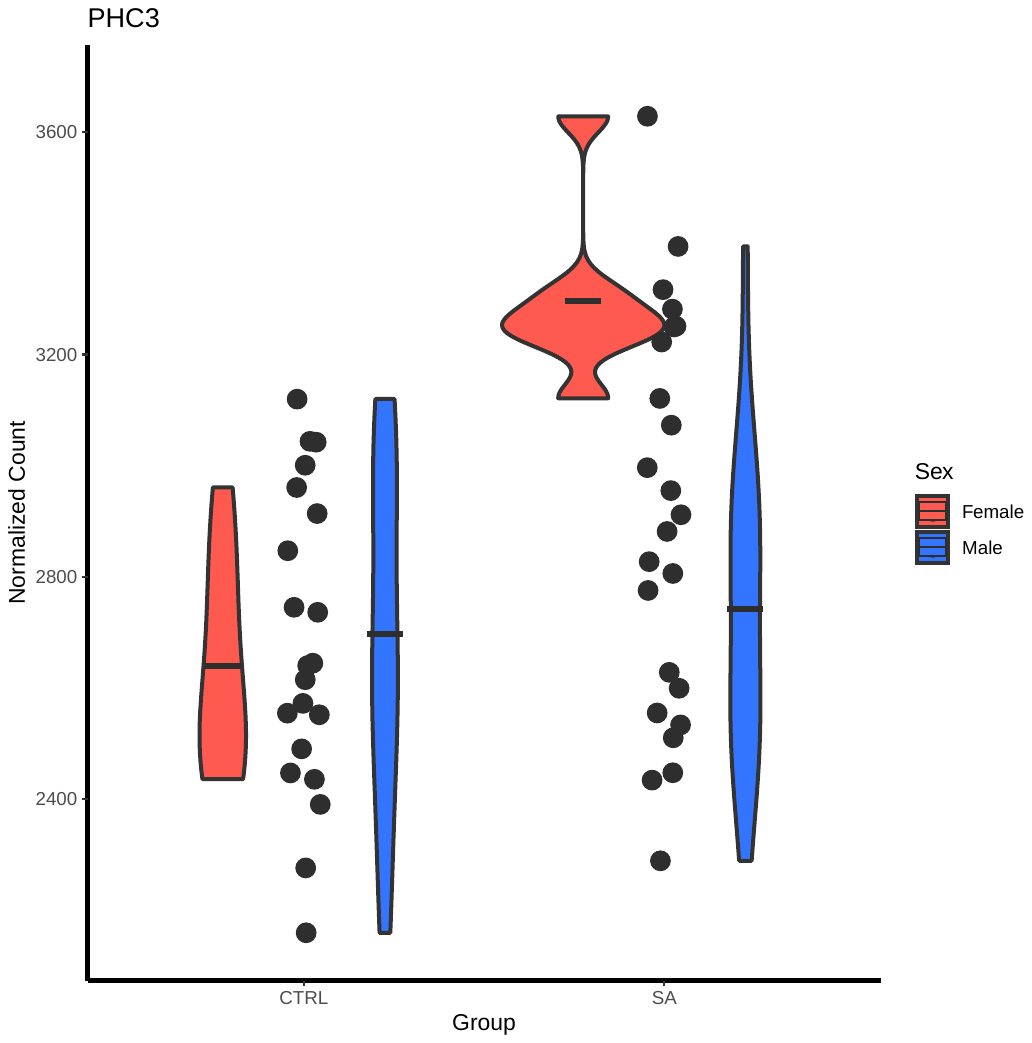  CTRL ELA | 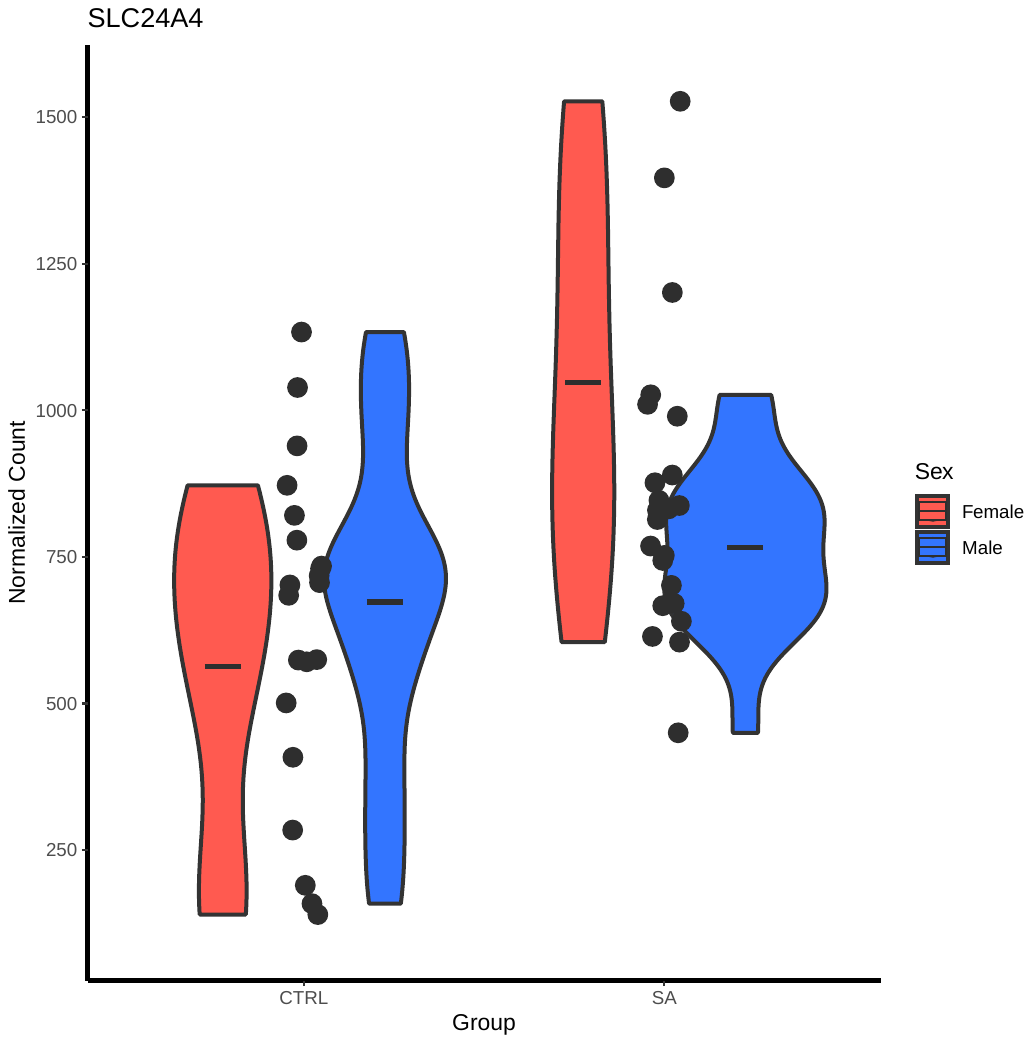  CTRL ELA |
| 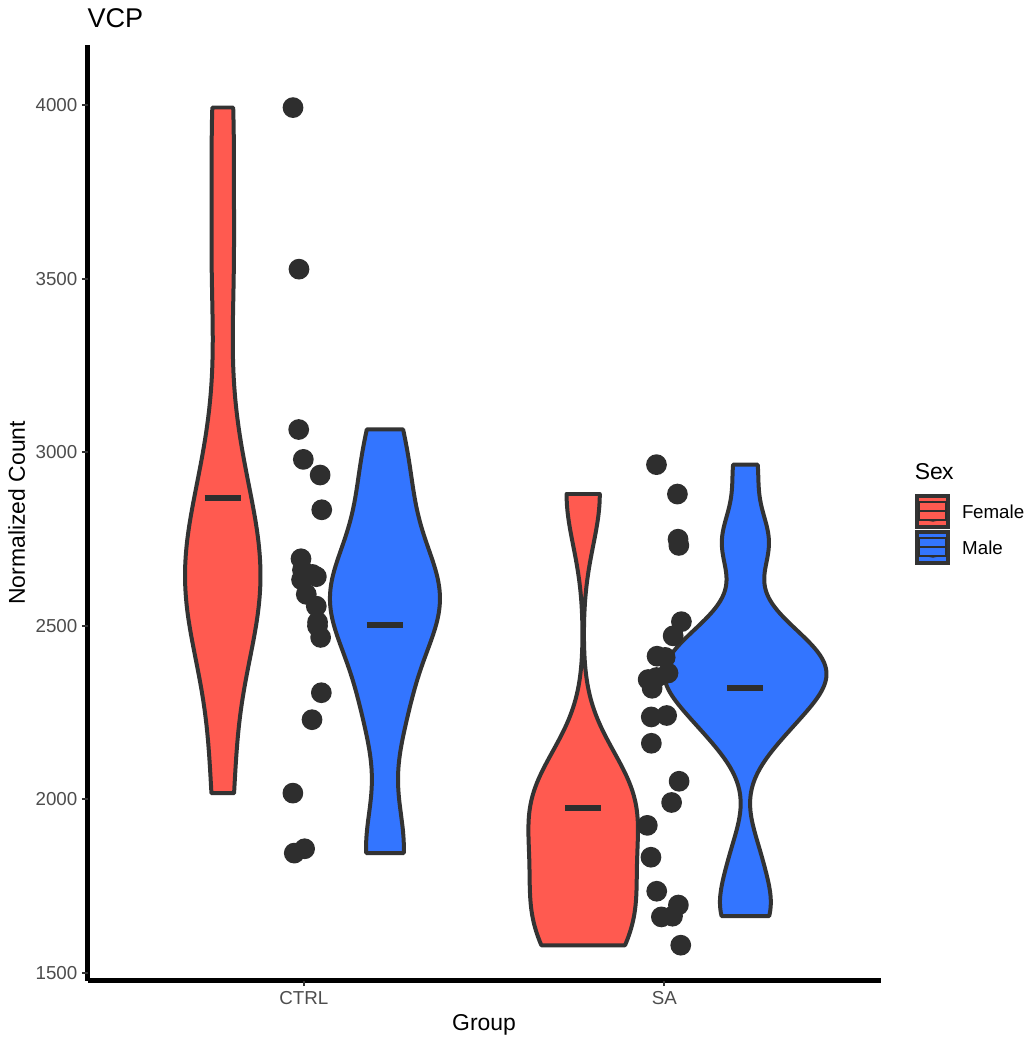  CTRL ELA | 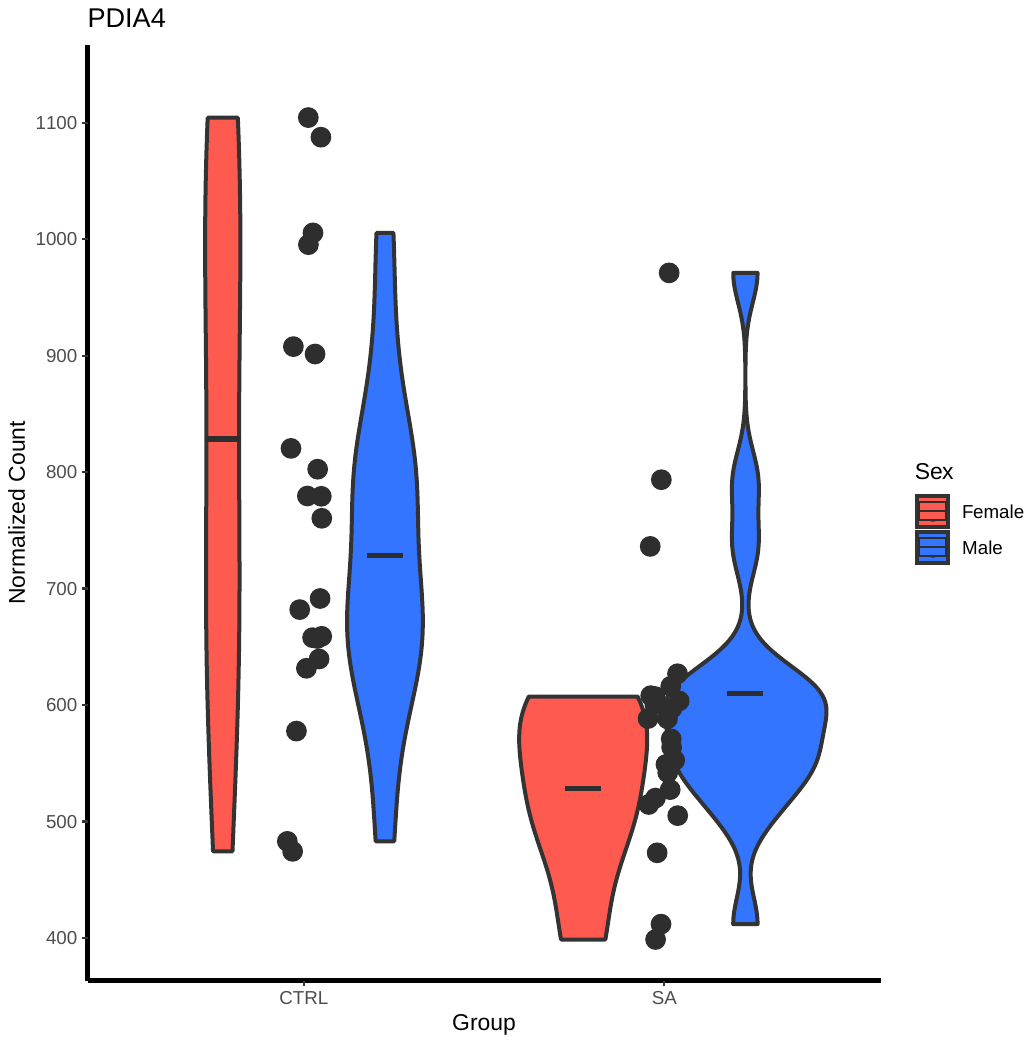  CTRL ELA | 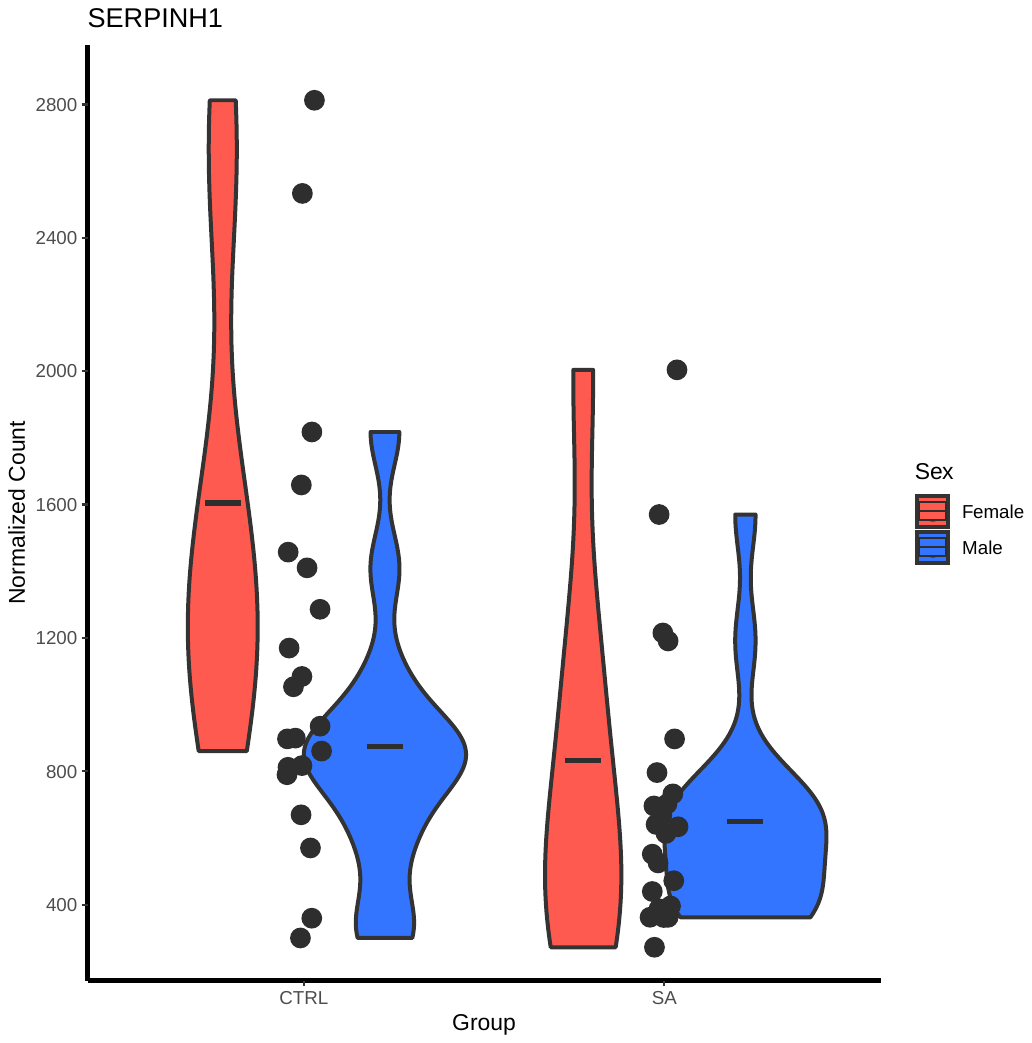  CTRL ELA | 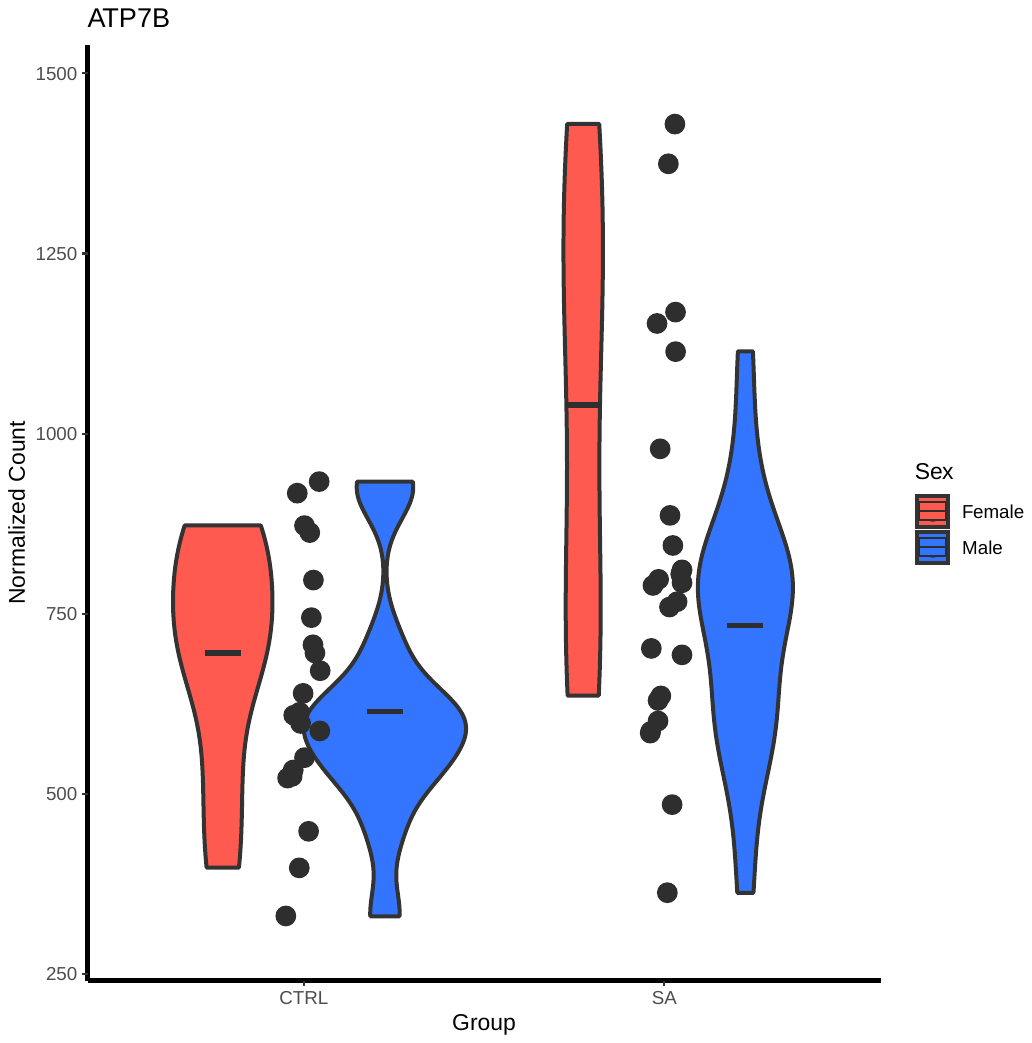  CTRL ELA |

A correlation analysis between principal components and sex revealed significant, moderate associations for specific components – namely PC1 (p=0.0082, r²=0.152), PC3 (p=0.03, r²=0.104), and PC5 (p=0.0067, r²=0.159) – suggesting sex contributes to variance in our data. Additionally, Welch Two Sample t-test was conducted to determine whether the mean values of PC1 (which attributes to 35.4% of variation) are significantly different between males (mean=3.776618) and females (mean=-7.553237), yielding t(27.245)=2.7398, p-value=0.01072, with 95% confidence interval [2.848402 19.811307], suggesting that there are differences in the expression patterns captured by PC1 between the two sexes. Similarly, the mean values of PC5 are significantly different between males (mean=1.293569) and females (mean=-2.587138), yielding t(22.219)=2.5907, p-value=0.01661, with 95% confidence interval [0.7759907 6.9854226]. Further, Welch Two Sample t-tests comparing the mean values of principal components between sexes – particularly for PC1 and PC5 – indicated significant differences in expression patterns between males and females. Specifically, PC1 (which accounts for 35.4% of variation) showed a mean difference (mean in males=3.776618; mean in females=-7.553237) with a t-value of 2.7398 (p=0.01072, 95% confidence interval [2.848402, 19.811307]), and PC5 also demonstrated a significant mean difference (mean in males=1.293569; mean in females=-2.587138; t=2.5907, p=0.01661, 95% confidence interval [0.7759907, 6.9854226]).

**Supplementary Fig. 3**

The DESeq function was used to explore the Sex:Group interaction, looking specifically at the interaction between the factors "Sex" and "Group" as a means to assess whether the effect of ELA is different between males and females. Intriguingly, many DEGs identified by the Sex:Group interaction exhibit sexual divergence in their differential expression (representative DEGs are presented in Supplementary Fig. 3), where 66.4% of DEGs demonstrate that the effect of ELA is more pronounced in females compared to males.

a) Representative plots of gene differential expression by sex in ELA, identified by DGE analysis including the Sex:Group interaction. This figure highlights sexual dimorphism in gene expression specific to ELA, illustrating distinct patterns with notable changes primarily observed in females. b) Boxplots representing the distribution of Cook's distances for each female subject across genes. The DESeq function calculates, for every gene and for every subject, a diagnostic test for outliers named Cook’s distance*.* Cook’s distance is a measure of how a subject is influencing the fitted coefficients for a gene. In addition to constructing PCA plots to find potential subject outliers (as performed above), boxplots of Cook’s distances can also be used to identify subjects with median values very different from other subjects. No female subjects deviate significantly from one another, suggesting there are no outlier subjects driving the expression differences observed. DEGs identified in the male CTRL vs. male ELA comparison, and DEGs identified in female CTRL vs. female ELA comparison were respectively cross-referenced with vascular cell type defining markers from recent studies. c) Matrix displaying the overlap of male and female DEGs from our study with vascular cell-type markers defined by Yang et al. (2022). d) Matrix displaying the overlap of male and female DEGs from our study with vascular cell-type markers defined by Garcia et al. (2022). DEGs were cross-referenced against both previously published datasets due to their different strategies in defining vascular-cell type defining markers.

a)

| 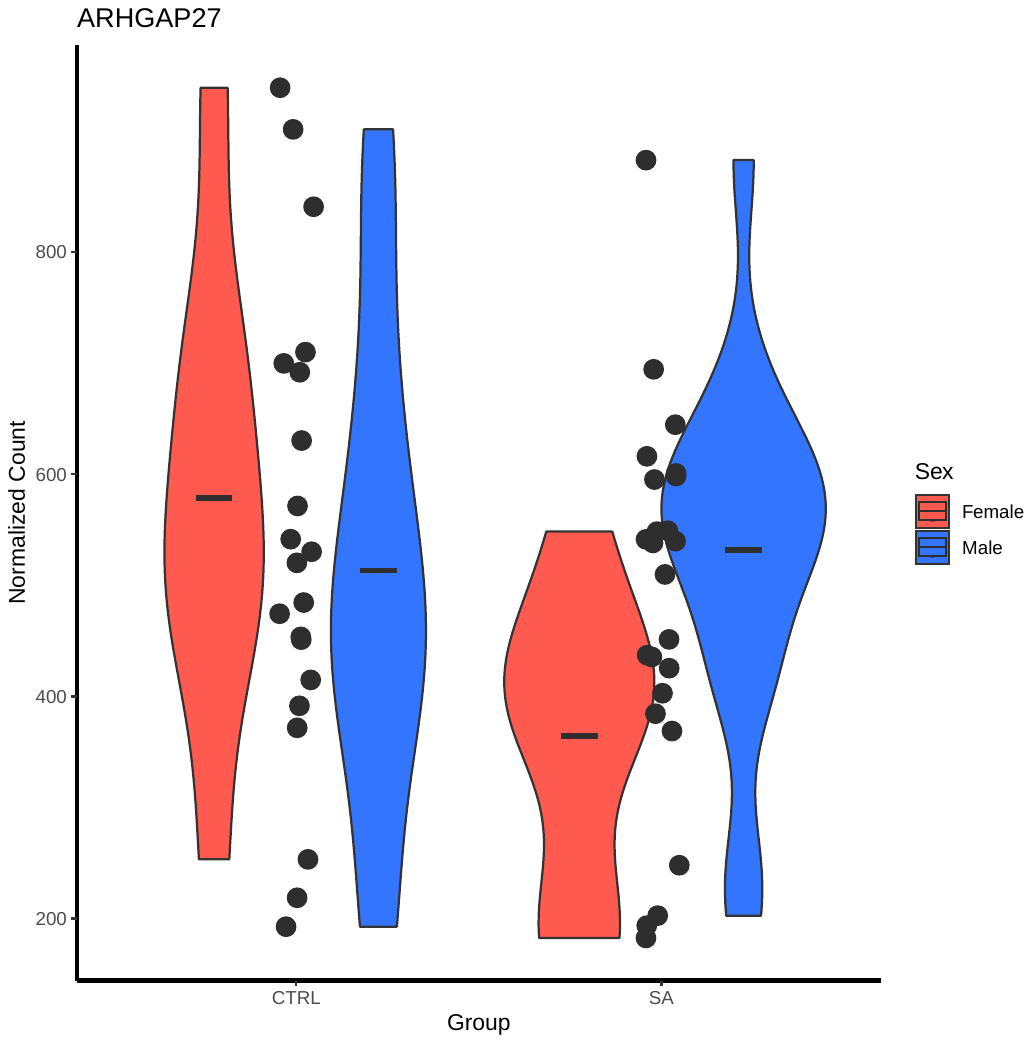  CTRL ELA | 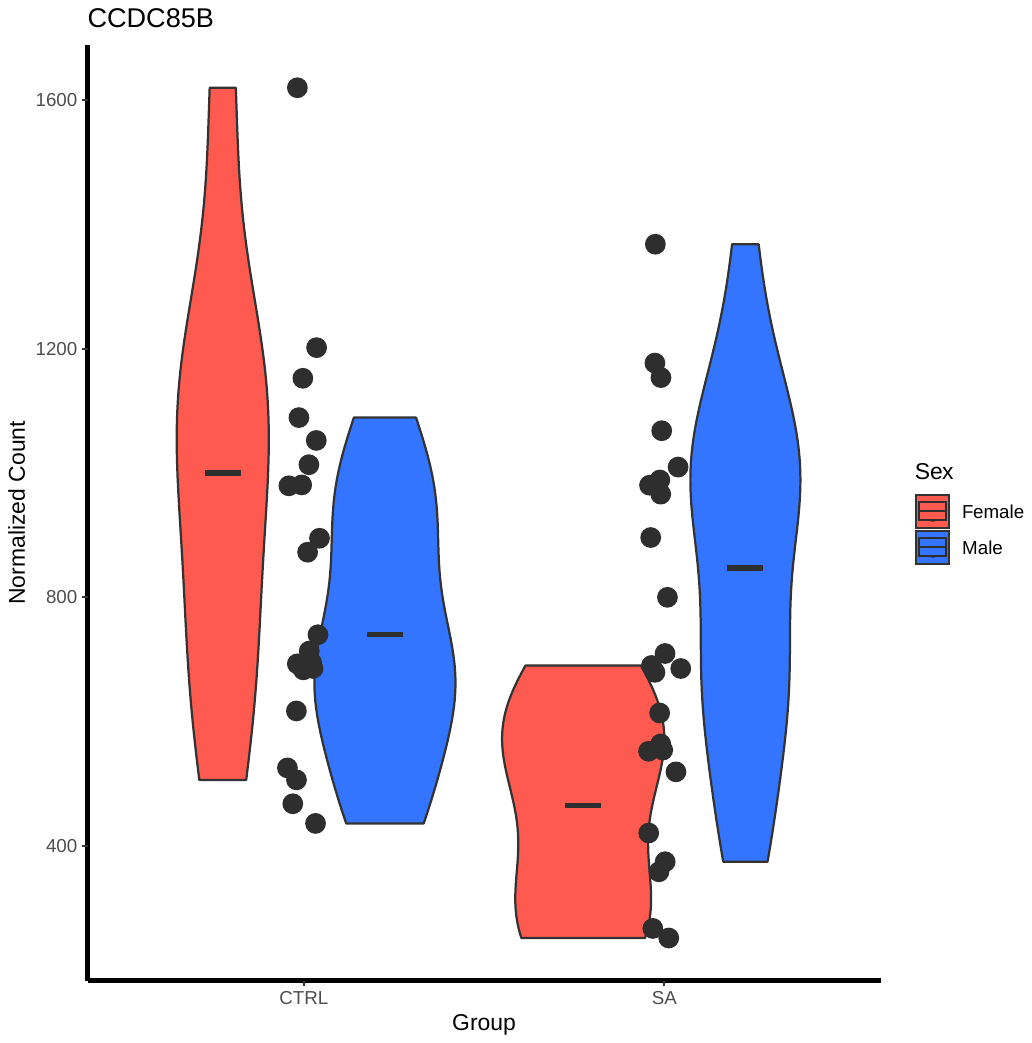  CTRL ELA | 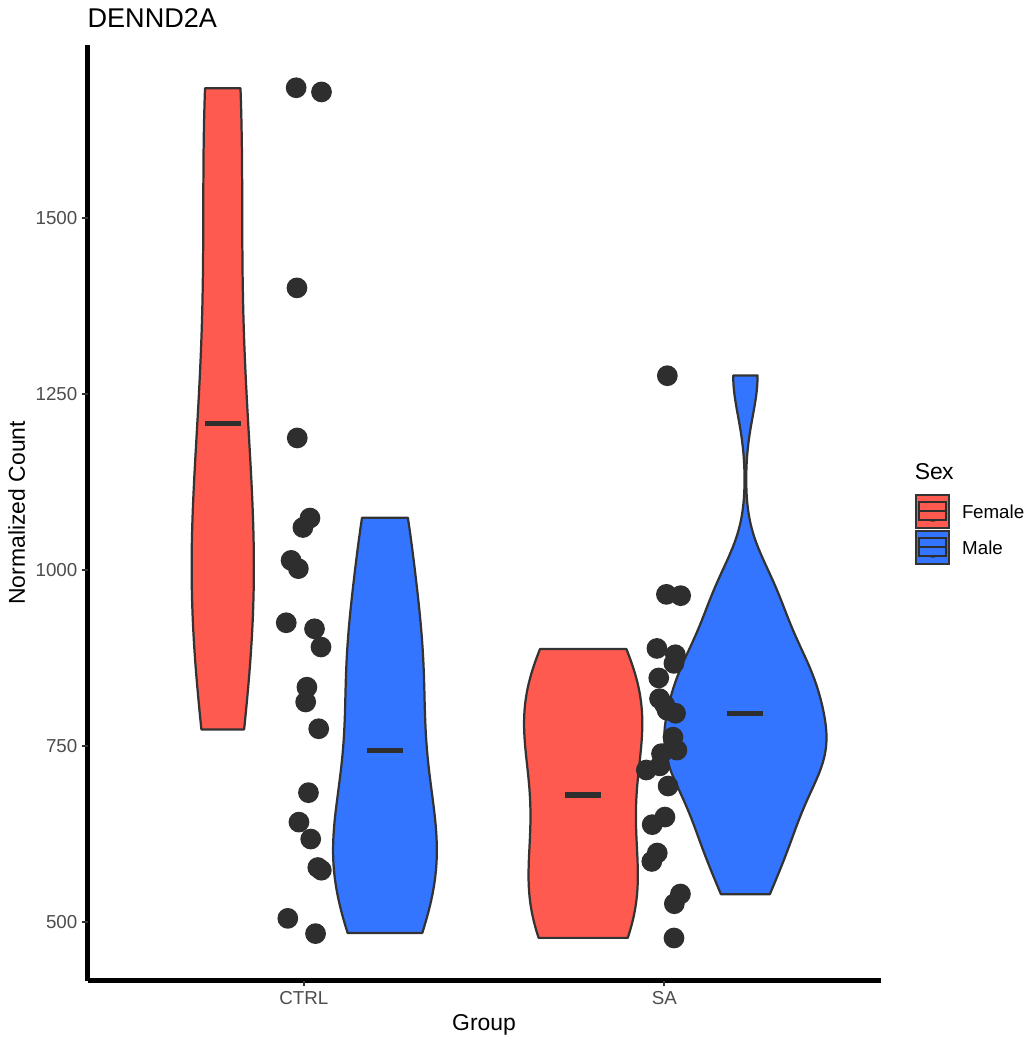  CTRL ELA | 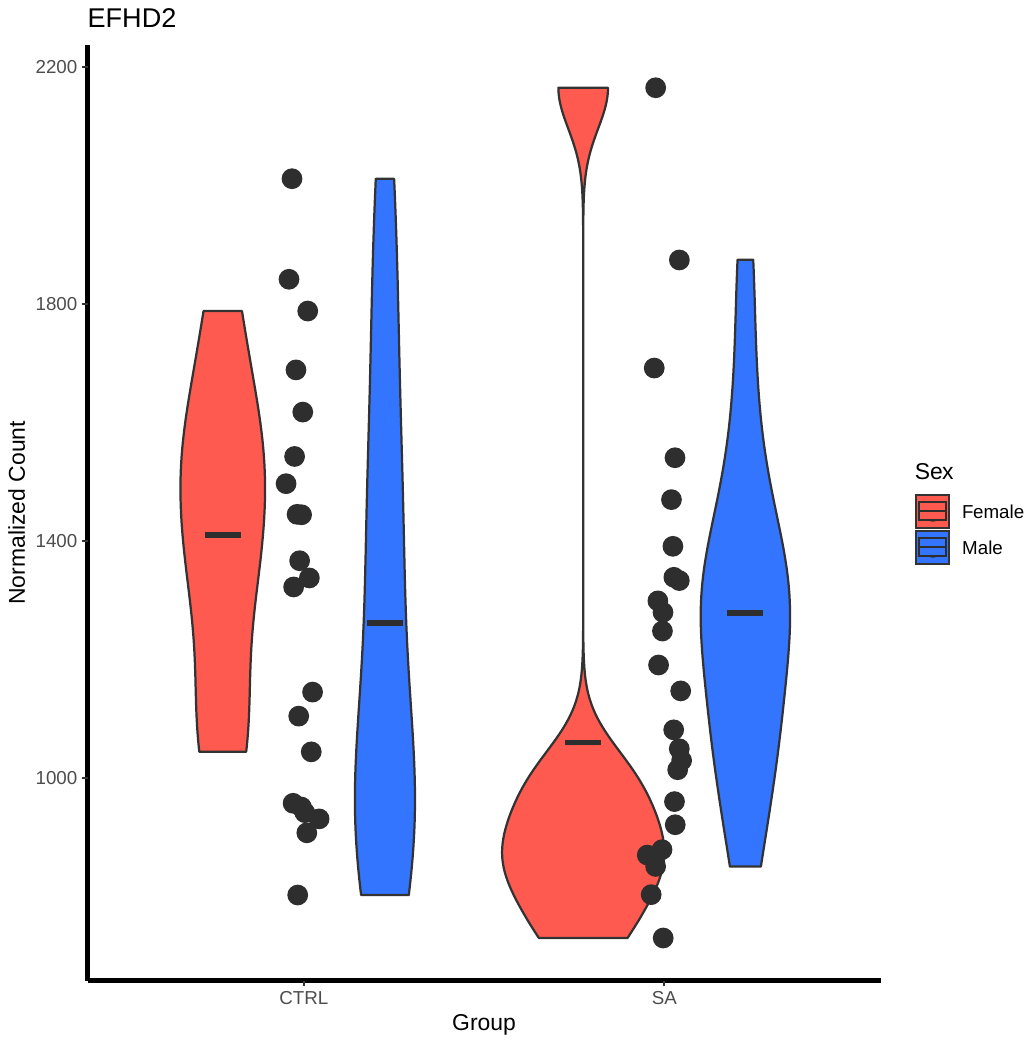 |
| --- | --- | --- | --- |
| 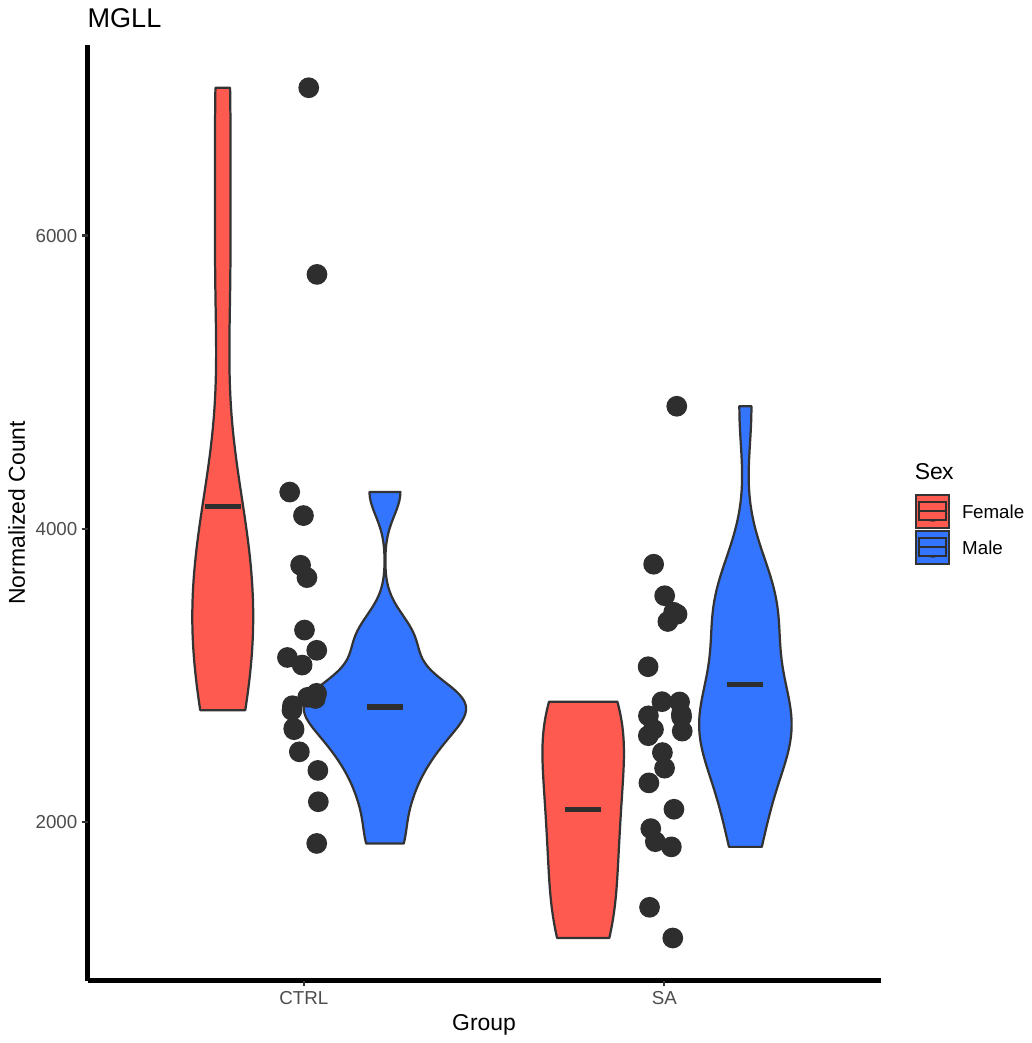  CTRL ELA | 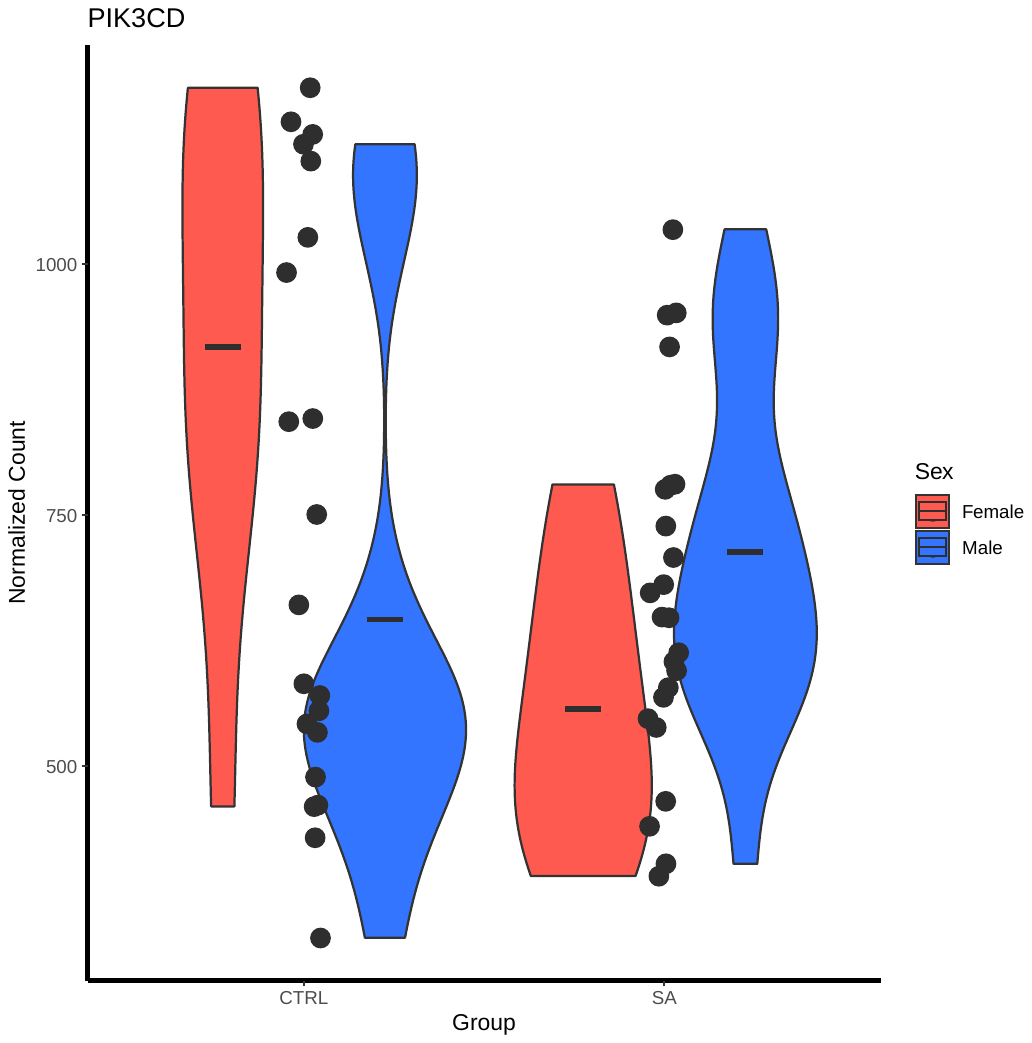  CTRL ELA | 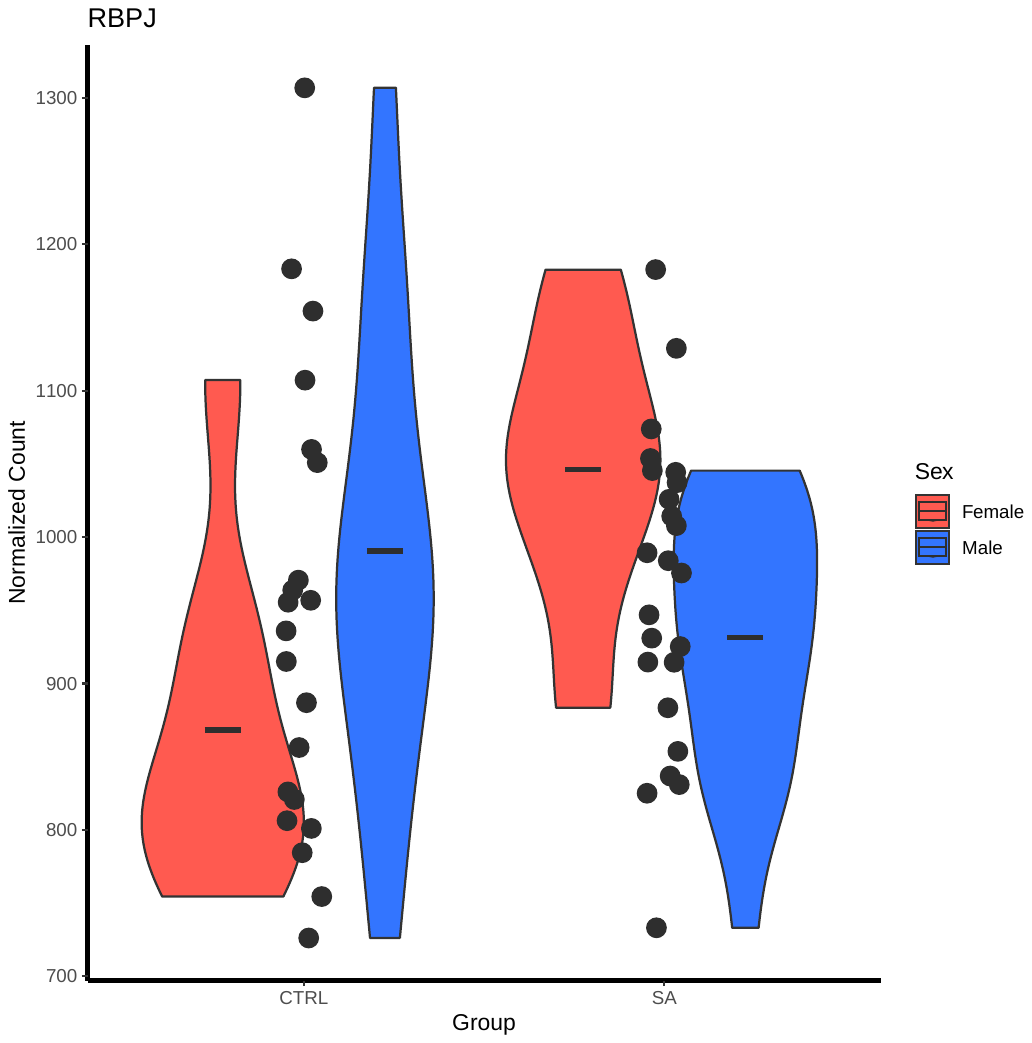  CTRL ELA | 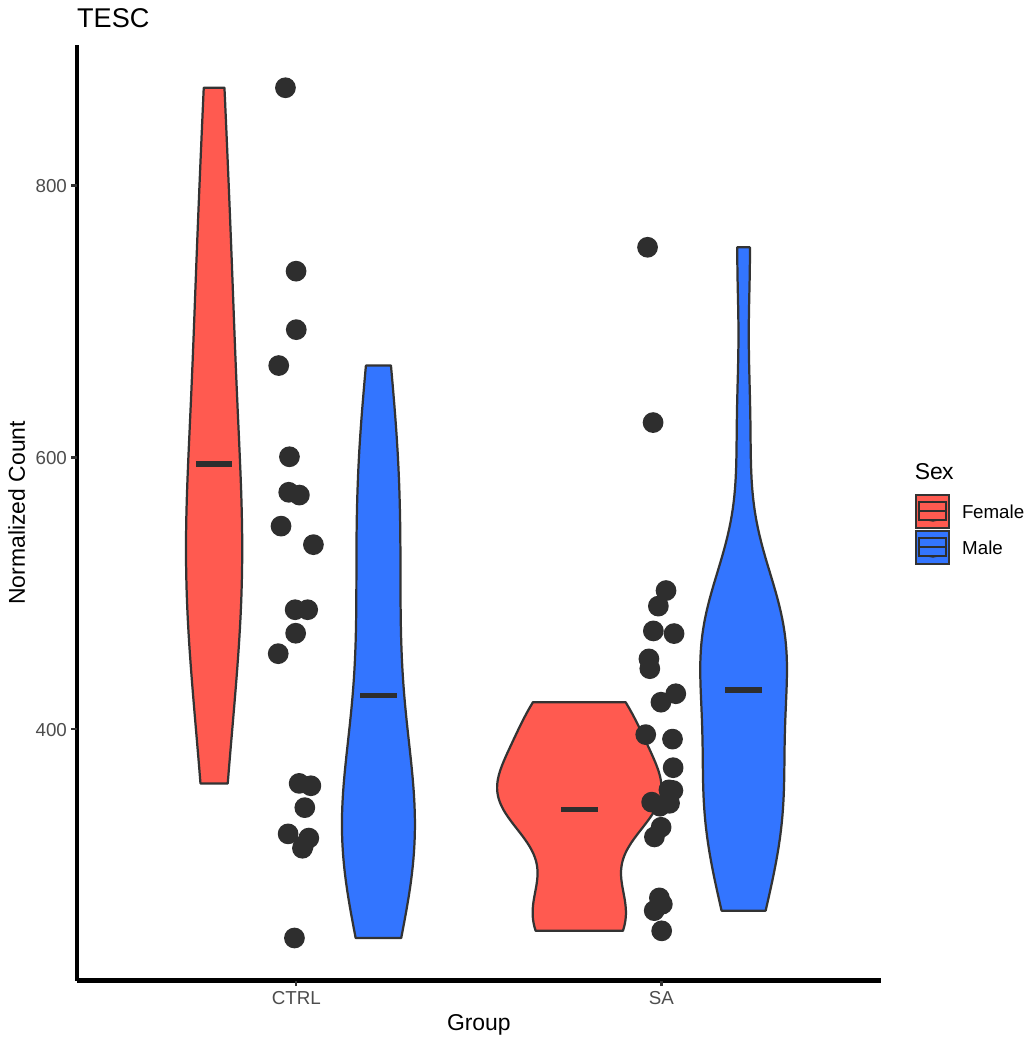  CTRL ELA  CTRL ELA |

b)

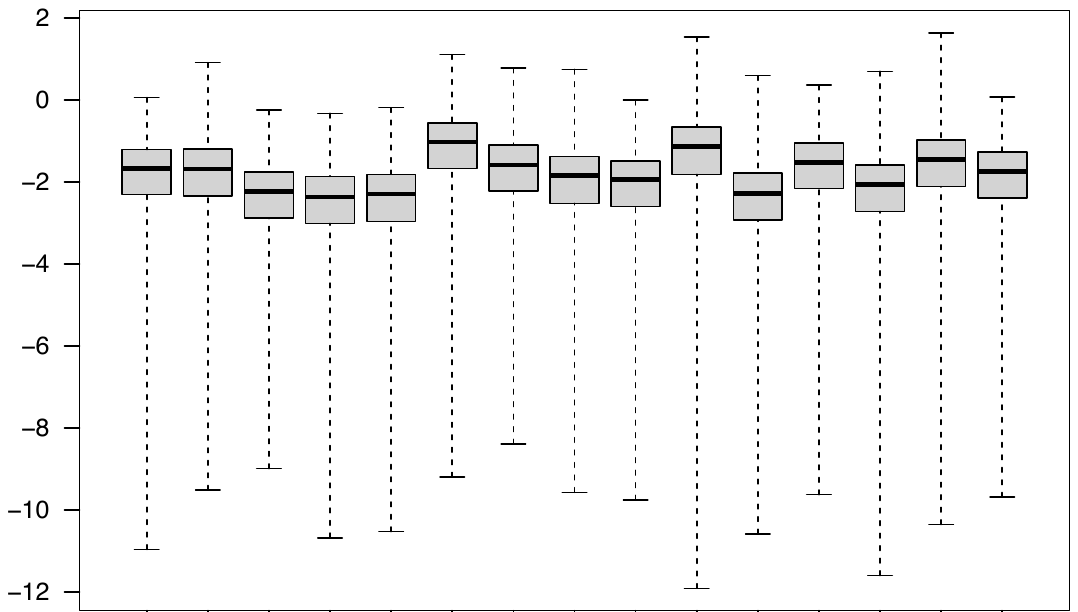

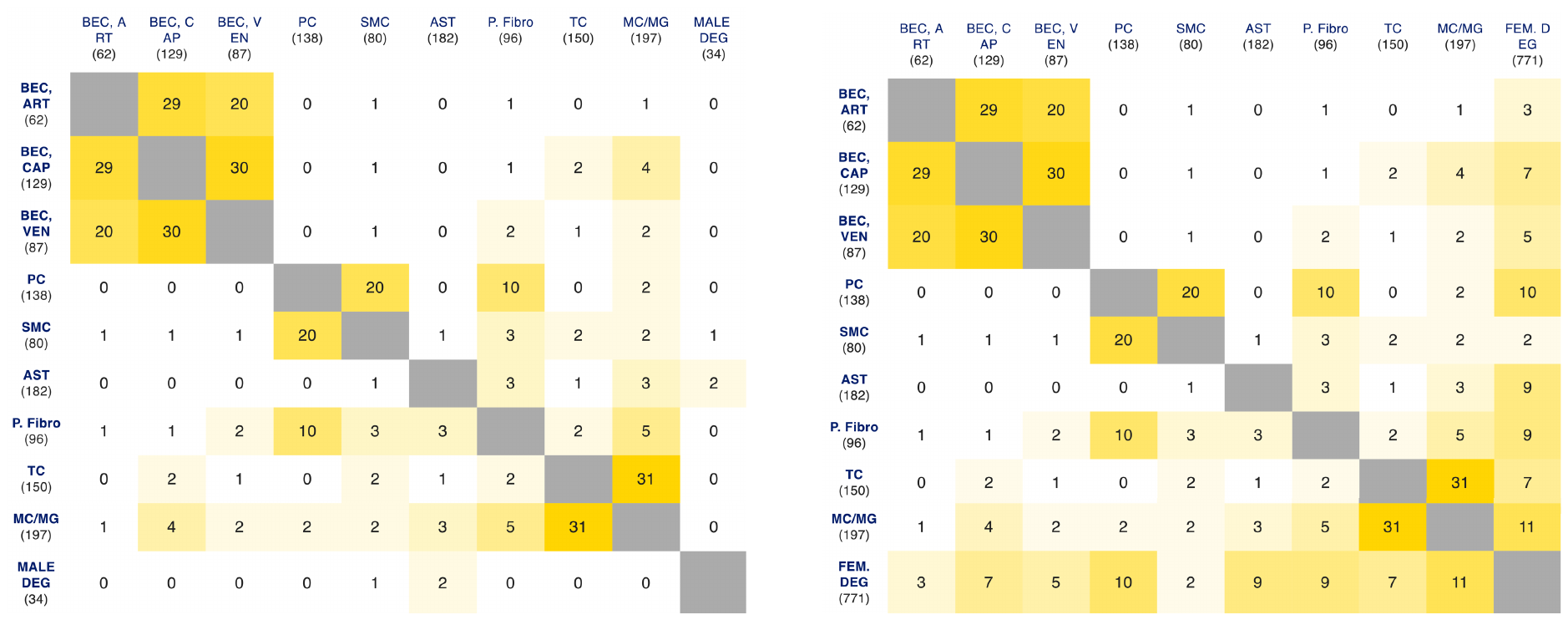
c)

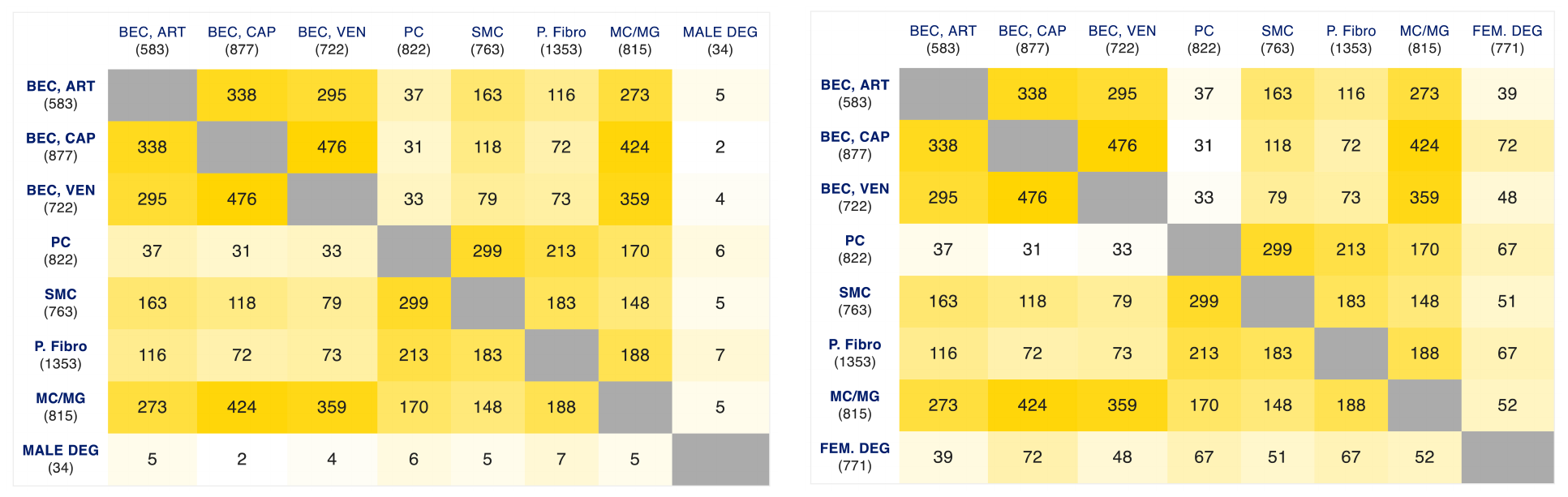
d)

**Supplementary Fig. 4**

Analysis of gene-disease associations for DEGs related to psychiatric disorders by querying the PsyGeNET database. a) Plot showing disease associations for male DEGs. b) Plot showing disease associations for female DEGs. c) Female DEGs were queried using STRING DB to examine whether their corresponding protein products belonged to interacting networks. Output indicates complex interaction between several networks. d-e) Cell-type specific expression of d) HOMER2 and e) TACC1. Average normalized counts and corresponding plots were extracted from the Human_BBB shiny app (<https://twc-stanford.shinyapps.io/human_bbb/>, Yang et al., 2022). HOMER2 is most highly expressed in endothelial cells and perivascular fibroblasts. TACC1 is similarly highly expressed in endothelial, pericyte, and smooth muscle cells.

a)

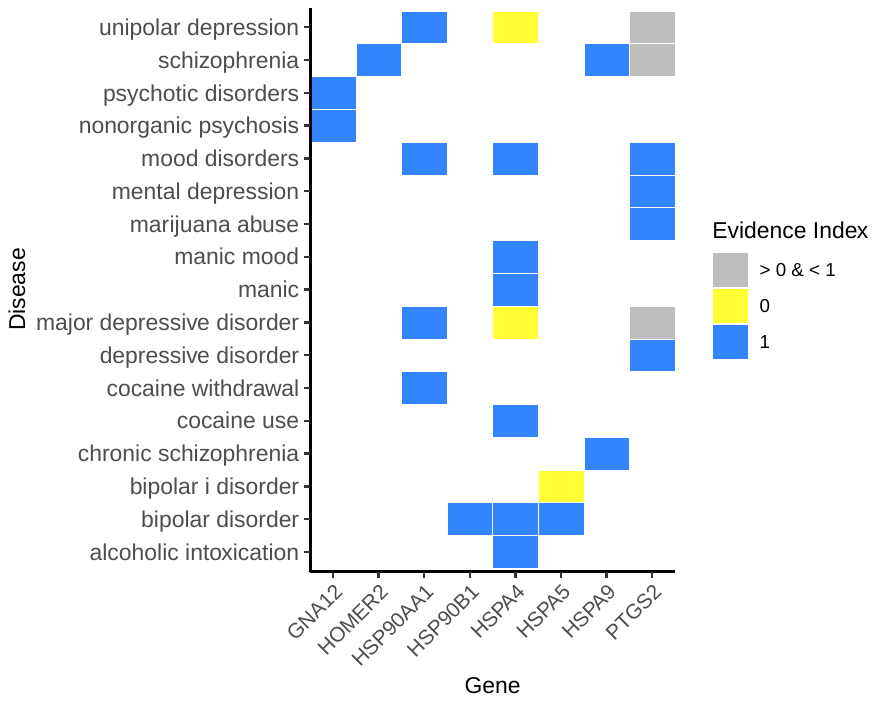

b)

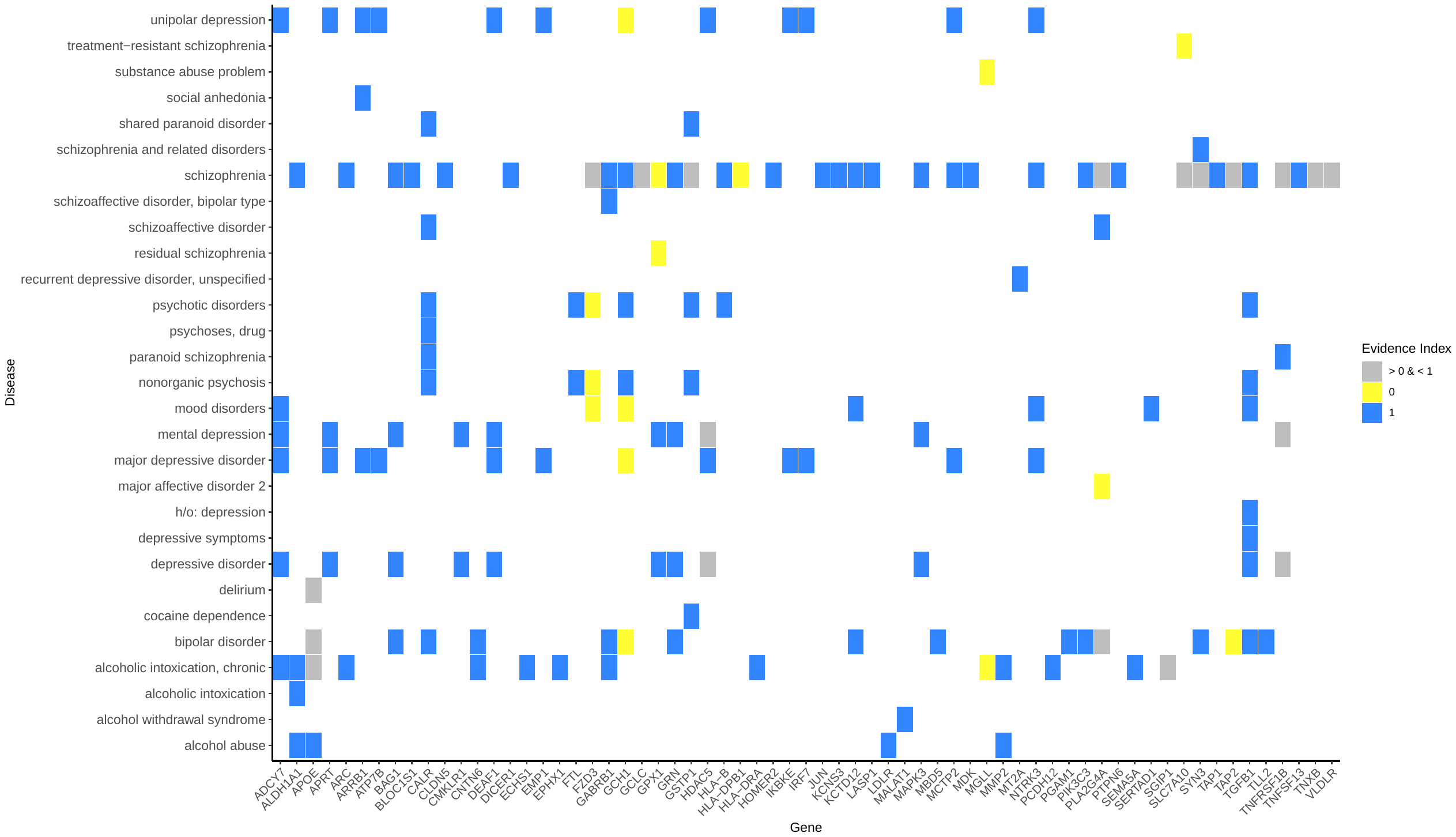

c)

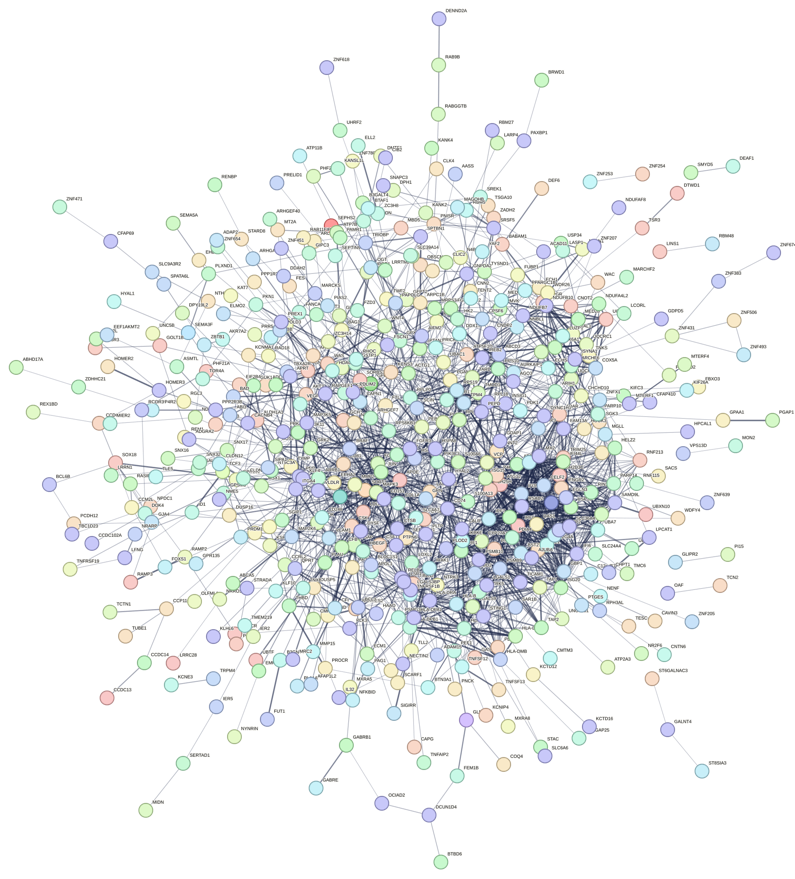

d)

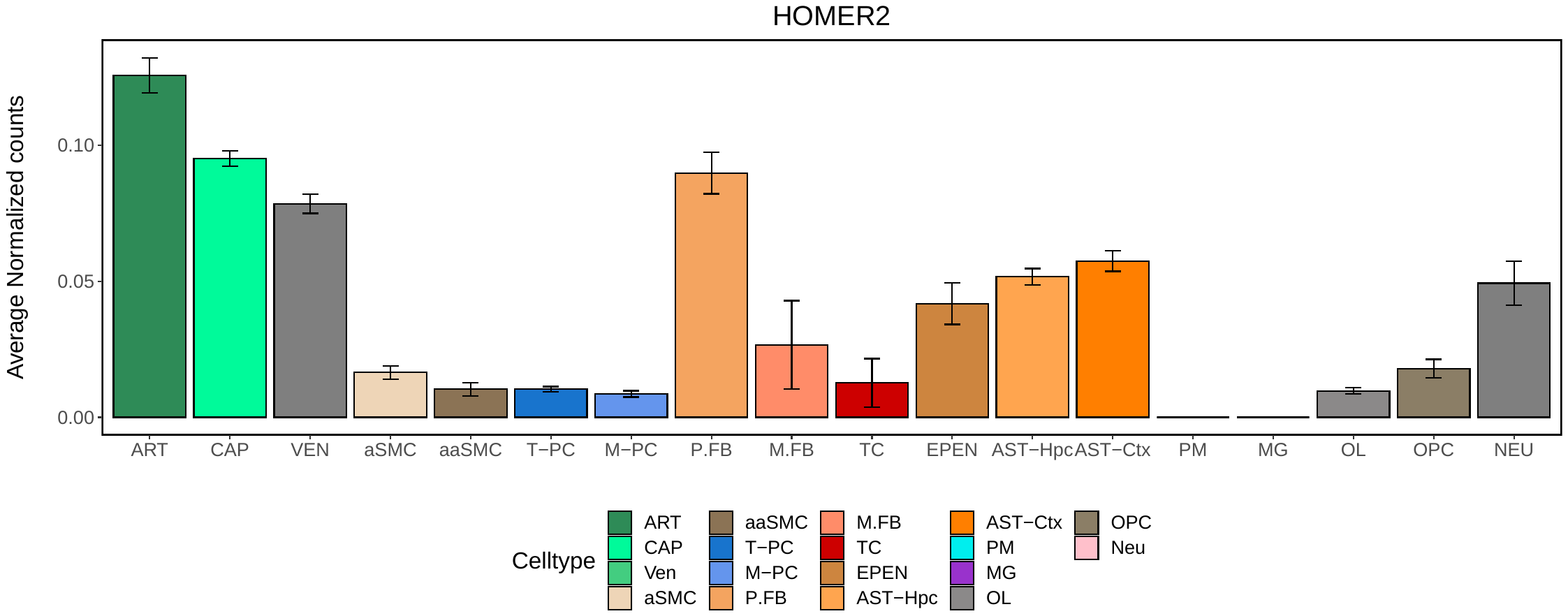
e)

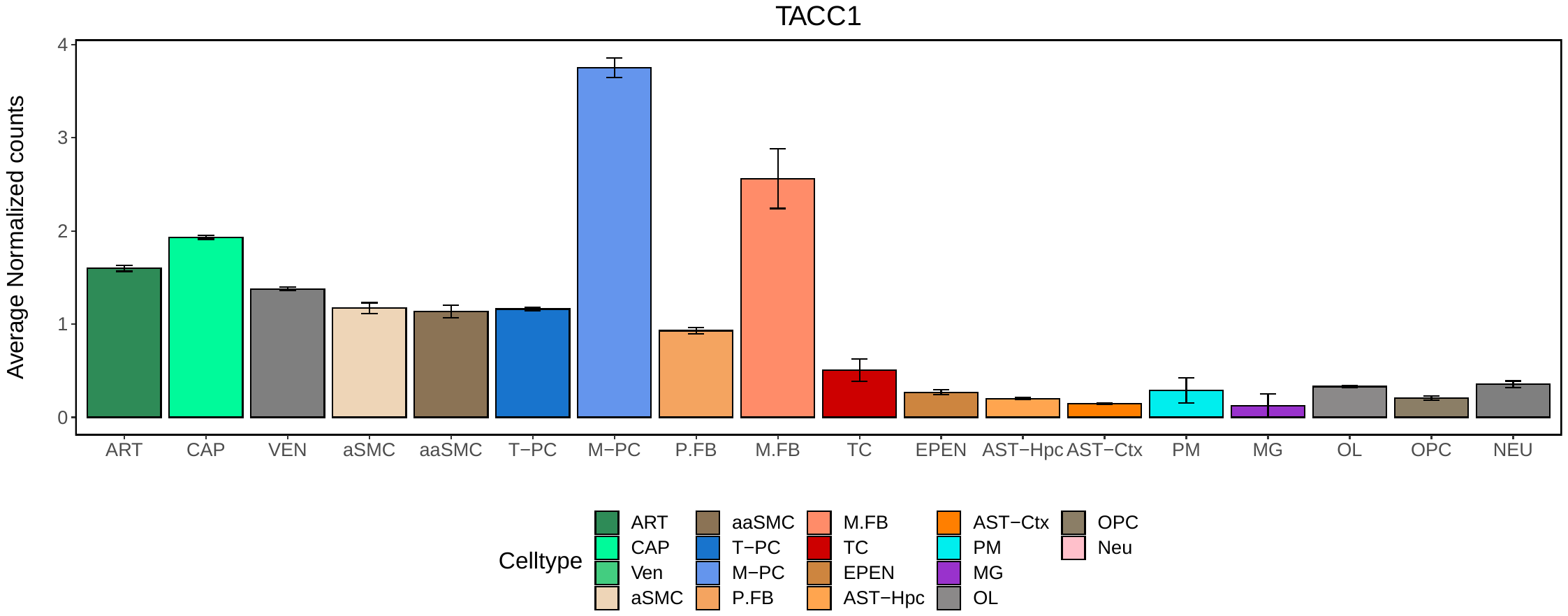

**Supplementary Fig. 5**

a-b) Venn diagrams showing a) overlap between WGCNA module genes and downregulated genes from the RRHO analysis in females; and b) overlap between WGCNA module genes and upregulated genes from the RRHO analysis in females. c) Overlap of genes within the turquoise and black modules with human homologues of genes affected by dual KLF2/4 knockout. This figure indicates substantial overlap, with 448 genes in the turquoise module (p-value = 5.1×10^−9^) and 193 genes in the black module (p-value = 3.3×10^−6^), indicating that the turquoise and black modules possessed a significantly large proportion of genes whose expression is impacted or regulated by KLF2 and KLF4, respectively. d) Raw sequencing counts from Lutz & Tanti et al. (2017) were re-analyzed using the same bioinformatic pipeline presented in this study, including pipelines for low count filtering, covariate identification, and DGE analysis to obtain log2 fold changes for genes known to have immune or vascular functions. A correlation analysis was conducted to assess the relationship between the log2 fold change values for these genes from both datasets. A moderately strong positive correlation was found between log2 fold changes of immune-related and vascular genes within the brain parenchyma and NVU, indicating a similar pattern of downregulation for these genes. e) Transcription factor FOXO1 and histone deacetylase HDAC1 were previously identified as repressors of CLDN5 expression at the NVU (Dudek et al., 2020). KLF2 and KLF4 were queried using STRING DB to examine whether their corresponding protein products interact with FOXO1 and HDAC1. Output indicates that there is indeed interaction between protein products, where HDAC1 and FOXO1 are potential upstream regulators that interact with KLF2 and KLF4 and, in turn, KLF2 and KLF4 directly interact with CLDN5. These interactions may be implicated in the modulation of neurovascular CLDN5 expression in stress-induced pathology. f) Venn diagrams demonstrating (left) overlap between female DEGs (identified from differential gene analysis), turquoise module genes (identified from WGCNA), and KLF2 motif-linked genes (identified from snATAC-seq data); and (right) overlap between female DEGs (identified from differential gene analysis), black module genes (identified from WGCNA), and KLF4 motif-linked genes (identified from snATAC-seq data). Below are the gene lists corresponding to the intersections of the respective Venn diagrams.

a)

| 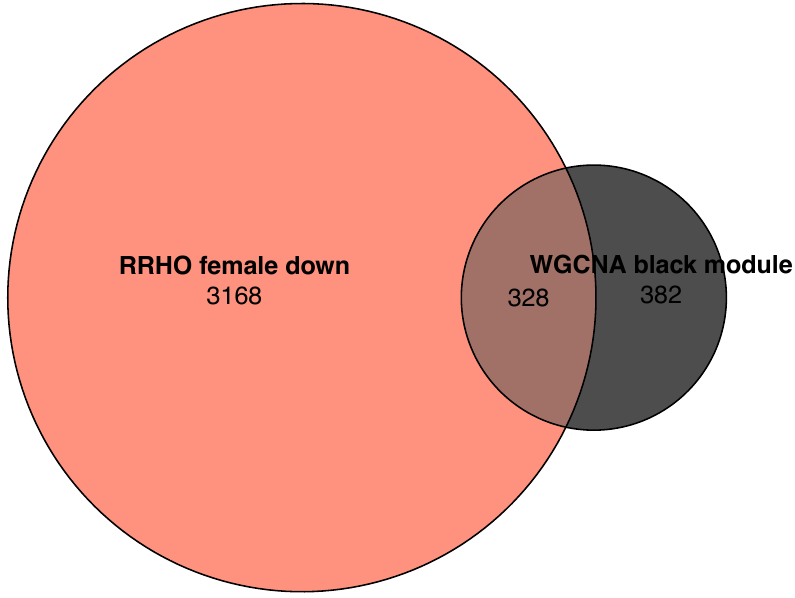 | 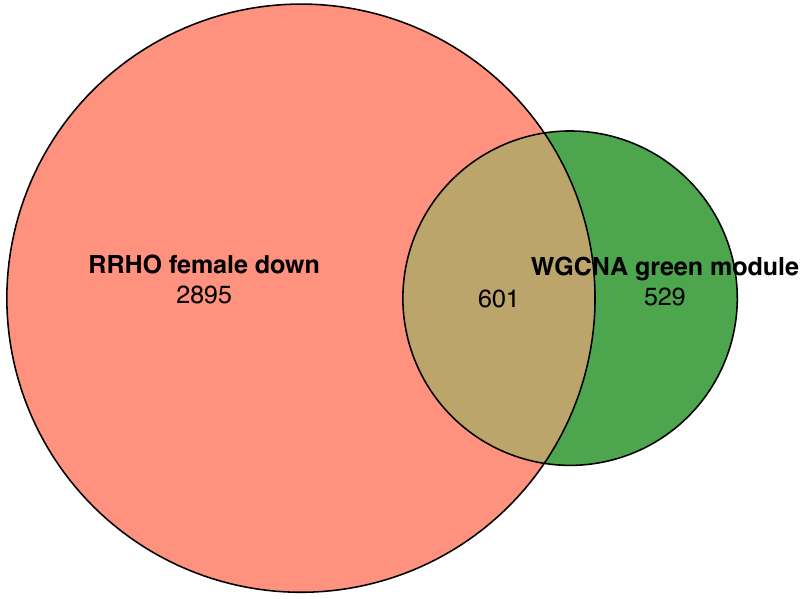 | 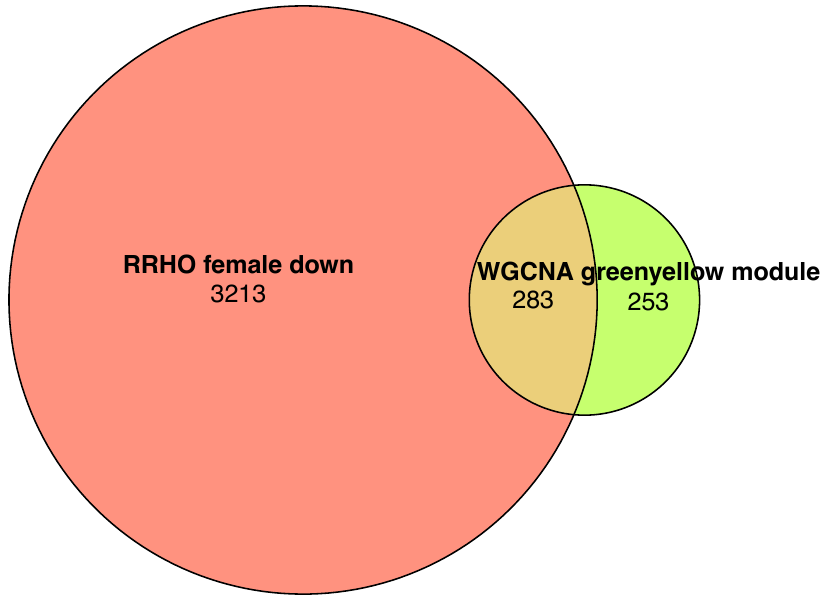 |
| --- | --- | --- |
| 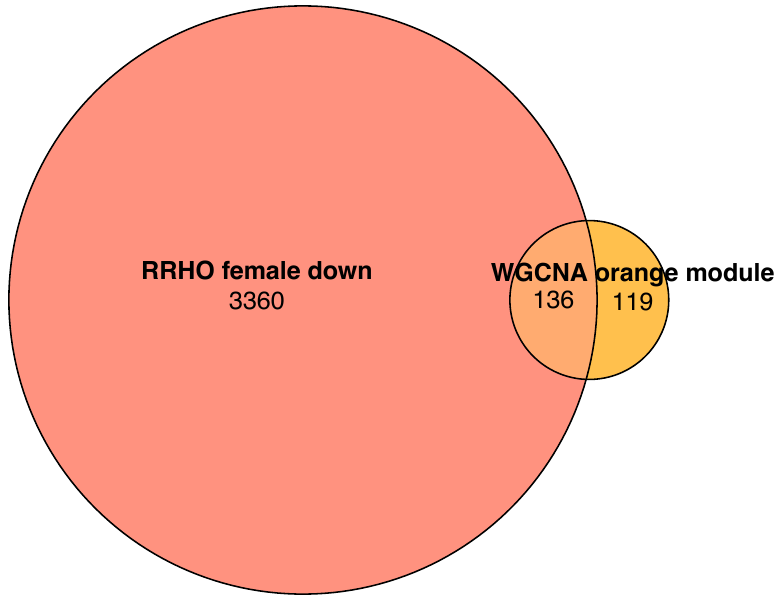 | 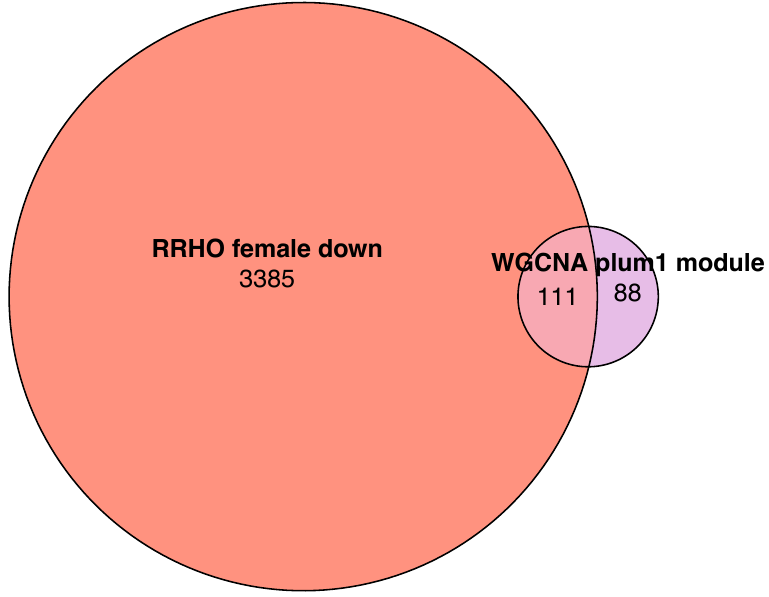 | 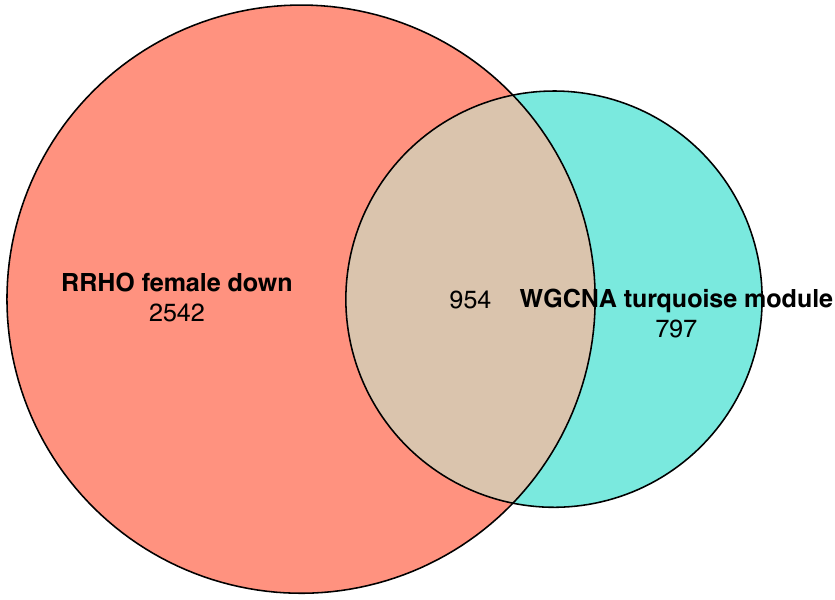 |
| 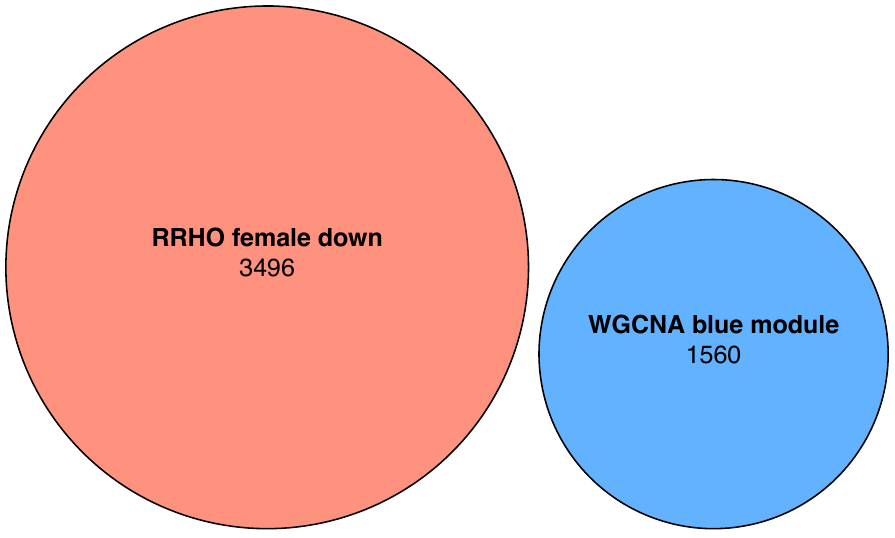 | 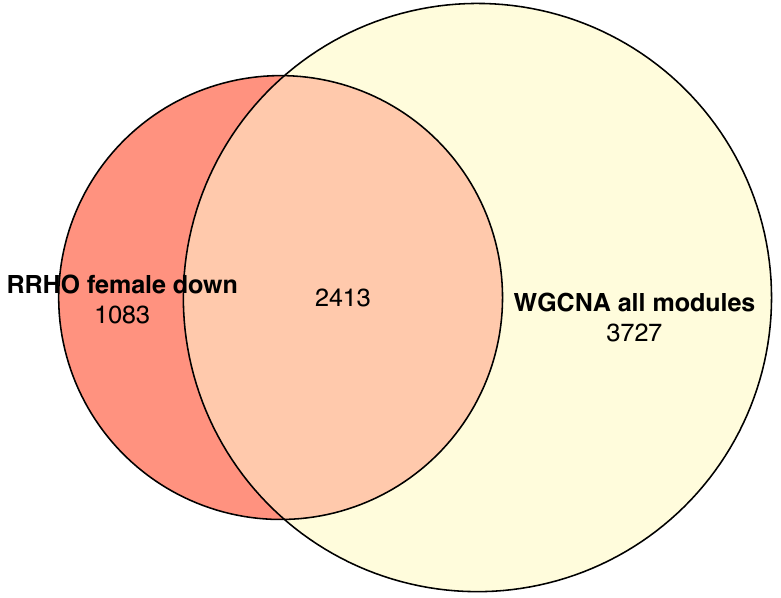 |  |
| b) |  |  |
| 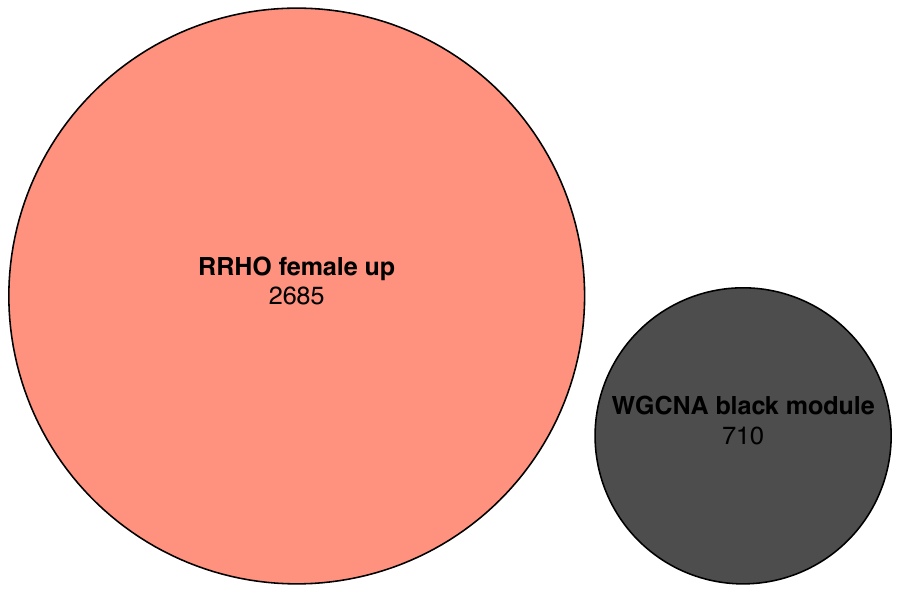 | 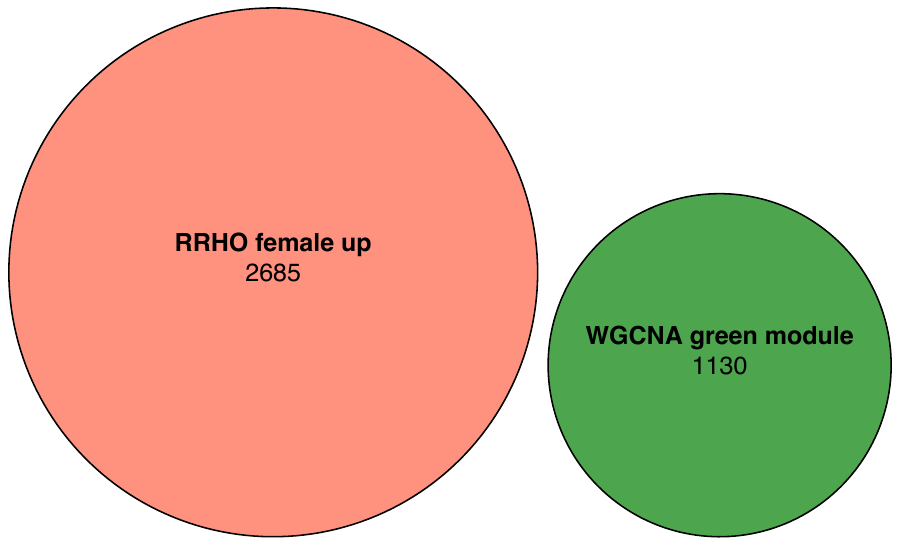 | 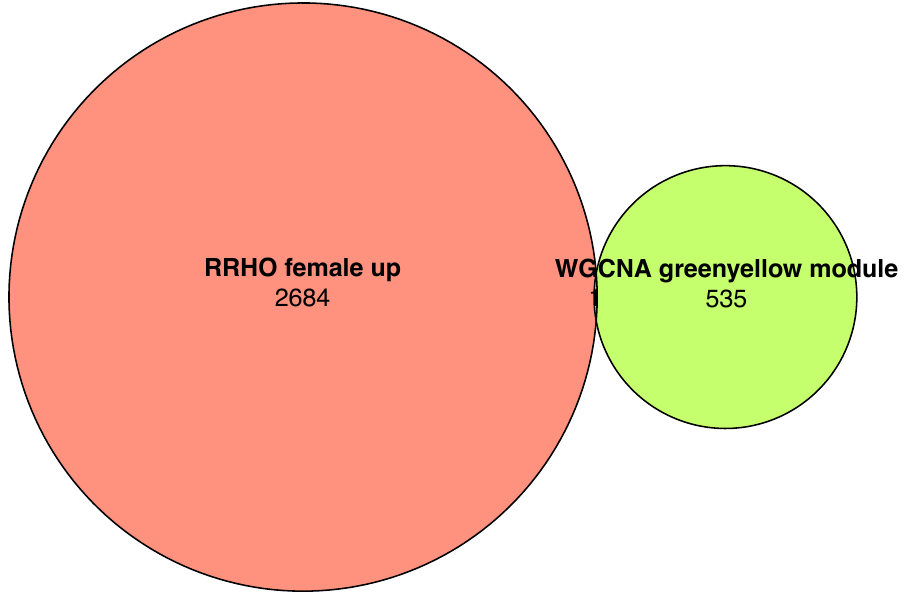 |
| 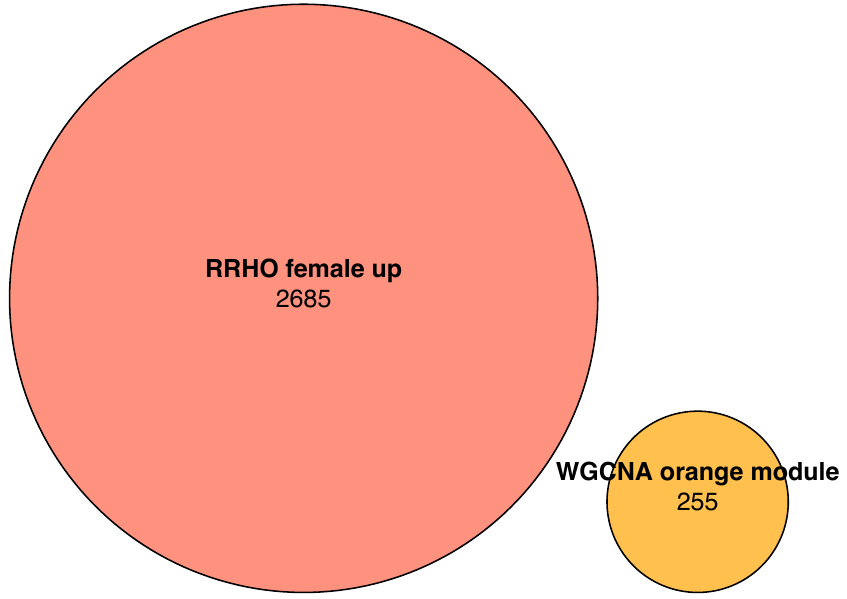 | 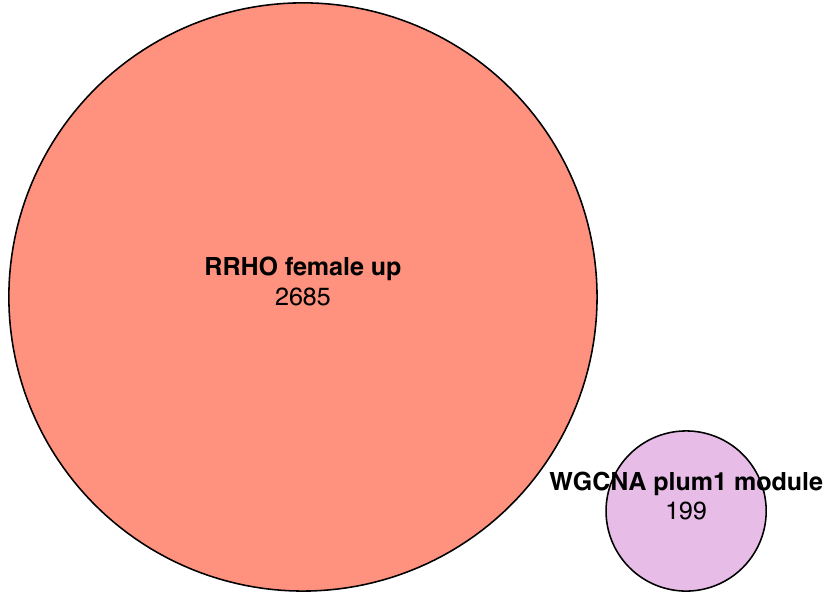 | 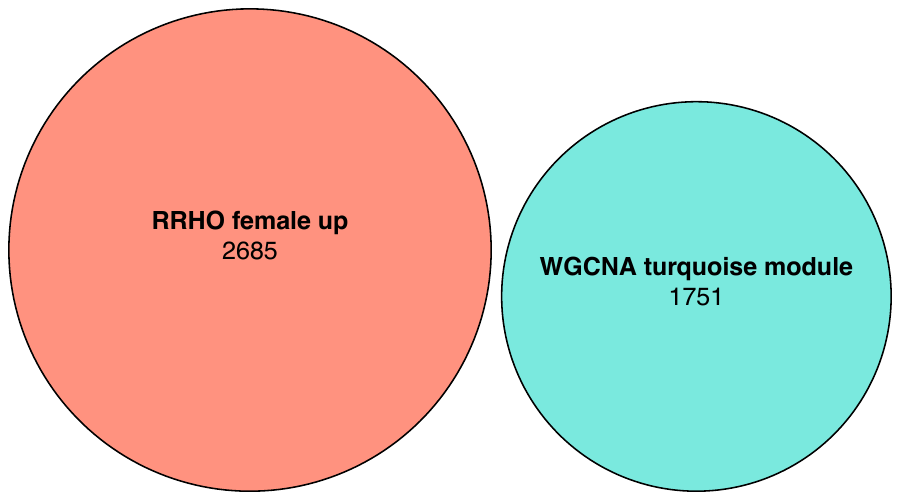 |
| 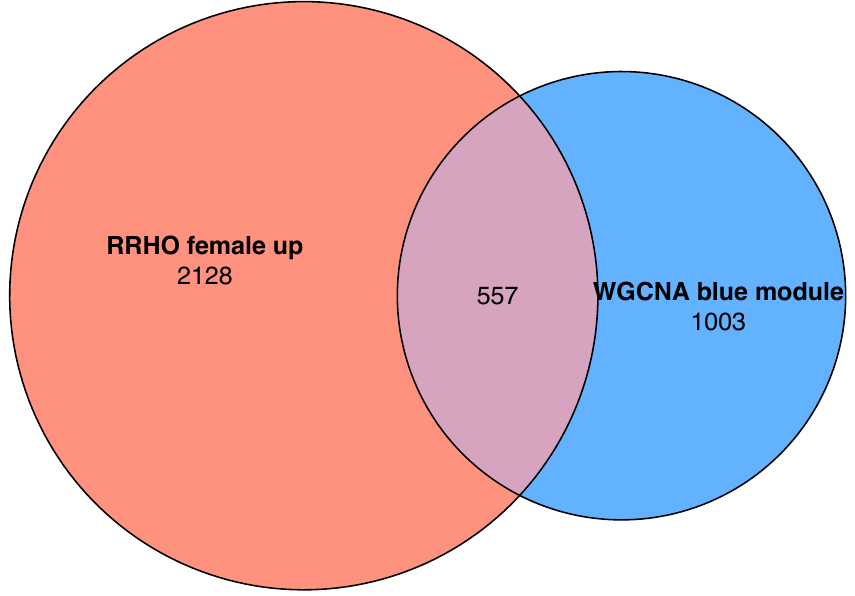 |  | |

c)

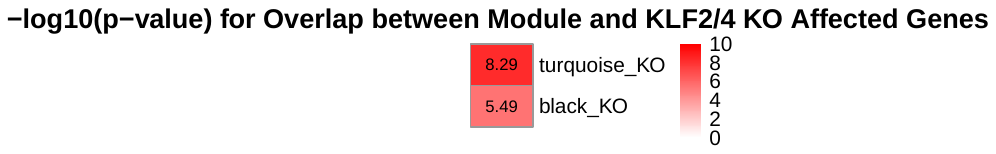

d)

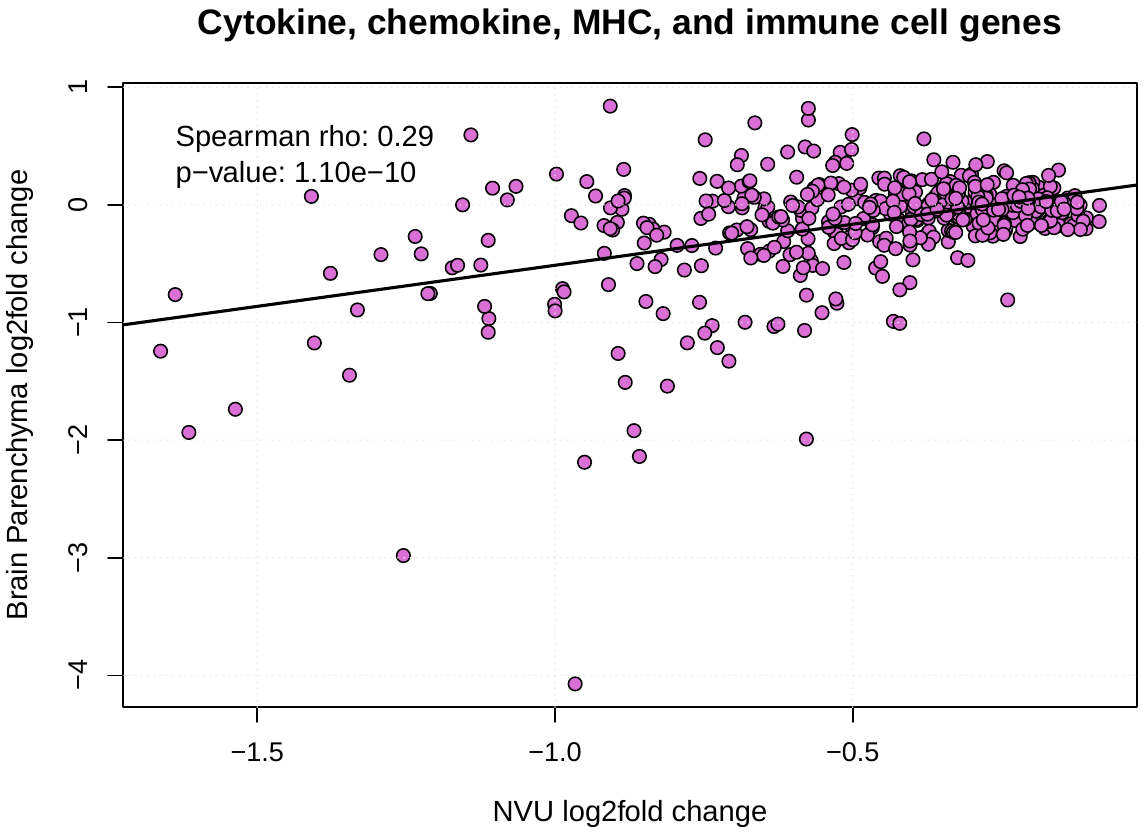

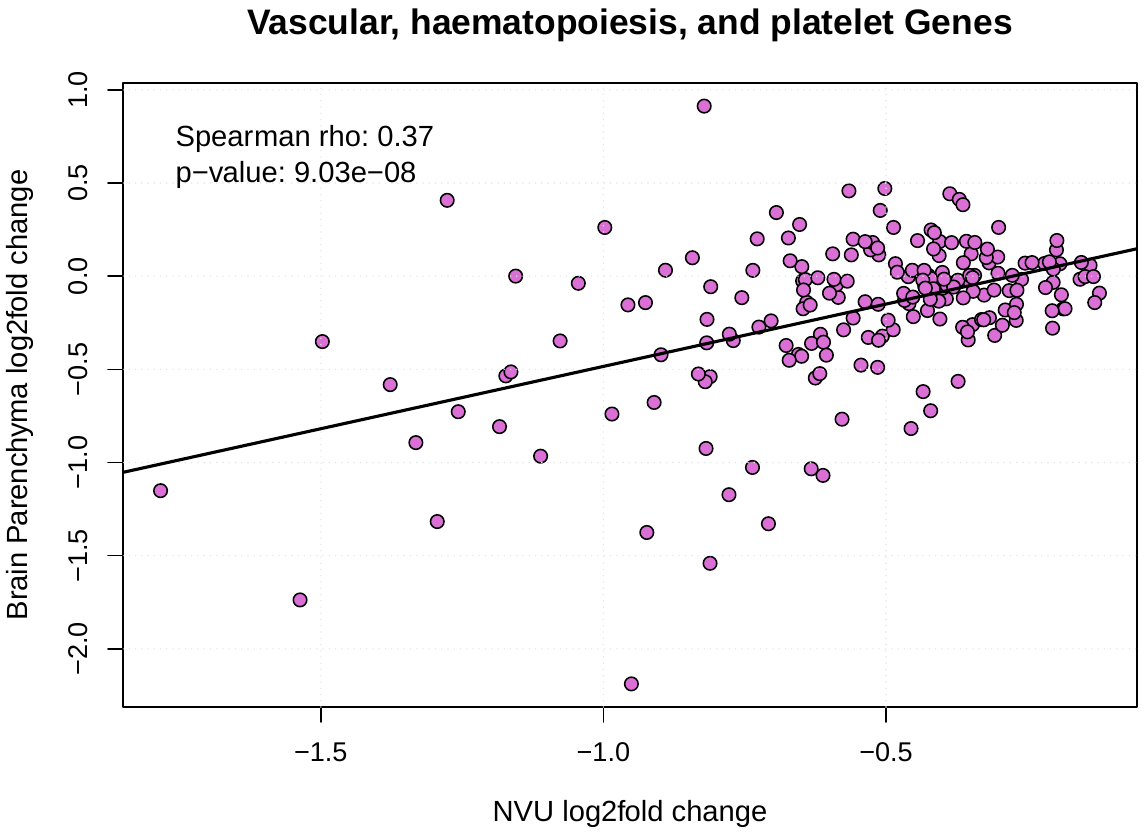

e)

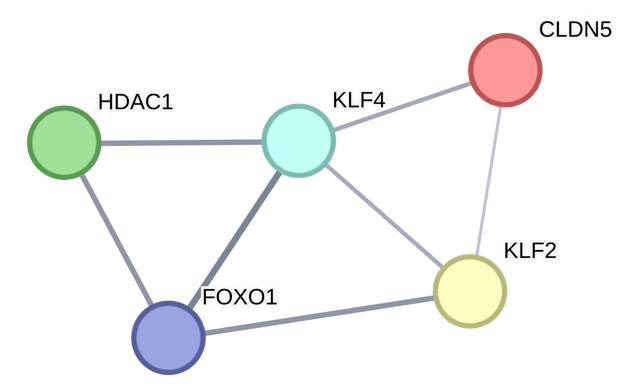

f)

| 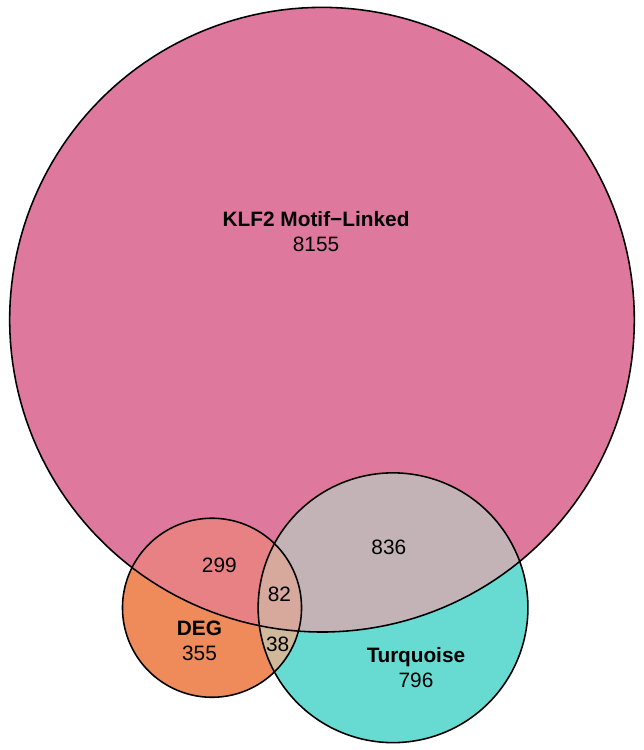 | 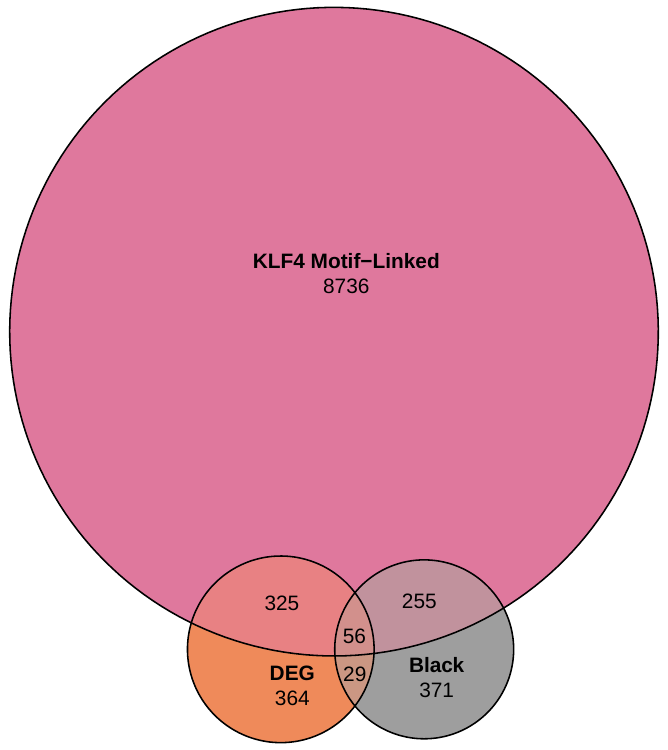 |
| --- | --- |

| **DEG ∩ turquoise module ∩ KLF2 motif-linked** | **DEG ∩ black module ∩ KLF4 motif-linked** |
| --- | --- |
| **ABHD17A** | ADAR |
| **ACTG1** | ARHGAP25 |
| **AKR7A2** | ATP2A3 |
| **AP1B1** | BLOC1S1 |
| **APRT** | BST2 |
| **ARRB1** | BTN3A1 |
| **ATP5F1D** | C12orf57 |
| **AURKAIP1** | CALR |
| **BAALC** | CCRL2 |
| **BABAM1** | CMTM3 |
| **BTBD6** | COX6A1 |
| **CAPN1** | DAP |
| **CD81** | ECSCR |
| **CH25H** | EFCC1 |
| **CHCHD10** | EVA1B |
| **CPNE2** | FOXS1 |
| **CTSD** | FSCN1 |
| **DEAF1** | GLIPR2 |
| **DGCR6L** | GPSM3 |
| **ECH1** | GSTP1 |
| **ECM1** | IFI35 |
| **ETV3** | IFIT3 |
| **FAM89B** | IRF9 |
| **FOXF1** | ISG15 |
| **GAMT** | KCNE3 |
| **GNB2** | KIF26A |
| **GPAA1** | KLF4 |
| **GRK2** | LGALS1 |
| **GUK1** | LUZP1 |
| **HCFC1R1** | MAGED2 |
| **HDAC5** | MXRA5 |
| **HOMER3** | NRROS |
| **HPCAL1** | OLFML1 |
| **HSPA8** | OLFML3 |
| **ID1** | PARP14 |
| **IFI6** | PLAC9 |
| **KIFC3** | PRICKLE1 |
| **KLF2** | PSMB10 |
| **MMP15** | PSMB8 |
| **MMP28** | PSMB9 |
| **MRC2** | PTGES |
| **MRPS34** | PTPN6 |
| **MT2A** | RPH3AL |
| **MXRA8** | SEPHS2 |
| **NDUFAF8** | SPTBN1 |
| **NHSL1** | TAP1 |
| **NR2F6** | TGFB1 |
| **NTHL1** | TMEM173 |
| **NUTF2** | TNFRSF19 |
| **OAF** | TNFRSF1B |
| **OAS1** | TOR4A |
| **OBSCN** | UNC93B1 |
| **PAMR1** | VOPP1 |
| **PDLIM2** | WDFY4 |
| **PEBP1** | ZNF205 |
| **PEF1** | ZNFX1 |
| **PGAM1** |  |
| **PGLS** |  |
| **PLEC** |  |
| **PLTP** |  |
| **PMVK** |  |
| **PPP1R14B** |  |
| **PRELID1** |  |
| **PREX1** |  |
| **PRR5** |  |
| **PSAP** |  |
| **PSMG3** |  |
| **PYGM** |  |
| **R3HCC1** |  |
| **RNF213** |  |
| **RPLP0** |  |
| **SAMD1** |  |
| **SELENON** |  |
| **SPATA20** |  |
| **STAC** |  |
| **TMEM219** |  |
| **TSR3** |  |
| **TYMP** |  |
| **UBTF** |  |
| **UQCRC1** |  |
| **VSTM4** |  |
| **ZNF428** |  |
